# A modular two yeast species secretion system for the production and preparative application of fungal peroxygenases

**DOI:** 10.1101/2020.07.22.216432

**Authors:** Pascal Püllmann, Anja Knorrscheidt, Judith Münch, Paul R. Palme, Wolfgang Hoehenwarter, Sylvestre Marillonnet, Miguel Alcalde, Bernhard Westermann, Martin J. Weissenborn

## Abstract

Fungal unspecific peroxygenases (UPOs) are biocatalysts of outstanding interest. Providing access to novel UPOs using a modular secretion system was the central goal of this work. UPOs represent an enzyme class, catalysing versatile oxyfunctionalisation reactions on a broad substrate scope. They are occurring as secreted, glycosylated proteins bearing a haem-thiolate active site and solely rely on hydrogen peroxide as the oxygen source. Fungal peroxygenases are widespread throughout the fungal kingdom and hence a huge variety of UPO gene sequences is available. However, the heterologous production of UPOs in a fast-growing organism suitable for high throughput screening has only succeeded once—enabled by an intensive directed evolution campaign. Here, we developed and applied a modular Golden Gate-based secretion system, allowing the first yeast production of four active UPOs, their one-step purification and application in an enantioselective conversion on a preparative scale. The Golden Gate setup was designed to be broadly applicable and consists of the three module types: i) a signal peptide panel guiding secretion, ii) UPO genes, and iii) protein tags for purification and split-GFP detection. We show that optimal signal peptides could be selected for successful UPO secretion by combinatorial testing of 17 signal peptides for each UPO gene. The modular episomal system is suitable for use in *Saccharomyces cerevisiae* and was transferred to episomal and chromosomally integrated expression cassettes in *Pichia pastoris*. Shake flask productions in *Pichia pastoris* yielded up to 24 mg/L secreted UPO enzyme, which was employed for the preparative scale conversion of a phenethylamine derivative reaching 98.6 % *ee*. Our results demonstrate a rapid workflow from putative UPO gene to preparative scale enantioselective biotransformations.

## Introduction

Fungal unspecific peroxygenases (UPOs) have recently emerged as novel hydroxylation biocatalysts. They solely rely on hydrogen peroxide as cosubstrate reaching impressive total turnover numbers on sp^3^-carbon hydroxylations of up to 300,000^1–4^. There is an estimated number of more than 4000 putative UPO genes currently annotated and widely spread within the fungal kingdom representing just a small fraction of the available genetic diversity^5^. To provide further insight into the natural function of UPOs as well as broadening the available substrate scope, it is crucial to access more enzymes from diverse phylogenetic backgrounds. It would be desirable to heterologously produce these enzymes utilising fast-growing standard laboratory hosts such as bacteria or yeast. These organisms would facilitate protein engineering and allow directed evolution campaigns for tailoring desirable traits. Although substantial work has been invested into the heterologous expression of the first discovered *Agrocybe aegerita* UPO (*Aae*UPO) using the yeast *Saccharomyces cerevisiae*, sufficient protein amounts of 8 mg/L were obtained as the result of an intensive directed evolution campaign^6^. This fundamental work led to several successful UPO studies on a range of substrates from agrochemicals to pharmaceuticals^7–10^.

The successful production was achieved by the introduction of nine amino acid exchanges. Four of these were localised within the 43 amino acid signal peptide, which orchestrates protein secretion in the natural fungal host as well as in *S. cerevisiae*. The engineered signal peptide combined with the wildtype *Aae*UPO enzyme resulted in a 27-fold increase in protein secretion yield highlighting the paramount importance of the signal peptide for heterologous production as already shown by others^11–16^. Recent studies reported the production of UPOs in *E. coli*^17,18^. However, it remains elusive whether these recombinant peroxygenases harbour comparable activities and stabilities to UPOs produced in eukaryotic hosts. The reported expression yields are substantially lower when compared to *S. cerevisiae* raising the question, whether enough functional protein could be produced for laboratory evolution campaigns.

Golden Gate cloning has proven to be an invaluable synthetic biology tool enabling seamless assembly of gene fragments utilising type IIs restriction enzymes^19–27^. It can be performed in an affordable and straightforward one-pot, one-step digestion-ligation setup with efficiencies of correct assembly close to 100 %. Golden Gate cloning utilises type IIs restriction enzymes like *Bsa*I or *Bbs*I, which are characterised by their ability to cut outside of their respective recognition sequence and leads to the creation of defined 4 base pair sticky overhangs. These overhangs can be specified by PCR, allowing a sequence defined, efficient and seamless assembly of nine and more gene fragments in a one-pot and one-step manner^20,27,28^.

For the detection of the target protein secretion in small volumetric amounts of yeast supernatant, a sensitive, high-throughput suitable, and protein-specific assay would be highly beneficial. Previously reported split-GFP (green fluorescent protein) systems, which rely on tagging the protein of interest with a short 15 amino acid peptide tag and subsequent GFP reconstitution, present an ideal tool for this task^29,30^. Superfolder GFP (sfGFP) possesses a classical β-barrel protein fold, built up out of eleven β-sheets. By genetic removal of one *C*-terminal beta-sheet (GFP11), the fluorescence signal of the remaining protein (GFP1-10) is quenched. Using the GFP11 peptide as protein-tag hence allows the specific measurement of target protein amounts by fluorescence using GFP1-10 as detector fragment, since upon interaction of both parts, the fluorescence of sfGFP is restored.

In this study, we envisioned a tripartite Golden Gate-based modular system. This system consists of the modules ‘signal peptide’, ‘UPO gene’ and ‘protein-tag’ (Fig. 1a). The ‘protein-tag’ module combines the affinity-based purification as well as the enzyme quantification by split-GFP. This *S. cerevisiae* expression system gave rise to a rapid workflow from UPO gene to heterologously produced and purified UPOs.

**Fig. 1.**
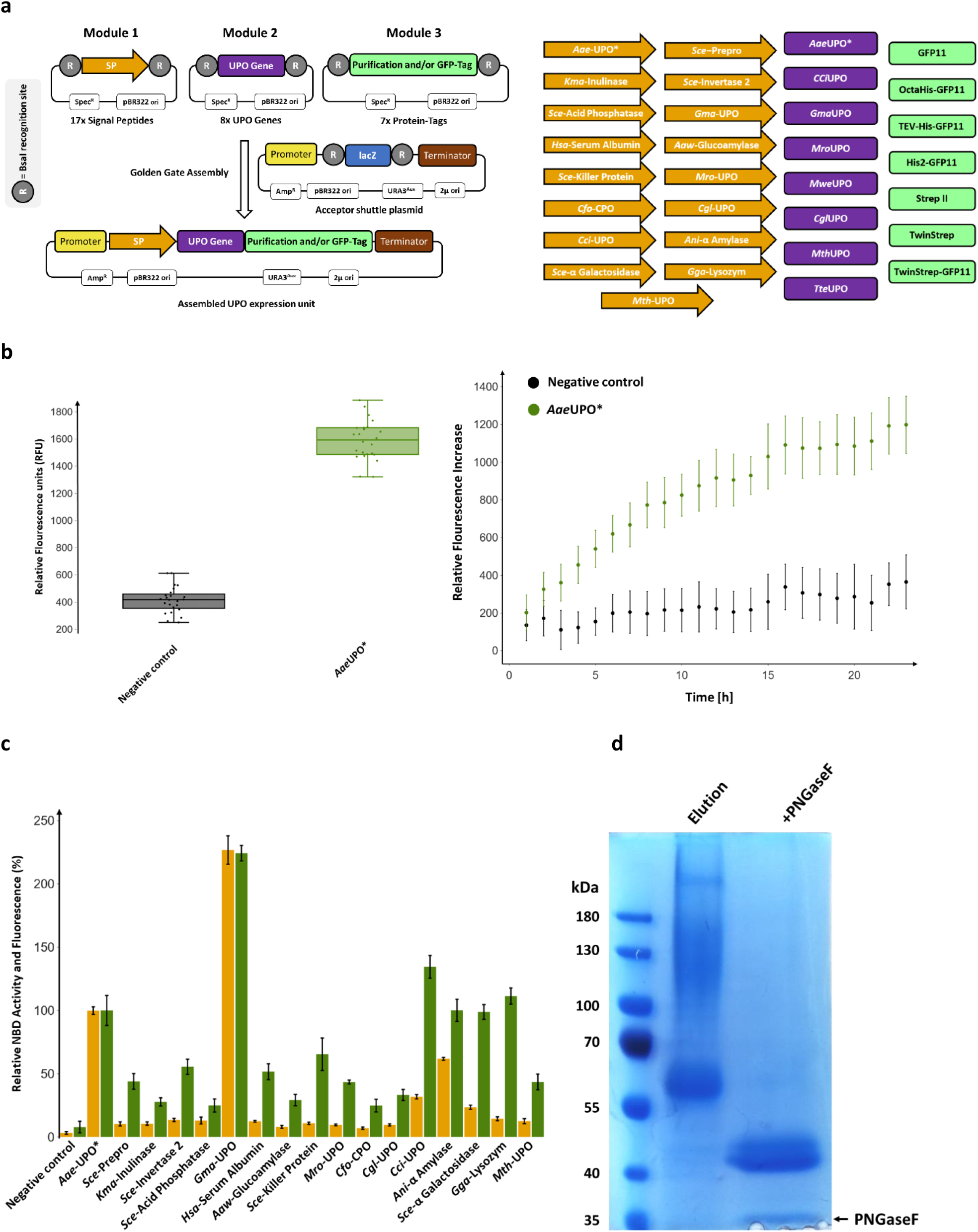
The Golden Gate system consisting of the modules signal peptide, UPO gene and protein-tag and its functional verification regarding split-GFP assay, signal peptide shuffling and purification using the model UPO *Aae*UPO* in *S. cerevisiae*. **a** *Left*: Concept of the modular Golden Gate system as a tripartite system, consisting of signal peptide (SP; contains ATG start codon), UPOgene (lacking start and stop codon) and *C*-terminal Tag (contains stop codon). *Right*: Overview of the individual parts of the modular shuffling systems, containing 17 signal peptides, 8 UPO genes and 7 *C*-terminal protein tags. Detailed sequence information of all parts can be found in Supplementary Table 2 and Supplementary Table 3. **b** Quantification of the UPO secretion in *S. cerevisiae* using the split-GFP system. Two constructs were utilised for testing, namely a previously derived yeast secretion variant of *Aae*UPO (*Aae*UPO*) and further including a *C*-terminal GFP11. The acceptor shuttle plasmid (pAGT572_Nemo 2.0) was used as negative control. *Left:* biological replicates (n=24) of *Aae*UPO* and the negative control were screened within the split-GFP assay. Relative fluorescence units (RFU) were measured at 0 and 72 h after adding GFP1-10. The values are shown as boxplots (*Aae*UPO*: median = 1589, s.d. 8.9 %; negative control: median = 416, s.d. 22.6 %) with individual data points shown as dots. *Right:* Continuous fluorescence measurements (24 hours; 23 time points) of each construct. Data are mean of fluorescence - background (background = first measurement after 1 hour) ± s.d. of biological replicates (n=24). **c** Screening of the constructed signal peptide shuffling library utilising *Aae*UPO* as reference protein. Values for 5-nitro-1,3-benzodioxole (NBD) conversion (orange bars) and fluorescence by split-GFP assay (green bars) were normalised to the previously used *Aae*UPO SP* -*Aae*UPO* construct (100 %). Data are mean ± s.d. of biological replicates (n=5). Primary data are displayed in Supplementary Table 8. Detailed information on the origin and the sequence of the signal peptides can be found in Supplementary Table 2. **d** SDS-PAGE of the purity of *Aae*UPO* after one step TwinStrep tag^®^ purification, utilising the designed TwinStrep-GFP11 purification/detection combination tag. Additionally, *Aae*UPO* was subjected to enzymatic deglycosylation by PNGaseF and analysed (right lane).

To give access to higher protein amounts, we designed two fully compatible episomal and one integrative plasmid for UPO production in the methylotrophic yeast *Pichia pastoris* (syn. *Komagataella phaffii*). In total, four active UPOs were heterologously produced in yeast for the first time. The obtained UPO yields using *P. pastoris* enabled the enantioselective hydroxylation of a phenethylamine derivative on a preparative scale.

## Results

### The modular Golden Gate UPO expression system

Three modules were designed for pre-defined assembly into an episomal *S. cerevisiae* shuttle expression plasmid. We created 32 modules consisting of 17 signal peptides (Module 1), 8 UPO genes (Module 2) and 7 protein-tags (Module 3, Fig. 1a). Module 3 is employed for affinity-based enzyme purification and/or split-GFP-based protein quantification. To verify the envisioned system for protein quantification, the *C*-terminal GFP11 detection tag (Module 3) was assembled with the previously evolved UPO signal peptide *Aae-*UPO* (Module 1) and the engineered peroxygenase *Aae*UPO* (Module 2)^6,31^. The successful split-GFP assay was validated by a significant fluorescence response in the sample with the secreted protein (Fig. 1b). Module 1, exhibiting 17 distinct signal peptides (SP), is the pivotal part for guiding the protein secretion. The signal peptide library consists of sequences originating from *S. cerevisiae*, further yeast organisms, basidiomycetes, ascomycetes and animals (Supplementary Table 2). Seven signal peptide sequences originate from (putative) UPOs or a closely related chloroperoxidase (*Cfu*CPO). To demonstrate the importance of the signal peptide, we assembled the *Aae*UPO* gene (Module 2) and the GFP11 tag (Module 3) with each of the 17 signal peptides (Module 1). UPO secretion levels were monitored by enzymatic activity using the 5-nitro-1,3-benzodioxole (NBD)^32^ assay as well as split-GFP detection (Fig. 1c).

All constructs showed significant secretion levels and enzymatic activities. The signal peptides *Cci*-UPO, *Ani*-α Amylase, *Sce*-α Galactosidase and *Gga*-Lysozyme led to similar protein concentrations as the evolved signal peptide *Aae-*UPO*. The signal peptide *Gma*-UPO resulted in a more than doubled activity and secretion of the *Aae*UPO* enzyme relative to the evolved *Aae*-UPO* signal peptide (220 % increase). This observation is particularly impressive considering that the signal peptide *Aae*-UPO* was evolved for the optimised secretion of *Aae*UPO* in *S. cerevisiae* by subjecting it to several rounds of directed evolution^6^. The signal peptide *Gma*-UPO originates from the putative *Galerina marginata* UPO (*Gma*UPO). When correlating normalised enzymatic activity and split-GFP-based fluorescence values of the signal peptide library, in most cases, higher fluorescence levels were measured than activity values. This observation indicates the occurrence of differing *Aae*UPO* enzyme variants depending on respective signal peptide cleavage. This could be due to the great diversity of the utilised signal peptides likely resulting in differing *N*-termini and affecting the enzymatic activity of the processed enzyme.

To give rise to a general, one-step protein purification protocol for UPOs, Module 3 was further extended to allow for simultaneous affinity-based protein purification and GFP11 based fluorescence detection. Several versions of the GFP11 tag in combination with Strep^®^- or Hexa/Octahistidine-affinity tags were generated and tested (Supplementary Table 3)^33,34^. We used the protein tags with the previously identified combination of signal peptide (*Gma*-UPO, Module 1) and UPO (*Aae*UPO*, Module 2) and identified the TwinStrep-GFP11 protein tag. This tag consists of a double 8 amino acid Strep II tag (Twin-Strep^®^-tag)^35^ and a *C*-terminal GFP11 sequence. Comparison of the modules GFP11 and TwinStrep-GFP11 revealed unaltered enzymatic activities but a significantly higher fluorescence response for the TwinStrep-GFP11 construct (Supplementary Fig. 2). This difference is probably due to better accessibility of the terminal GFP11 portion since the overall size of the tag is increased(27 vs 59 amino acids), and several flexible linkers are included. SDS PAGE analysis revealed the successful one-step purification of the mature protein *Aae*UPO* (Fig. 1d).

**Fig. 2.**
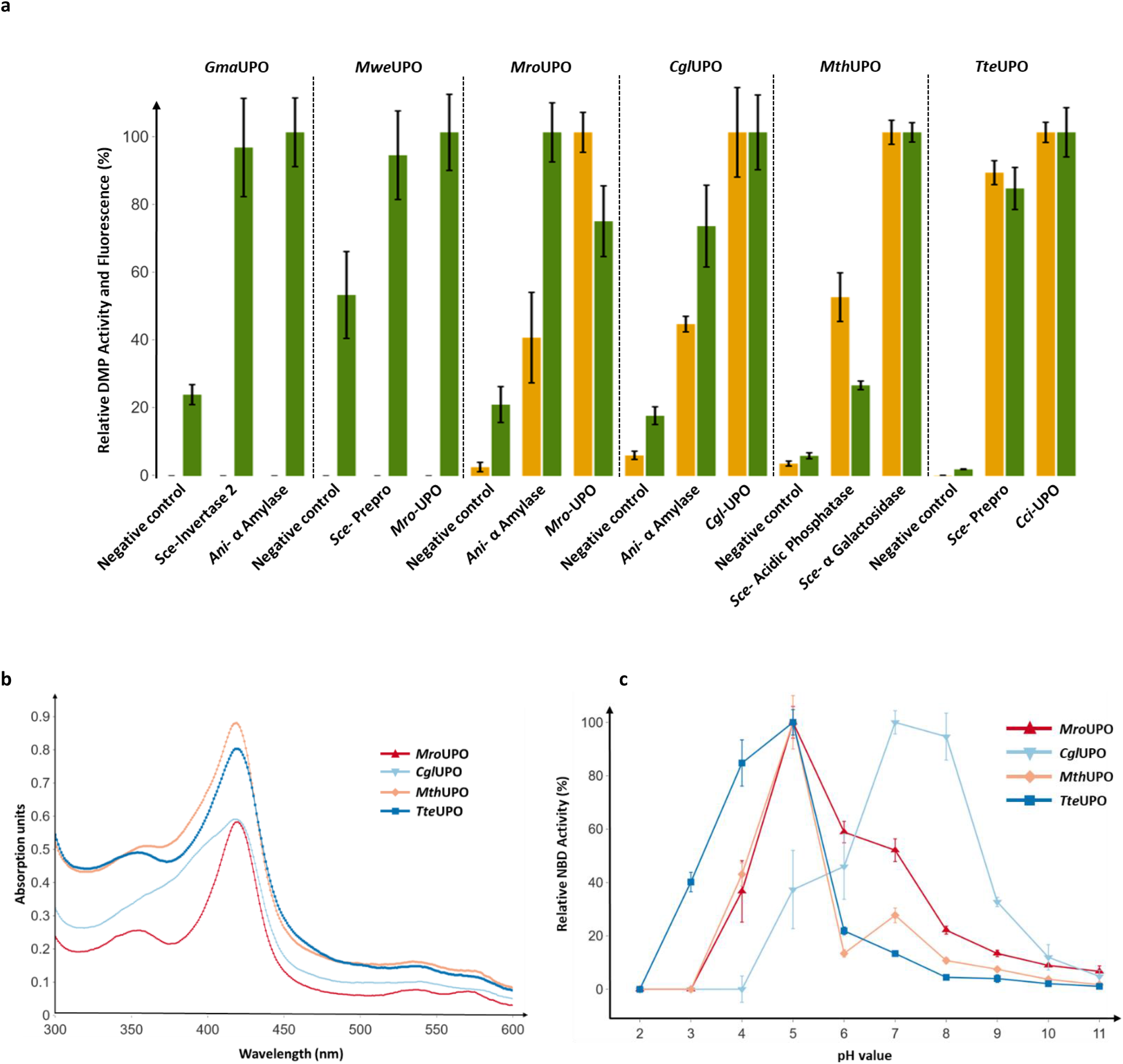
Through signal peptide shuffling identified novel UPO construct and their analysis of UVabsorption spectra and pH profiles. **a** Golden Gate signal peptide shuffling was applied for the testing of described and putative UPO genes, and the two best signal peptide/UPO gene combinations are displayed. *Gma*UPO, *Mwe*UPO, *Mro*UPO and *Cgl*UPO were screened in combination with a GFP11-tag. *Mth*UPO and *Tte*UPO were screened using the TwinStrep-GFP11 protein tag. UPO enzyme activity was determined by monitoring the conversion of 2,6-dimethoxyphenol (DMP) to cerrulignone. The highest average fluorescence (split-GFP) and conversion values (DMP) within one enzyme panel were set to 100 %, and the other values normalised accordingly. Data are mean ± s.d. of biological replicates (n=4). Corresponding primary data are displayed in Supplementary Table 9. **b** UV-Vis absorption spectra of the purified peroxygenases *Mro*UPO, *Cgl*UPO, *Mth*UPO and *Tte*UPO in the wavelength range between 300 and 600 nm (measurement interval: 1 nm). **c** pH profiles of *Mro*UPO, *Cgl*UPO, *Mth*UPO and *Tte*UPO catalysed enzymatic conversion of 5-nitro-1,3-benzodioxole (NBD) to 4-nitrocatechol. The highest mean activity of a respective enzyme was set to 100 % and the other values normalised accordingly. Data are means ± s.d. of measurements performed in triplicates. Corresponding primary data are displayed in Supplementary Table 10.

### Utilisation of the modular system for the heterologous production of novel UPOs

To demonstrate that the modular system can provide quick access to UPOs, we picked seven UPO genes to be expressed in *S. cerevisiae* with three being undescribed putative UPOs. Four UPOs were previously described and were produced in their natural hosts—*Marasmius rotula* UPO (*Mro*UPO)^36^, *Marasmius wettsteinii* UPO (*Mwe*UPO)^5^, *Chaetomium globosum* UPO (*Cgl*UPO)^37^—or are heterologously expressed in an *Aspergillus oryzae* strain (*Coprinopsis cinerea* UPO (*Cci*UPO))^38^.

Two putative UPO sequences were selected based on sequence alignments and data bank searches using the short-type peroxygenase *Cgl*UPO as a template. Two sequences were retrieved from fungi classified as thermophilic: *Myceliophthora thermophila* (*Mth*UPO) and *Thielavia terrestris* (*Tte*UPO)^39^, bearing 72 % and 51 % sequence identity to *Cgl*UPO, respectively. The predicted long-type UPO gene *Gma*UPO derived from the basidiomycete *Galerina marginata* and was selected based on its high sequence identity (71 %) with *Aae*UPO.

All genes were introduced as modules (Module 2) into the Golden Gate system and subjected to random shuffling utilising all 17 signal peptides (Module 1).

Out of the seven UPO genes, six were secreted in *S. cerevisiae* in combination with at least two signal peptides (Fig. 2a). *Cci*UPO showed no secretion with any of the signal peptides. *Mwe*UPO and *Gma*UPO were identified by the split-GFP assay, but no activity was detected using the colourimetric 2,6-dimethoxyphenol (DMP) assay^8^. *Mwe*UPO, *Mro*UPO and *Cgl*UPO were the only UPOs, which showed the highest activities with their endogenous signal peptides, where *Mro*UPO and *Mwe*UPO share the same native signal peptide. *Mth*UPO and *Tte*UPO showed remarkable secretion levels in the microtiter plate setup, leading to 17-fold (*Mth*UPO) and 50-fold (*Tte*UPO) split-GFP signal intensities above background level. A high signal peptide promiscuity was observed for *Mth*UPO and *Tte*UPO with at least 5 and 8 suitable signal peptides, respectively (Supplementary Figs. 3 and 4).

**Fig. 3.**
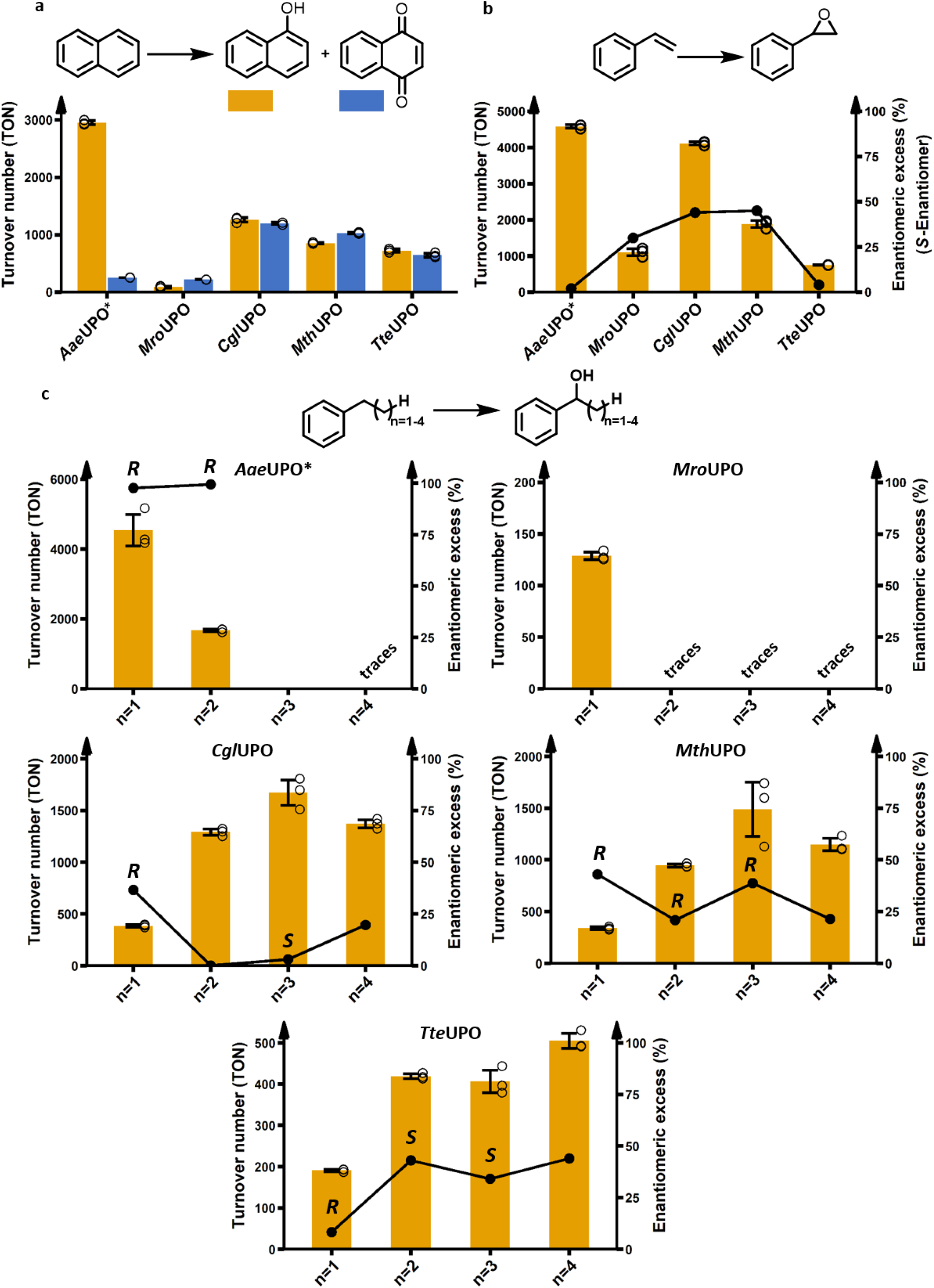
Enzymatic activity assessment of the peroxygenases regarding aromatic hydroxylation, epoxidation and sp3-carbon hydroxylation. All reactions were performed for 1 h using 1 mM of substrate. Bar charts display the obtained turnover numbers (TON) within one hour. The lines correspond to the enantiomeric excess %. Data are mean ± s.d. measurements derived from biological triplicates with individual data points shown as circles. See supplementary information for further details. **a** Conversion of naphthalene to naphthol and 1,4-naphthoquinone. **b** Conversion of styrene to styrene oxide. **c** A homologous row of phenylethane, phenylpropane, phenylbutane and phenylpentane hydroxylation, respectively focusing on hydroxylation of the benzylic carbon. The alcohol enantiomer is indicated by an (R) or (S). The exact enantiomer for phenylpentane was not determined. See Supplementary Fig. 14 for occurrence of side-products. For MroUPO conversion of phenylethane no enantioselectivity could be determined.

**Fig. 4.**
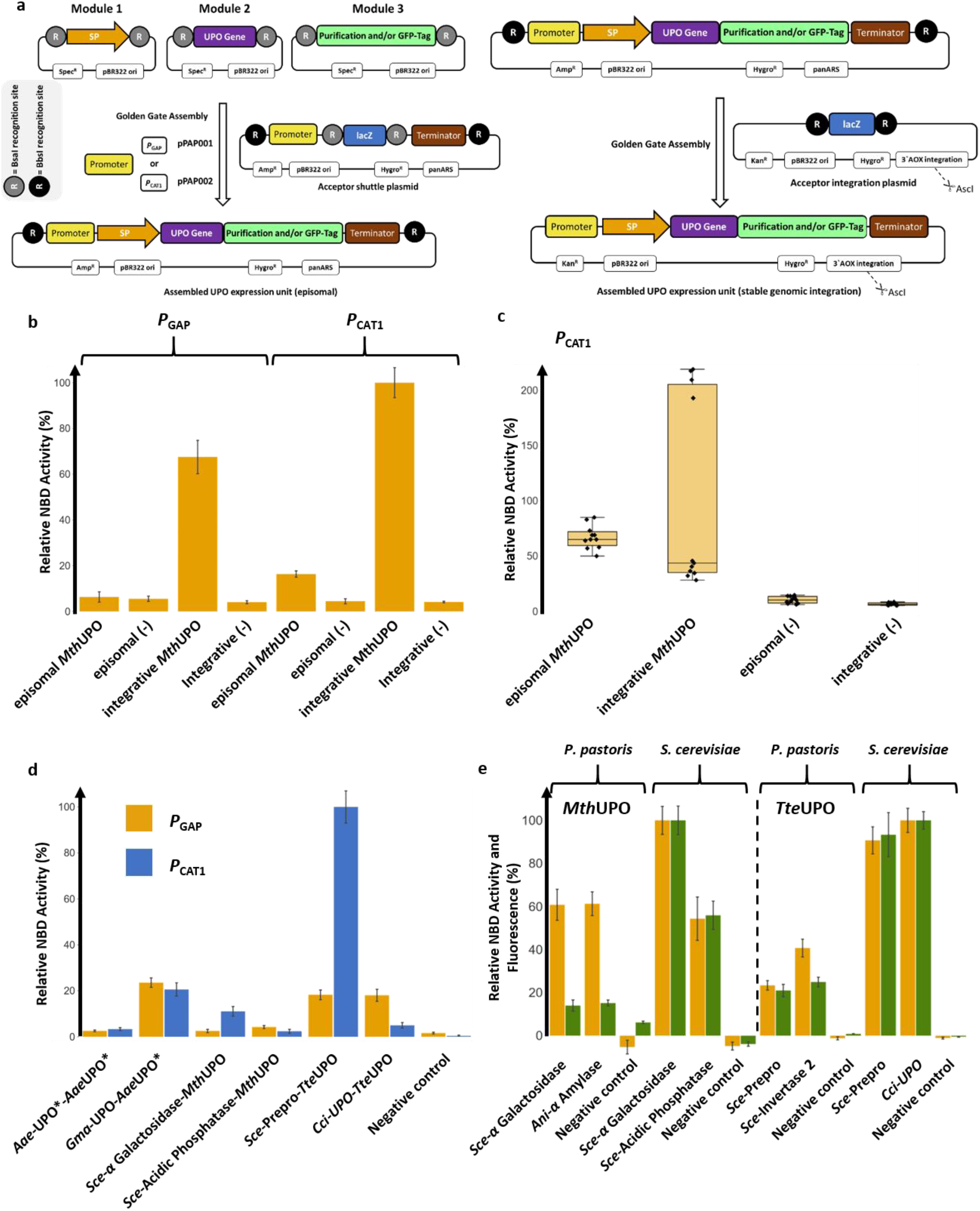
The compatible modular Golden Gate setup utilising episomal and integrative *P. pastoris* plasmids and its application. **a** *Left*: Overview of the designed episomal *P. pastoris* screening setup. All previously created basic modules are compatible to be used within this system. Two episomal plasmids wer e designed harbouring the constitutive strong promoter *P*_GAP_ or the strong inducible promoter *P*_CAT1_. *Right:* Identified gene constructs can be directly transferred in a one-pot Golden Gate reaction (*Bbs*I) from the episomal plasmid to an integrative plasmid. After linearisation (AscI digest) this plasmid can be integrated into the genomic 3’AOX region of *P. pastoris*. **b** Comparison of relative activities of 5-nitro-1,3-benzodioxole (NBD) conversion for different *P. pastoris* constructs bearing the tripartite combination of α Galactosidase signal peptide-*Mth*UPO-TwinStrep-GFP11. *P*_GAP_ bearing constructs were screened utilising Glucose (1.5 % (w/v)) as sole carbon source. *P*_CAT1_ bearing constructs were screened utilising a dual feeding strategy (0.5 % (v/v) glycerol and 1.5 % (v/v) methanol) as primary and inducible carbon sources. The highest expression mean is set to 100 % and all data normalised. Data are mean ± s.d. of biological replicates (n=6) originating from streak outs of one previously screened colony of the respective construct. **c** Comparison of relative activities of NBD conversion of *P*_CAT1_ based constructs bearing the tripartite combination of α Galactosidase signal peptide-*Mth*UPO-TwinStrep-GFP11. Box plots of biological replicates (n=11) of individual *P. pastoris* colonies for each construct. The highest expression mean is set to 100 % and all data normalised (episomal *Mth*UPO: median = 65, s.d. 15.0 %; integrative *Mth*UPO: median = 44, s.d. 83.3 %; episomal (−): median = 10, s.d. 28.2 %; integrative (−): median = 6, s.d. 16.4 %). **d** Comparison of relative activities of NBD conversion for different episomal *P. pastoris* constructs (6 biological replicates each) using the indicated signal peptide-UPO combinations as well as a TwinStrep-GFP11 tag. *P*_GAP_ (yellow bars) and *P*_CAT1_ (blue bars). The highest expression is set to 100 %, and all data are normalised. Data are mean ± of biological replicates (n=6). e Direct comparison of episomal UPO production of the two best signal peptide-UPO combinations for *Mth*UPO and *Tte*UPO as identified by a previously performed signal peptide shuffling approach in both yeast species. Episomal *P. pastoris* expressions utilised the *P*_CAT1_. The highest mean expression and activity for each enzyme is set to 100 %, and all data are normalised. Data are mean ± s.d. of biological replicates (n=6). NBD conversion activity (orange) and relative fluorescence units (green). All primary data are displayed in Supplementary Tables 11-14.

### Purification and characterisation of the identified UPOs

All secreted UPOs in combination with their best signal peptides were equipped with the TwinStrep-GFP11 tag, produced in 1 L shake flask scale and purified by affinity chromatography. The occurrence and primary sequence of each UPO was confirmed by tryptic digest and mass spectrometric peptide analysis (Supplementary Table 6). *Aae*UPO* analysis revealed the amino acids ‘EPGLPP’ being the first detectable residues at the *N*-terminus in accordance with previous results^40^. This finding indicates that the new signal peptide *Gma*-UPO leads to a comparable cleavage pattern as the evolved *Aae*-UPO* signal peptide. The split-GFP response and the NBD activity also exhibited the same ratio for both signal peptides (Fig. 1c), which further strengthens the point of a similar cleavage pattern. Both UPOs were produced utilising their native signal peptide (signal peptide annotated as *Mro*-UPO in both cases). Fragments derived from the signal peptide *Mro-*UPO (11 amino acids for *Mro*UPO and 9 amino acids for *Mwe*UPO) were identified by MS analysis, suggesting a different cleavage pattern compared to the natural host^5^. Obtained *N*-termini of *Gma*UPO and *Mth*UPO are in agreement with the predicted cleavage sites based on alignments with the enzymes *Aae*UPO* and *Cgl*UPO, respectively. The *N*-terminus of *Cgl*UPO could not be resolved. For *Tte*UPO, a peptide fragment of 10 amino acids of the utilised signal peptide (*Sce*- Prepro) was identified.

*Gma*UPO and *Mwe*UPO were not further studied as the purified enzymes did not exhibit any activity towards the colourimetric peroxygenase substrates DMP and NBD. Biochemical parameters were therefore determined for *Mro*UPO, *Cgl*UPO, *Mth*UPO and *Tte*UPO. UV absorption profiles showed the expected characteristic peroxygenase haem-thiolate features. A Soret band with a maximum around 420 nm (*Mro*UPO: 419 nm; *Cgl*UPO: 418 nm; *Mth*UPO: 420 nm and *Tte*UPO: 419 nm) and two Q-bands in the range of 537 to 546 and 569 to 573 nm (Fig. 2B)^2^ were detected. *Cgl*UPO revealed a broader Soret band shape as well as less pronounced Q-bands. The respective carbon monoxide complex exhibited absorption maxima around 444 nm with a less detectable signal for *Mro*UPO (Supplementary Fig. 5).

**Fig. 5.**
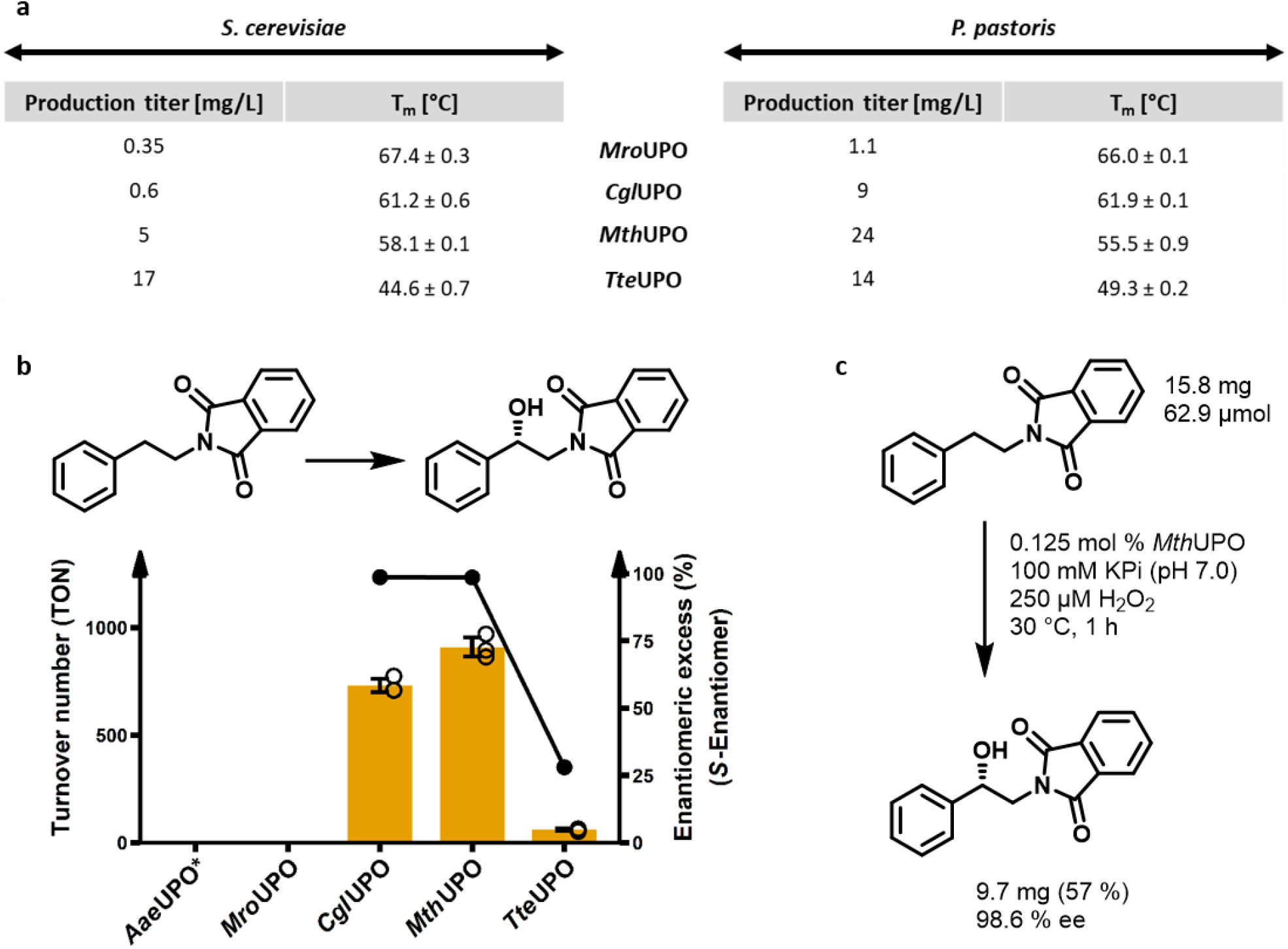
Expression yields and thermostabilities of UPOs derived from the different yeast systems and conversion of a phenethylamine derivative on analytical and preparative scale. **a** Comparison of volumetric production titre of recombinant UPOs in shake flask scale (1 L) between *S. cerevisiae* (episomal construct) and *P. pastoris* (integrative construct) as obtained after ultrafiltration of the respective culture supernatant. UPOs were produced and secreted utilising their natural signal peptide (*Mro*UPO and *Cgl*UPO) or a previously identified suitable exogenous signal peptide *Mth*UPO (*Sce*- α Galactosidase) and TteUPO (*S. cerevisiae*: *Cci*-UPO; *P. pastoris*: *Sce*- Prepro). For all *P. pastoris* production setups the methanol inducible promoter *P*_CAT1_ was utilised. Thermal denaturation midpoints (T_m_) for the four UPOs produced in both organisms were determined in biological triplicates using purified protein samples (in 100 mM potassium phosphate; pH 7.0) using differential scanning fluorimetry (DSF). **b** Bar chart showing turnover number within one hour for the benzylic hydroxylation of *N*-phthaloyl-phenethylamine by *P. pastoris* produced *Aae*UPO*, *Mro*UPO, *Cgl*UPO, *Mth*UPO and *Tte*UPO. Turnover data are mean ± s.d. of measurements made in triplicates. TON determined by GC-MS and *ee* % by chiral HPLC (Supplementary Figs. 16-18). **c** Preparative scale conversion of *N*-phthalimide protected phenethylamine using *P. pastoris* produced *Mth*UPO.

Protein purity and glycosylation were analysed by SDS-PAGE. Native deglycosylation was performed using PNGaseF (Supplementary Fig. 6). All obtained molecular weights after deglycosylation were in agreement with the calculated primary sequence and the peptide analysis by mass spectrometry. *Mro*UPO exhibited a weak band at approx. 42 kDa that was retained after deglycosylation. *Cgl*UPO revealed a smeared band in the range of 55 to 130 kDa. Deglycosylation led to the occurrence of two distinct protein bands of approx. 37 and 33 kDa indicating different protein subtypes. *Mth*UPO and *Tte*UPO showed an intensive smeared band in the range of 55 to 200 kDa. This smear was converted into distinct protein bands upon deglycosylation with approx. 38 kDa and 36 kDa for *Mth*UPO and *Tte*UPO, respectively.

To gain insights into the impact of the glycosylation for enzymes’ activities, the UPOs were deglycosylated in the native state and assessed for their activity towards NBD (Supplementary Fig. 7). The enzymatic activity of *Cgl*UPO and *Mro*UPO was not significantly and only by 30 % reduced, respectively. The activity was substantially impaired for *Tte*UPO and *Mth*UPO, leading to a complete loss and approx. 85 % decrease, respectively, in enzymatic activity.

We next evaluated the pH-dependencies of the enzymes using NBD as a substrate (Fig. 2c). *Mro*UPO, *Mth*UPO and *Tte*UPO exhibited a similar profile with maximum activity at slightly acidic conditions (pH 5), whereas *Cgl*UPO’s activity optimum was at pH 7. *Tte*UPO showed a broader tolerance towards lower pH values, retaining medium (pH 3; 40 %) and high activity (pH 4.0; 80 %) at acidic conditions. The obtained values for *Mro*UPO and *Cgl*UPO are in good agreement with previous data obtained with homologously produced enzyme^36,37^.

### Enzymatic epoxidation and hydroxylation experiments

The heterologously produced UPOs were tested towards their substrate specificity and activities by investigating three distinct reaction types: aromatic hydroxylation (sp^2^-carbon), epoxidation of an alkene and the benzylic hydroxylation (sp^3^-carbon) of phenylalkanes with varying alkyl chain lengths from two to five carbons (Fig. 3). All reactions were performed at the same conditions and assessed for the achieved turnover number (TON) within one hour. Substantially differing behaviour could be observed between *Aae*UPO* and the novel heterologously produced UPOs regarding substrate conversion, specific product formation and stereoselectivity. *Aae*UPO* proved to be the only enzyme displaying a high specificity for single hydroxylation of naphthalene leading to 1-naphthol (92 % of the formed product, Fig. 3a). The other UPOs exhibited a strong tendency for further oxidation leading to the dione product 1,4-naphthoquinone. The epoxidation of styrene (Fig. 3b) was efficiently catalysed by *Aae*UPO* (4580 TON) in combination with a poor stereoselectivity (2 % *ee*). *Cgl*UPO exhibited comparable epoxidation activities (4110 TON) and an enantioselectivity of 44 % *ee*. For *Mth*UPO, TON decreased to 1100 but revealed the highest stereoselectivity (45 % *ee*). The studies of the benzylic hydroxylation of phenylalkanes—phenylethane to phenylpentane—confirmed the preference of *Aae*UPO* towards short alkane chain length (Fig. 3c)^3^. Starting from 4500 turnovers for the conversion of phenylethane and deteriorating to no product formation and only traces of benzylic hydroxylation using phenylbutane and phenylpentane, respectively (for other product formations see Supplementary Fig. 15). *Cgl*UPO and *Mth*UPO exhibited an inverted trend with increasing product formations for longer alkyl chain lengths, exhibiting the lowest activity for the phenylethane hydroxylation.

The highest activity was detected in both cases using phenylbutane (*Cgl*UPO: 1670 TON, *Mth*UPO: 1490 TON) with only slightly decreased activity for phenylpentane as a substrate and the only significant side-product being the further oxidation of the benzylic alcohol to the corresponding ketone (Supplementary Fig. 15). *Tte*UPO showed a similar preference towards long-chain phenylalkanes with the highest TON for phenylpentane conversion (500 TON). *Tte*UPO represented the only UPO with a significant specificity towards the formation of the *S*-enantiomer for phenylpropane and phenylbutane. For phenylpentane, it revealed the formation of the opposite alcohol enantiomer than the other tested UPOs.

### Expanding the modular UPO secretion system to *Pichia pastoris*

The methylotrophic yeast *P. pastoris* (syn. *Komagataella phaffii*) constitutes an attractive heterologous production host with a steadily growing toolbox of valuable synthetic biology parts such as plasmids, promoters and signal peptides^41,42^. *P. pastoris* can reach high cell densities, efficiently perform post-translational modifications such as glycosylation and disulfide-linkage and offers a set of strong and tightly regulated promoters for target gene expression. Amongst other factors, these properties render *P. pastoris* a widely used eukaryotic host for the large scale industrial production of therapeutic proteins and industrial enzymes^43^. We investigated the adaptation of the modular system for use in *P. pastoris*. Therefore two novel episomal *P. pastoris* expression plasmids were designed and assembled. They contain a previously described autonomously replicating sequence coined panARS, which confers episomal stability and a hygromycin B marker gene for antibiotic selection^44,45^. The constructed episomal plasmids differ by the employed promoter: the strong constitutive glyceraldehyde-3-phosphate dehydrogenase promoter (*P*_GAP_, plasmid pPAP001) and the recently described strong methanol inducible catalase promoter (*P*_CAT1,_ pPAP002)^46^. The plasmids were constructed to allow direct implementation of the tripartite modular UPO secretion system, consisting of Module 1 (signal peptide), Module 2 (UPO gene) and Module 3 (*C*-terminal tag, Fig. 4a; left). To further allow the genomic integration to generate stable *P. pastoris* cell lines for antibiotic-free large scale enzyme production in shake flasks or fermenters, a third plasmid (pPAP003) was constructed. The episomal plasmids are designed to enable direct transfer of the identified best transcription unit (promoter-signal peptide-gene-tag-terminator) combination to the integration plasmid. This transfer requires only an additional Golden Gate assembly reaction using the restriction enzyme *Bbs*I (Fig. 4a; right).

We tested all *P. pastoris* plasmids using the newly discovered peroxygenase *Mth*UPO in combination with the *Sce*-α Galactosidase signal peptide. The constructs proved to be functional and led to an NBD conversion signal (Figure 4b). *P*_GAP_ based secretion was generally lower in comparison to *P*_CAT1_, and the episomal *P*_GAP_ UPO activity was not distinguishable from the negative control. The integrative plasmids outperformed their episomal counterparts significantly with a factor of 5 for *P*_CAT1_. A similar observation however in a varying degree was made testing the enzymes *Aae*UPO* and *Tte*UPO (Supplementary Fig. 8), indicating that the integrative constructs are promoting higher UPO secretion levels than their episomal counterparts.

To gain insights into interclonal variabilities of UPO secretion, episomal and integrative plasmids were transformed into *P. pastoris.* Individual colonies were cultivated and tested for UPO secretion. The episomal construct showed diminished mean activity but a substantially lower clonal variability than the integrative plasmid when tested with NBD (Fig. 4c). This high variability of the secretion level for the integrative plasmid is presumably due to different numbers of copy insertions into the *P. pastoris* genome, which might also lead to different colony sizes (Supplementary Fig. 9).

To investigate and compare the secretion levels of episomal *P*_GAP_ and *P*_CAT1_ harbouring plasmids, twelve constructs were generated harbouring the peroxygenases *Aae*UPO*, *Mth*UPO and *Tte*UPO. All promoter combinations (*P*_GAP_ and *P*_CAT1_) and the two previously identified signal peptides were constructed in combination with the respective UPO gene and analysed for NBD activity. All constructs resulted in a significant NBD conversion (Fig. 4d). The previously observed 220 % improved *Aae*UPO* secretion in *S. cerevisiae* by combining *Aae*UPO* with the signal peptide *Gma-*UPO was found to be even more pronounced using the episomal *P. pastoris* constructs (*P*_CAT1_: 620 %). Besides the striking influence of the promoter on the secretion level, also the combination of the signal peptide and the promoter proved to be pivotal. For *Tte*UPO, using the promoter *P*_CAT1_ in combination with the *Sce*- Prepro signal peptide led to the highest detected activity with a 20-fold higher signal compared to the *Cci*-UPO signal peptide. The same signal peptide variations employing the *P*_GAP_ promoter, however, resulted in similar secretion levels. This demonstrates besides the crucial role of the chosen signal peptide (Fig. 1c and 2a) an additionally pivotal influence of the promoter/signal peptide combination on the UPO secretion.

To gain insights into the different signal peptide preferences for secretion in *P. pastoris*, the signal peptide shuffling approach was repeated in *P. pastoris*, choosing the episomal *P*_CAT1_ bearing plasmid (Supplementary Figs. 10 and 11). For *Mth*UPO the signal peptides *Sce*- α Galactosidase and *Ani*- α Amylase proved to be most suitable, and *Sce*- Prepro and *Sce*- Invertase 2 were identified as top hits for *Tte*UPO. Interestingly, *Sce*- Invertase 2 has not been identified amongst the top hits in *S. cerevisiae* whereas the best signal peptide (*Cci*-UPO) for secretion in *S. cerevisiae* (Fig. 4d) was not identified in the *P. pastoris* screen.

To compare episomal *S. cerevisiae* and *P. pastoris* secretion, the two best performing constructs for *Mth*UPO and *Tte*UPO were selected. This species comparison (Fig. 4e) indicates that the episomal *S. cerevisiae* secretion is superior to the episomal *P. pastoris* production. In the case of *Mth*UPO, both *P. pastoris* constructs led to approx. 60 % of NBD conversion in comparison to the most suitable *S. cerevisiae* construct while already exhibiting higher NBD conversion rates than the second most suitable signal peptide for secretion in *S. cerevisiae* (*Sce*- Acidic Phosphatase). The split-GFP fluorescence assay revealed a diminished response for the *P. pastoris* set up relative to the *S. cerevisiae* constructs. Regarding *Tte*UPO, the best *P. pastoris* construct (*Sce*-Invertase 2) led to approx. 40 % of relative NBD conversion when compared to the best *S. cerevisiae* construct (*Cci*-UPO). For *Tte*UPO the split GFP assay followed a linear pattern when comparing species, without revealing a diminished response for *P. pastoris*.

### Comparison of shake flask UPO production in *P. pastoris* and *S. cerevisiae*

By using the constructed integrative plasmid pPAP003 and the *P*_CAT1_ promoter, stable *P. pastoris* cell lines were constructed for the production of five UPOs: *Aae*UPO*, *Mro*UPO, *Cgl*UPO, *Mth*UPO and *Tte*UPO (Fig. 5a). Utilising *P. pastoris* led to substantially higher production titres in all cases, except for *Tte*UPO. The rather low yields of *Mro*UPO and *Cgl*UPO produced in *S. cerevisiae* could be increased substantially when using *P. pastoris* (*Mro*UPO: 3-fold, *Cgl*UPO: 15-fold). The *Mth*UPO production yield was improved 5-fold when produced in *P. pastoris*. Regarding *Tte*UPO, the product titre was decreased in *P. pastoris* by approx. 20 %, however, still maintaining an overall high yield. The production titres of *S. cerevisiae* derived *Tte*UPO (17 mg/L), and *P. pastoris* derived *Mth*UPO (24 mg/L) are the highest yields for shake flask cultivation of recombinant fungal peroxygenases reported thus far. The transfer of the *Pichia* expression system to a fed-batch bioreactor might yield production levels at the g/L scale, due to the higher cell densities achievable within this format as already demonstrated^40^. All proteins were purified using the TwinStrep-tag and analysed by SDS-PAGE (Supplementary Fig. 12). Highly pure enzyme preparations were obtained after one-step TwinStrep purification. Based on the successful production in both organisms, thermostability values (denaturation midpoint; T_m_) of the four UPOs were assessed using differential scanning fluorimetry (Fig. 5a). Comparing the obtained values of the respective UPOs derived from both organisms were similar to a variation of 0.7 to 4.7 °C. The highest thermostability values were obtained for *Mro*UPO with 67.4 and 66.0 °C for *S. cerevisiae* and *P. pastoris*, respectively. The two UPOs derived from thermophilic fungi, *Mth*UPO and *Tte*UPO, exhibited no superior thermostability when compared to the closest related enzyme *Cgl*UPO. *Tte*UPO revealed the lowest thermostability in the tested group with 44.6 and 49.3 °C for *S. cerevisiae* and *P. pastoris*, respectively.

### Enantioselective hydroxylation of an *N*-protected phenethylamine on a preparative scale

To gain insights into the ability of the enzymes to convert industrially relevant molecules in an enantioselective manner, we selected *N*-protected phenethylamine as a substrate. The hydroxylation of phenethylamine derivatives at the benzylic position provides access to a plethora of pharmaceutically important classes like beta-blockers and sympathomimetics^47^.

The peroxygenases *Aae*UPO*, *Mro*UPO, *Cgl*UPO, *Mth*UPO and *Tte*UPO were produced in *P. pastoris*, purified and assessed for their activity on *N*-phthaloyl-phenethylamine. *Aae*UPO* and *Mro*UPO exhibited no formation of the benzylic alcohol product, and *Tte*UPO performed 60 TON within an hour while achieving an enantioselectivity of 28 % *ee* (Fig. 5b, Supplementary Figs. 16 and 18). *Cgl*UPO and *Mth*UPO revealed the highest activities with 730 and 908 TON, respectively, within the 1 h reaction setup (Supplementary Fig. 16). *Mth*UPO showed an over-oxidation to the ketone amounting to 222 TON (Supplementary Fig. 17). The enantioselectivity proved to be excellent for *Cgl*UPO and *Mth*UPO with 98.7 % *ee* and 98.6 % *ee* (Supplementary Fig. 18).

Harnessing the high production titre of *Mth*UPO in *P. pastoris* (24 mg/L) in combination with the previously observed good substrate conversion and high enantioselectivity we aimed for the proof of concept enantioselective synthesis of (*S*)-(+)-2-*N*-Phthaloyl-1-phenylethanol on a preparative scale (Fig. 5c). In a first upscaling reaction (300 mL total volume) 0.125 mol % of *Mth*UPO derived without further purification from concentrated *P. pastoris* supernatant were used as catalyst loading. The upscaled reaction (30 °C; 1 h) led to the synthesis of 9.70 mg (S)-(+)-2-*N*-Phthaloyl-1-phenylethanol (57 % purified yield) and an enantiomeric excess of 98.6 % (Supplementary Fig. 18).

## Discussion

Fungal unspecific peroxygenases (UPOs) have gained substantial interest as versatile hydroxylation catalyst since their initial discovery 16 years ago^2^. The most significant limitation for the wider application of UPOs arguably remains the heterologous production utilising a fast-growing host. Thus far, only one UPO could be produced and engineered within a system amenable to high throughput: the *S. cerevisiae* secretion variant *Aae*UPO*^6^.

Building on the therein developed expression setup, we started our endeavour to construct a versatile UPO secretion system. The constructed Golden Gate-based platform consists of a signal peptide library (Module 1), UPO genes (Module 2) and protein-tags (Module 3). This format enabled the first report of successful yeast secretion of six UPOs—two of them (*Mth*UPO and *Tte*UPO) derived from genome, and secretome data had not been characterised as UPOs before^39^. The whole expression platform could be subsequently transferred to *P. pastoris*, resulting in excellent UPO expression yields allowing for preparative scale hydroxylation reactions.

Since the only enzyme out of the panel that could not be produced (*Cci*UPO) belongs to the group of long-type UPOs, and it previously took considerable effort to engineer the long-type UPO *Aae*UPO towards secretion in yeast, one could argue that the heterologous production of long-type UPOs seems to be more challenging. In fact, *Mro*UPO, *Cgl*UPO, *Mth*UPO and *Tte*UPO, which could be initially produced in yeast and characterised within the scope of this work, all belong to the class of short-type UPOs. Recent work in our laboratory suggests that gene shuffling of long-type UPOs can offer a viable option to obtain a library of active and structurally diverse long-type UPOs expanding the panel of available recombinant peroxygenases^48^. The hypothesised pivotal role of the employed signal peptides for the successful secretion of UPOs was manifested within this study. Even in case of the laboratory evolved peroxygenase *Aae*UPO* harbouring an evolved signal peptide, the secretion could be further improved utilising the signal peptide *Gma-*UPO derived from another UPO by 2.2-fold in *S. cerevisiae* and even 6-fold in *P. pastoris*. The significance of a suitable signal peptide−gene combination is moreover underlined by the observation that almost all of the 17 tested signal peptides proved to be functional in combination with at least one UPO gene. The studies revealed substantial differences in signal peptide acceptance and promiscuity among the different UPOs. Whereas *Aae*UPO* only showed pronounced activity in combination with two signal peptides (*Aae*-UPO* and *Gma-*UPO) other UPOs like *Mth*UPO and *Tte*UPO could be produced when combined with multiple and diverse signal peptides regarding sequence length and composition of the overall tested panel. Although all previously identified signal peptide−gene combinations led to productive secretion in *Pichia pastoris* as well, subtle differences and preferences were shown by a signal peptide shuffling for *Mth*UPO and *Tte*UPO in *P. pastoris.* Interestingly, the α factor leader signal peptide (*Sce*-Prepro), which is often used as a gold standard signal peptide for target protein secretion in *P. pastoris*^42,43^, was only identified as a top hit in combination with *Tte*UPO.

The GFP11 detection tag proved to be an indispensable asset to distinguish secretion from activity^30^. Between different UPOs, the variation in fluorescence could be further pronounced based on different accessibilities of the split-GFP-tag. This tendency was shown for *Aae*UPO* where the TwinStrep-GFP11 tag (59 amino acids) yielded a 4-fold increased signal intensity relative to the shorter GFP11 tag (27 amino acids). In some cases, like *Cgl*UPO (Fig 2a), the fluorescence response greatly differed from the activity depending on the employed signal peptide. This observation might be explained by different cleavage patterns at the *N*-terminus depending on the respective signal peptide leading to slightly altered overall structures and hence activities of the mature, processed enzyme.

The peroxygenases *Tte*UPO and *Mth*UPO displayed unprecedented expression levels in *S. cerevisiae* and *P. pastoris*, even outperforming the secretion engineered enzyme *Aae*UPO*^6^. The adaptation to episomal plasmid expression in *P. pastoris* proved that the entire modular system could be readily transferred to other organisms. The application in *P. pastoris* furthermore paves the way towards future-directed evolution enterprises entirely performed in *P. pastoris*, further streamlining the workflow from gene discovery to construct identification and large scale protein production. In comparison to the *S. cerevisiae* based episomal system, the *P. pastoris* based episomal plasmid expression of *Mth*UPO retained 60 % of the activity (Fig. 4e). However, there is still plenty of potential for *P. pastoris* production optimisation utilising different promoters, carbon sources, induction and co-feeding strategies^46,49^. A substantial influence of the promoter−signal peptide combination was observed, as shown for *Tte*UPO in Fig. 4d. This impact represents an aspect that can be further investigated in detail, for example, by expanding the modular system, including an additional shuffling library for promoters. Production of *Mth*UPO utilising the integrative plasmid led to substantially improved production when compared to the episomal counterpart. Nevertheless, the obtained interclonal variation is substantial, rendering the episomal plasmid expression more suitable for high throughput endeavours (Fig. 4c).

The relevance of expanding the scope of recombinant UPOs is reflected by the fact, that *Cgl*UPO, *Mth*UPO and *Tte*UPO displayed a different substrate specificity when compared to the well-characterised enzyme *Aae*UPO* (Fig. 3). For the benzylic hydroxylation of the homologous phenylalkane row, ranging from phenylethane to phenylpentane *Aae*UPO* displayed the highest activity on phenylethane and only traces of product for phenylbutane and phenylpentane conversion. The enzymes *Cgl*UPO, *Mth*UPO and *Tte*UPO on the contrary, displayed the lowest activities for phenylethane and highest for phenylbutane and –pentane conversion, respectively. *Tte*UPO furthermore catalyses the formation of the opposite alcohol enantiomer compared to the other UPOs for the conversion of phenylpropane to phenylpentane. Good activities and excellent enantioselectivities could also be achieved when challenging the enzymes for the benzylic hydroxylation of *N*-phthalimide protected phenethylamine for *Cgl*UPO and *Mth*UPO. This observation is vastly different from *Aae*UPO*, displaying no product formation and no known enantioselective conversion of substrates of similar structure. High UPO production yields in *P. pastoris* enabled the preparative conversion of a phenethylamine derivative by *Mth*UPO.

In summary, these observations prove that the built workflow from UPO gene, followed by identification of suitable expression constructs via signal peptide shuffling in combination with high throughput screening in *S. cerevisiae* as well as *P. pastoris* and subsequent production upscaling can lead to highly enantioselective preparative product formations of pharmaceutically valuable building blocks.

In the future, this workflow could be applied to other UPO genes or generally genes of interest, which are suitable for production in yeast, especially for proteins that might require efficient post-translational modifications such as glycosylation and disulfide linkage. Besides target protein secretion, the expression plasmids also allow for intracellular production when no signal peptide is attached. To allow other researchers to harness the modular yeast system, we deposited all relevant plasmids (signal peptides, protein tags and expression plasmids) as a kit with the non-profit plasmid repository Addgene (*Yeast Secret and Detect* Kit).

## Acknowledgment

M.J.W and A.K. thank the Bundesministerium für Bildung und Forschung („Biotechnologie 2020+ Strukturvorhaben: Leibniz Research Cluster“, 031A360B and 031A360E) for generous funding. P.P. thanks the Landesgraduiertenförderung Sachsen-Anhalt for a PhD scholarship. J.M. thanks the Friedrich-Naumann-Stiftung for a PhD scholarship. Prof. Jürgen Pleiß (University of Stuttgart), Prof. Dirk Holtmann and Sebastian Bormann (DECHEMA Frankfurt) are kindly acknowledged for sharing putative and described UPO gene sequences. We would like to thank Cătălin Voiniciuc (Leibniz Institute of Plant Biochemistry Halle) for fruitful discussions regarding protein production in *Pichia pastoris* and furthermore providing Pichia plasmid parts and the X33-strain. Prof. Karin Breunig (Martin Luther University Halle-Wittenberg) is kindly acknowledged for generously supplying genomic DNA of *K. lactis*. We would like to especially thank Anja Ehrlich (Leibniz Institute of Plant Biochemistry Halle) for outstanding technical support and patience regarding chiral HPLC analyis of phenethylamine conversions. Furthermore, Prof. Markus Pietzsch and Dr. Franziska Seifert (Martin Luther University Halle-Wittenberg) are kindly acknowledged for discussions and providing access to the DSF device for thermostability measurements. Our sincere gratitude also goes to Prof. Martin Hofrichter and Dr. Harald Kellner (Technical University of Dresden) for discussions and providing the protein sequence of *Mwe*UPO. Dr. Swanhild Lohse (IPB Halle) is acknowledged for the initial construction of the utilised pAGT572 backbone structure of episomal *S. cerevisiae* plasmids.

## Author contributions

M.J.W and P.P. designed the research. P.P. performed all experiments apart from the enzymatic conversions in Figure 3 (performed by A.K.), Figure 5 (performed by J.M. and P.R.P. and co-designed by B.W.) and the protein identification by MS (performed by W.H.). P.P. and S.M. designed the modular Golden Gate yeast system and M.A. developed the 96-well *S. cerevisiae* expression system. M.J.W. and P.P. wrote the manuscript. All authors contributed to the proofreading of the manuscript.

## Materials & Correspondence

Correspondence and requests for materials should be addressed to M.J.W.

## Additional information

Supplementary information is available.

## Data availability

The authors declare that the data supporting the findings of this study are available within the paper and its Supplementary Information files. Source data are available from the corresponding author upon reasonable request.

## ONLINE METHODS

### Chemicals

Solvents were used as received without further purification. Ethyl acetate and acetone were utilised in GC ultra-grade (≥ 99,9 %) from Carl Roth (Karlsruhe, DE). Acetonitrile was purchased from Merck (Darmstadt, DE) in gradient grade for LC (≥ 99,9 %). Deuterated solvents for NMR spectroscopy were purchased from Deutero (Kastellaun, DE). All further reaction chemicals were purchased either from Sigma-Aldrich (Hamburg, DE), TCI Chemicals (Tokyo, JP), Merck (Darmstadt, DE), abcr (Karlsruhe, DE) or Fluka Chemika (Buchs, CH) and used as received. (R)-(+)-1-Phenyl-1-propanol (98 % purity) was purchased from abcr (Karlsruhe, DE). 1-Naphthol (99 % purity) was purchased from Merck (Darmstadt, DE). Naphthalene (99 % purity), 1,4-Naphthoquinone (97 % purity), Propyl benzene (99 % purity; GC), Butyl benzene (≥ 99 % purity), Ethyl benzoate (analytical standard),(R)-(+)-1-Phenyl-1-butanol (97 % purity), 2,6-Dimethoxyphenol (99 % purity), 5-Nitro-1,3-benzodioxole (98 % purity), Styrene (>99 % purity) and Hydrogen peroxide (30 % (v/v)) were purchased from Sigma-Aldrich. (±)-1-Phenyl-1-propanol (98 % purity) was purchased from Fluka Chemika. Pentyl benzene (> 98 % purity), Ethyl benzene (> 99 % purity), (±)-1-Phenyl-1-pentanol (> 98 % purity), (±)-1-Phenylethyl alcohol (> 98 % purity), (±)-1-Phenyl-1-butanol (> 98 % purity), (R)-(+)-1-Phenylethyl alcohol (> 98 % purity) and (±)-Styrene oxide (> 98 % purity) were purchased from TCI Europe (Eschborn, DE), all of them were used as received.

### Lab ware

Specialised 96-well half deep well microtiter plates for yeast cell growth and protein expression (model type: CR1496c) were purchased from EnzyScreen (Heemstede, NL) and were sealed with a fitting CR1396b Sandwich cover.

### Enzymes and cultivation media

For cultivation of *E. coli* cells terrific broth (TB) media from Carl Roth (Karlsruhe, DE) was used. For cultivation of *S. cerevisiae* cells D-Galactose, Peptone and Synthetic Complete Mixture (Kaiser) Drop-Out (-URA) were purchased from Formedium(Hunstanton, GB). Yeast nitrogen base (without amino acids) and Yeast extract were purchased from Carl Roth (Karlsruhe, DE). For *P. pastoris* cultivation Methanol (99,9 % Chromasolv purity grade) purchased from Honeywell Chemicals (Seelze, DE) was used as additional carbon source. PNGase F and BsaI were purchased from New England Biolabs (Ipswich, US). BbsI and FastDigest AscI were purchased from ThermoFisherScientific (Waltham, US) and T4 DNA Ligase from Promega (Madison, US).

### Bacterial and Yeast strains

For all cloning purposes and plasmid propagation, *E.coli* DH10B cells (ThermoFisherScientific, Waltham, US) were utilised. The protein sfGFP1-10 was produced utilising the *E.coli* BL21(DE3) strain (ThermoFisherScientific, Waltham, US) capable of T7 promoter dependent target protein expression. In the case of all work regarding *Saccharomyces cerevisiae,* the diploid strain INVSc1 (genotype: MATa his3D1 leu2 trp1-289 ura3-52 MAT his3D1 leu2 trp1-289 ura3-52) (ThermoFisherScientific, Waltham, US) was used. All work regarding *Pichia pastoris* was performed utilising the mut^+^ Strain X-33 (ThermoFisherScientific, Waltham, US).

### Oligonucleotides and Gene parts

All oligonucleotides were purchased in the lowest purification grade “desalted” and minimal quantity at Eurofins Genomics (Ebersberg, DE). The *Pichia pastoris* Cat1 promoter was purchased as a gene part from Twist Bioscience (San Francisco, US). The genes of the *Aae*UPO variant PaDa-I, *Gma*UPO, *Mwe*UPO and the sfGFP 1-10 gene were purchased as plasmid-cloned genes from Eurofins Genomics (Ebersberg, DE). The genes of *Cgl*UPO, *Mth*UPO and *Tte*UPO were retrieved as codon optimised (*S. cerevisiae* codon usage) gene strands from Eurofins Genomics.

### Achiral gas chromatography–mass spectrometry (GC-MS)

Measurements were performed on a Shimadzu GCMS-QP2010 Ultra instrument (Shimadzu, Kyoto, JP) using a SH-Rxi-5Sil MS column (30 m × 0.25 mm, 0.25 μm film, Shimadzu, Kyoto, JP) or OPTIMA 5MS Accent column (25 m × 0.20 mm, 0.20 μm film, Macherey-Nagel, Düren, DE) and helium as carrier gas. 1 μl of each sample was injected splitless with an injection temperature of 280 °C. The split/splitless uniliner inlets (3.5 mm, 5.0 × 95 mm for Shimadzu GCs, deactivated wool) from Restek (Bad Homburg, DE) were utilised and regenerated if needed by CS-Chromatography (Langerwehe, DE). The temperature program was adjusted, as shown in Supplementary Table 5. The interface temperature was set to 290 °C. Ionisation was obtained by electron impact with a voltage of 70 V, and the temperature of the ion source was 250 °C. The MS is equipped with dual-stage turbomolecular pumps and a quadrupole enabling a selected ion monitoring acquisition mode (SIM mode). Calibration and quantification w ere implemented in SIM mode with the corresponding *m/z* traces, as shown in Supplementary Table 5. The detector voltage of the secondary electron multiplier was adjusted in relation to the tuning results with perfluorotributylamine. The GC-MS parameter was controlled with GCMS Real Time Analysis, and for data evaluation, GCMS Postrun Analysis (GCMSsolution Version 4.45, Shimadzu, Kyoto, JP) was used.

### Chiral gas chromatography–mass spectrometry (GC-MS)

Measurements were performed on a Shimadzu GCMS-QP2020 NX instrument (Shimadzu, Kyoto, JP) with a Lipodex E column (25 m × 0.25 mm, Macherey-Nagel, Düren, DE) and helium as carrier gas. 1 μl of each sample was injected splitless with an OPTIC-4 (Shimadzu, Kyoto, JP) injector utilising a temperature profile in the liner (35 °C, 1 °C/s to 220 °C hold 115 s). The column temperature program was adjusted as shown in Supplementary Table 5. The interface temperature was set to 200 °C. Ionisation was obtained by electron impact with a voltage of 70 V, and the temperature of the ion source was 250 °C. The MS is equipped with dual stage turbomolecular pumps and a quadrupole enabling a selected ion monitoring acquisition mode (SIM mode). Calibration and quantification were implemented in SIM mode with the corresponding *m/z* traces, as shown in Supplementary Table 5. The detector voltage of the secondary electron multiplier was adjusted in relation to the tuning results with perfluorotributylamine. The GC-MS parameters were controlled with GCMS Real Time Analysis, and for data evaluation GCMS Postrun Analysis (GCMSsolution Version 4.45, Shimadzu, Kyoto, JP) was used.

### Column and analytic thin layer chromatography

All solvents for column chromatography were purchased from Merck Millipore (Darmstadt, DE) and distilled prior to use. Column chromatography was carried out using Merck silica gel 60 (40 – 63 μm). For analytic thin layer chromatography, Merck TLC silica gel 60 F254 aluminium sheets were used. Compounds were visualised by using UV light (254/366 nm).

### Nuclear magnetic resonance (NMR)

NMR spectra were recorded using a 400 MHz Agilent DD2 400 NMR spectrometer at 25 °C. The chemical shifts of 1H NMR spectra are referenced on the signal of the internal standard tetramethylsilane (δ = 0.000 ppm). Chemical shifts of 13C NMR spectra are referenced on the solvent residual signals of CDCl_3_ (δ = 77.000 ppm).

### Electrospray ionisation mass spectrometry (ESI-MS)

ESI mass spectra were recorded on an API3200 Triple Quadrupole mass spectrometer (AB Sciex) equipped with an electrospray ion source (positive spray voltage 5.5 kV, negative spray voltage 4.5 kV, source heater temperature 400 °C).

### Specific optical rotation

Specific optical rotations of compounds were recorded on a P-2000 Digital Polarimeter (JASCO, Pfungstadt, DE) utilising a wavelength of 589 nm.

### Chiral HPLC

HPLC chromatograms were recorded on an Agilent High Performance LC (Agilent Technologies, Waldbronn, DE). The used chiral column material was Chiralpak AS-H HPLC (Daicel, Tokyo, JP) (25 cm × 4.6 mm). Substances were dissolved in HPLC-grade isopropanol prior to analysis, and a sample volume of 5 μL injected. The eluent (20 % isopropanol, 80 % *n*-hexane) was used in a flow rate of 1 mL/min with the runtime of 30 min at 30 °C.

### Microwave reactions

Microwave reactions were carried out using an Initiator+ device (Biotage, Düsseldorf, DE).

