## Supplemental Information for "A modular two yeast species secretion system for the production and preparative application of fungal peroxygenases"

### I. General procedures

**Expression plasmid construction for *S. cerevisiae*.** A Level 1 Golden Gate based shuttle expression plasmid was constructed using a pAGT572 plasmid[1] as backbone structure, which can be propagated in *E.coli* as well as *S. cerevisiae*. The expression plasmid enables bacterial antibiotic selection (ampicillin resistance) and yeast auxotrophy selection (URA3 marker).To enable expression of a target gene a Gal 1.3 Promoter- a truncated, modified version of the widespread GAL1 Promoter is integrated upstream and a strong DIT1 terminator downstream of the cloning acceptor site. As placeholder for a target gene sequence a lacZ cassette (approx. 600 bp) is integrated, which enables β- galactosidase based blue/white selection of transformants based on the conversion of X-Gal. Upon digestion with BsaI the lacZ cassette is released, and a fitting open reading can be integrated in frame (e.g. Signal Peptide-Gene-C-terminal Tag) into the plasmid, thereby reconstituting a fully functional expression plasmid. The constructed expression plasmid was coined pAGT572_Nemo_2.0. Using the pAGT572 plasmid backbone and the GAL1 Promoter as units a second expression plasmid coined pAGT572_Nemo was constructed that follows the same functionality and principle but exhibits the original GAL1 promoter.

**Expression plasmid construction for *Pichia pastoris*.** The construction of suitable *P. pastoris* expression plasmids followed the same design principle- enabling a fully integrated shuffling system in both species. Two level 1 Golden Gate based shuttle expression plasmids were constructed, which can be propagated in *E.coli* (Ampicillin Resistance) as well as *P. pastoris* (Hygromycin B resistance). To enable episomal plasmid propagation in *P. pastoris,* the plasmids were equipped with a previously described functional ARS sequence[2], which was PCR amplified from *Kluveromyces lactis* genomic DNA. To enable expression of a target gene, the plasmids exhibit the strong constitutive GAP promoter (pPAP 001) or the strong methanol inducible promoter CAT1 (pPAP002), both in combination with a strong GAP terminator (tGAP). As placeholder for a target gene sequence, a lacZ cassette (approx. 600 bp) is integrated. Upon digestion, with BsaI the lacZ cassette is released and a fitting open reading can be integrated in frame (e.g. Signal Peptide-Gene-C-terminal Tag) into the plasmid, thereby reconstituting a fully functional expression plasmid.
For the stable integration of transcription units into the *P. pastoris,* a third universal integrative plasmid (pPAP003) was designed. This shuttle plasmid was constructed, which can be propagated in *E.coli* (Kanamycin Resistance) as well as *P. pastoris* (Hygromycin B resistance). As placeholder for a target transcription unit a lacZ cassette (approx. 600 bp) is integrated, enabling blue/white selection of transformants based on the conversion of X-Gal. Upon digestion, with BbsI the lacZ cassette is released and a fitting transcription unit (Promoter- ORF-Terminator) can be integrated (derived from respective pPAP001 and pPAP002 episomal plasmids as donors) into the plasmid, thereby reconstituting a fully functional integration plasmid.
Several parts (GAP promoter, GAP terminator, AOX integration marker and Hygromycin resistance marker) of the constructed plasmids were PCR amplified and derived from a previously introduced Golden Gate based *P. pastoris* assembly system, coined GoldenPiCS[3]. These parts are particularly suitable since they have been already domesticated regarding internal BsaI and BbsI recognition sequences. The occurrence of these recognition sites within the plasmid backbone would greatly diminish Golden Gate Assembly efficiency.

**Golden Gate Cloning of Level 0 standard parts.** All genetic parts were cloned as individual Level 0 standard modules into the universal Level 0 acceptor plasmid pAGM9121 (Spectinomycin$^{R}$). Therefore three functional units were pre-defined: a) signal peptide (contains start codon); b) gene (lacking start and stop codon) and c) C-terminal Protein-tag (contains stop codon). The chosen 4 bp sticky overhangs that are released upon Type II s enzyme treatment (BsaI and BbsI) and guide subsequently a correct reassembly were chosen accordingly to the nomenclature of gene assembly as described within the ModularCloning (MoClo) system[4]. An overview of the reassembly concept is provided in Supplementary Fig. 1.

For the cloning of the individual modules suitable Oligonucleotides were designed to allow for cloning into pAGM9121. Primers followed the general scheme: **5´** *BbsI recognition site – two spacer nucleotides – matching 4 bp overhang specifying module type* (signal peptide, gene or C-terminal tag) – *template binding sequence*. Fragments were amplified by PCR from a suitable template sequence or generated by hybridisation of two complementary Oligonucleotides (in the case of many short signal peptides). PCR and hybridisation products were analysed as small aliquot by Agarose gel electrophoresis for occurrence of the expected size and the remaining sample subsequently recovered and purified using a NucleoSpin® Gel and PCR Clean-up Kit (Macherey-Nagel, Düren, DE).

Golden Gate reactions were performed in a total volume of 15 µL. The final reaction volume contained 1-fold concentrated T4 ligase buffer (Promega, Madison, US). Prepared reaction mixtures containing ligase buffer, acceptor plasmid (20 fmol) and the corresponding insert (20 fmol) was adjusted to 13.5 µL with dd$H_2O$. In a final step, the corresponding enzymes were quickly added. First, a volume of 0.5 µL of the respective restriction enzyme BbsI (5 units/µL) was added and then 1 µL (1–3 units/µL) of T4 ligase. Golden Gate reactions were performed for 3 hours (37 °C) and concluded by an additional enzyme inactivation step (80 °C; 20 min).

The whole Golden Gate reaction volume was used to transform chemically competent *E.coli* DH10B cells. After heat shock transformation and recovery the mixture was plated in different quantities on selective LB Agar plates (50 µg × $mL^{-1}$ X-Gal; 100 µg × $mL^{-1}$ Spectinomycin; 150 µM IPTG). Based on the occurrence of the lacZ selection marker one can easily distinguish between white colonies (recombined plasmid) and empty plasmid (blue). In general, the described protocol led to several thousand recombinant colonies with a nearly absolute proportion (> 99 %) of recombined, white colonies. Single colonies were checked for correct insert sizes by means of colony PCR (pAGM9121 sequencing primer). Positively identified clones were inoculated into 4 mL of TB-Medium (100 µg × $mL^{-1}$ Spectinomycin) and corresponding plasmid DNA prepared (NucleoSpin Plasmid Kit (Macherey-Nagel, Düren, DE)). After verification of the correct, intended insert sequence by Sanger Sequencing (Eurofins Genomics, Ebersbach, DE) respective plasmids were included for further use within the Golden Gate cloning approaches.

**Golden Gate Cloning of Expression Plasmids.** The expression plasmids (*S. cerevisiae*: pAGT572_Nemo and pAGT572_Nemo 2.0; *P. pastoris*: pPAP001 and pPAP002) were used as respective acceptor plasmid for the assembly of the individual tripartite open reading frames (**5´** Signal Peptide-Gene-C-terminal Tag **3´**). The individual parts were thereby derived as parts from standard level 0 plasmids (pAGM9121 backbone), which can be released from the pAGM9121 backbone upon BsaI restriction digest.

Golden Gate reactions were performed in a total volume of 15 µL. The final reaction volume contained 1-fold concentrated T4 ligase buffer. Prepared reaction mixtures containing ligase buffer, the acceptor plasmid (20 fmol) and the corresponding inserts as level 0 modules (Signal Peptide, Gene, C-terminal Tag) were added to 20 fmol each and the overall volume adjusted to 13.5 µL with dd$H_2O$. In the case of a signal peptide, shuffling approaches 17 different pAGM9121- Signal Peptide combinations were added in equimolar ratios (1.2 fmol each). In a final step, the corresponding enzymes were quickly

added. First, a volume of 0.5 µL of the restriction enzyme BsaI (10 units/µL) was added and then 1 µL (1–3 units/µL) of T4 ligase. Golden Gate reactions were performed using a temperature cycling program (50x passes) between 37 °C (2 min) and 16 °C (5 min) and concluded by an additional enzyme inactivation step (80 °C; 20 min).
The whole Golden Gate reaction volume was used to transform chemically competent *E.coli* DH10B cells. After heat shock transformation (90 seconds at 42 °C) and recovery the mixture (approx. 320 µL) was split into two fractions, 50 µL were plated on selective LB Agar plates (50 µg × $mL^{-1}$ X-Gal; 100 µg × $mL^{-1}$ Ampicillin; 150 µM IPTG) and the remaining volume used to directly inoculate 4 mL TB Medium (100 µg × $mL^{-1}$ Ampicillin) to preserve the genetic diversity of the shuffling library. The following day the success of the Golden Gate reaction was evaluated based on the performed blue/white screening, discriminating the empty plasmid (lacZ; blue) from recombined, white colonies. In general, the described protocol for ORF assembly and signal peptide shuffling as special case led to several hundred recombinant colonies with a high proportion (> 90 %) of recombined, white colonies. In the case of single defined, "unshuffled" constructs single colonies were checked for correct insert sizes by means of colony PCR (using respective plasmid sequencing primer). Positively identified clones were inoculated into 4 mL of TB-Medium (100 µg × $mL^{-1}$ Ampicillin) and corresponding plasmid DNA prepared (NucleoSpin Plasmid Kit (Macherey-Nagel, Düren, DE)). In the case of shuffled signal peptide constructs, plasmid DNA was prepared as a library by direct inoculation of the transformation mixture into the liquid culture and subsequent DNA isolation (see above).

**Plasmid transformation into *S. cerevisiae*.** Respective single plasmids or plasmid mixtures (pAGT572_Nemo or pAGT572_Nemo 2.0 backbone) were used to transform chemically competent *S. cerevisiae* cells (INVSc1 strain) by polyethylene glycol/lithium acetate transformation. INVSc1 cells were prepared and stored at – 80 °C in transformation buffer (15 % (v/v) glycerol; 100 mM lithium acetate; 500 µM EDTA; 5 mM Tris-HCl pH 7.4) as 60 µL aliquots until usage.
For transformation, an amount of 100 ng of the plasmid preparation was added to 10 µL of lachssperm DNA (10 mg/mL; Sigma Aldrich, Hamburg, DE) and mixed. This mixture was then added to a thawed aliquot of INVSc1 cells on ice. 600 µL of transformation buffer (40 % (v/v) poly ethylene glycol 4000; 100 mM lithium acetate; 1 mM EDTA; 10 mM Tris-HCl pH 7.4) were added and the cells incubated under rigid shaking (30 °C; 850 rpm) for 30 min. Afterwards, 70 µL of pure DMSO were added and the cells incubated for a further 15 min at 42 °C without shaking. Finally, the cells were precipitated by short centrifugation, the supernatant discarded, and the cell pellet resuspended in 350 µL sterile dd$H_2O$. Different volumes were plated on Synthetic Complement (SC) Drop Out plates supplemented with 2 % (w/v) glucose as carbon source and lacking Uracil as an auxotrophic selection marker. Plates were incubated for at least 48 hours at 30 °C till clearly background distinguishable white colonies appeared.

**Microtiter Plate cultivation of *S. cerevisiae*.** For peroxygenase production in microtiter plate format specialised 96 half deep well plates were utilised. The model type CR1496c was purchased from EnzyScreen (Heemstede, NL) and plates were covered with fitting CR1396b Sandwich cover for cultivation. Plates and covers were flushed before every experiment thoroughly with 70 % ethanol and air-dried under a sterile bench until usage. In each cavity, 220 µL of minimal expression medium were filled and inoculated with single, clearly separated yeast colonies using sterile toothpicks. The minimal selective expression medium (1x concentrated SC Drop stock solution lacking uracil; 2 % (w/v) galactose; 71 mM potassium phosphate buffer pH 6.0; 3.2 mM magnesium sulfate; 3.3 % (v/v) ethanol;

50 mg/L hemoglobin; 25 mg/L chloramphenicol) was freshly prepared out of sterile stock solutions immediately before each experiment, mixed and added to the cavities.
After inoculation of the wells the plates were covered, mounted on CR1800 cover clamps (EnzyScreen) and incubated in a Minitron shaking incubator (Infors, Bottmingen, SUI) for 72 h (30 °C; 230 rpm). After cultivation, the cells were separated from the peroxygenase containing supernatant by centrifugation (3400 rpm; 50 min; 4 °C).

**Shake flask cultivation *S. cerevisiae*.** For larger production volumes in shake flask cultures, alterations were made. As preculture 50 mL of SC Drop out selection media (+ 2 % (w/v) Raffinose as non-repressible carbon source and 25 mg/L chloramphenicol) was inoculated with one single colony derived from a selection plate (SC Drop; -Uracil) and grown for 48 h (30 °C; 160 rpm; 80 % humidity). This incubation typically resulted in a final $OD_{600nm}$ of approx. 12 to 13. The main expression culture was inoculated with a starting optical density of 0.3. For large scale peroxygenase production rich non-selective expression medium (20 g/L peptone; 10 g/L yeast extract; 2 % (w/v) galactose; 71 mM potassium phosphate buffer pH 6.0; 3.2 mM magnesium sulfate; 3.3 % (v/v) ethanol; 25 mg/L chloramphenicol) was utilised. Cultivation was performed in 2.5 L Ultra yield flasks (Thomson Instrument, Oceanside, US) in a final culture volume of 500 mL per flask after sealing the flask with breathable Aeraseal tape (Sigma Aldrich, Hamburg, DE) allowing for gas exchange. The main cultures were incubated for further 72 h (25 °C; 110 rpm; 80 % humidity). After cultivation, the cells were separated from the peroxygenase containing supernatant by centrifugation (4300 rpm; 35 min; 4 °C).

**Supernatant ultrafiltration and Protein purification.** The previously prepared supernatant was concentrated approx. 20-fold by means of ultrafiltration. Therefore a Sartocon Slice 200 membrane holder (Sartorius, Göttingen, DE) was equipped with a Sartocon Slice 200 ECO Hydrosart Membrane (10 kDa nominal cut-off; Sartorius) within a self-made flow setup. The flow system for ultrafiltration was operated by an EasyLoad peristaltic pump (VWR International, Darmstadt, DE).
In a first step, the cleared supernatant (1 L) was concentrated approx. 10 fold to a volume of 100 mL and 900 mL of purification binding buffer (100 mM Tris-HCl pH 8.0, 150 mM NaCl) were added as a buffer exchange step. This sample was then concentrated approx. 20-fold to a final volume of 50 mL. Protein purification was implemented utilising the C-terminal attached double Strep II Tag (WSHPQFEK), coined TwinStrep® (Iba Lifesciences, Göttingen, DE). As column material, Strep-Tactin®XT Superflow® columns (1 mL or 5 mL; Iba Lifesciences) were chosen and the flow system operated by an EasyLoad peristaltic pump (VWR). In a first step, the column was equilibrated with 5 column volumes (CVs) binding buffer. The concentrated sample (50 mL) was filter sterilised (0.2 µm syringe filter) and applied to the column with an approximate flow rate of 1 mL/min. After application, the column was washed with 6 CVs binding buffer. Elution was performed based on binding competition with biotin, therefore approx. 2 CV of elution buffer (100 mM Tris-HCl pH 8.0, 150 mM NaCl; 50 mM biotin) were applied to the column. The pooled elution fraction was then dialysed overnight (4 °C) against 5 L of storage buffer (100 mM potassium phosphate pH 7.0) using ZelluTrans dialysis tubing (6-8 kDa nominal cut-off; Carl Roth) and the recovered, dialysed sample stored at 4 °C till further use.

**Plasmid transformation into *P. pastoris*.** Respective single plasmids or plasmid mixtures (pPAP001 or pPAP002 backbone) were used to transform electrocompetent *P. pastoris* cells (X-33 strain) by means of electroporation. Electrocompetent X-33 cells were prepared according to a condensed transformation protocol for *P. pastoris*[5]. Cells were stored in BEDS solution (10 mM bicine-NaOH pH 8.3, 3 % (v/v) ethylene glycol, 5 % (v/v) DMSO and 1 M sorbitol) as 60 µL aliquots (-80 °C) till further use.

For the transformation of episomal plasmids (pPAP001 and pPAP002) 20 ng of the circular plasmid (predilution in dd$H_2O$ to a 20 ng/µL stock solution) were added to one aliquot of on ice thawed competent X-33 cells. The cell-plasmid mix was then transferred to an electroporation cuvette (2 mm gap), and the cuvette cooled for 10 minutes on ice prior to the transformation. Electroporation was performed using a Micropulser Device (Bio-Rad, Hercules, US) and using manual implemented, standardised settings (1.5 kV, 1 pulse) for all transformation setups, leading to a general pulse interval of 5.4 to 5.7 ms. Immediately after electroporation cells were recovered in 1 mL of ice-cold YPD-Sorbitol solution (10 g/L peptone, 5 g/L yeast extract, 500 mM sorbitol), transferred to a new reaction tube and incubated for one hour under rigid shaking (30 °C, 900 rpm) in a Thermomix device (Eppendorf, Hamburg, DE). After incubation, cells were precipitated by centrifugation (5.700 rpm, 5 min). The supernatant was discarded and the cells resuspended in 200 µL of fresh YPD medium. 100 µL of the suspension was then plated on selective YPD Agar plates supplemented with 150 µg/mL Hygromycin B. Plates were incubated at 30 °C for at least 48 h till clearly visible colonies appeared. In general, the described setup led to the occurrence of several hundred colonies per plate.

For the transformation of integrative plasmids (pPAP003 backbone) the setup was slightly modified since, in this case, linearised plasmids are used for transformation. Therefore previously prepared circular plasmid DNA was digested with AscI (Isoschizomer: SgsI). 2.5 µg of the respective plasmid DNA were mixed with 3 µL of 10x fold FastDigest Buffer, the volume adjusted to 29.5 µL using dd$H_2O$ and in the last step, 0.5 µL of FastDigest SgsI added. Digestion was performed overnight (16 h, 37 °C) and terminated by an enzyme inactivation step (20 min, 80 °C). Linearised plasmid DNA was then subsequently prepared according to the manufacturer instruction using a Nucleospin ® Gel and PCR clean up Kit (Macherey-Nagel, Düren, DE). The transformation of *P. pastoris* was performed in a congruent manner as described before, with the exception of using 100 ng linearised plasmid for transformation, since the overall transformation efficiency is substantially reduced in comparison to the transformation of the circular, episomal plasmid.

**Microtiter Plate cultivation expression in *P. pastoris*.** For peroxygenase production in microtiter plate format specialised 96 half deep well plates were utilised. The model type CR1496c was purchased from EnzyScreen (Heemstede, NL) and plates were covered with fitting CR1396b Sandwich cover for cultivation. Plates and covers were flushed before every experiment thoroughly with 70 % ethanol and air-dried under a sterile bench until usage. Each cavity was filled with 220 µL of buffered complex medium (BM) and inoculated with single, clearly separated yeast colonies using sterile toothpicks. Basic BM (20 g/L peptone; 10 g/L yeast extract; 100 mM potassium phosphate buffer pH 6.0; 1x YNB (3.4 g/L yeast nitrogen base without amino acids; 10 g/L ammonium sulfate); 400 µg/L biotin; 3.2 mM magnesium sulfate; 25 mg/L chloramphenicol; 50 mg/L haemoglobin; 150 mg/L Hygromycin B) was freshly prepared out of sterile stock solutions immediately before each experiment, mixed and added to the cavities.

Depending on the type of utilised promoter (pPAP001: GAP and pPAP002: CAT1), the BM medium was supplemented with different carbon sources for cultivation and induction, respectively. pPAP001 constructs were cultivated utilising 1.5 % (w/v) of glycerol or glucose as sole carbon source. In the case of the methanol inducible CAT1 promoter, a mixed feed strategy was employed combining 0.5 % (w/v) of glycerol with 1.5 % (v/v) methanol.

After inoculation of the wells the plates were covered, mounted on CR1800 cover clamps (EnzyScreen) and incubated in a Minitron shaking incubator (Infors, Bottmingen, SUI) for 72 h (30 °C; 230 rpm). After cultivation, the cells were separated from the peroxygenase containing supernatant by centrifugation (3400 rpm; 50 min; 4 °C).

**Shake flask cultivation *P. pastoris*.** For the large scale protein production using shake flasks genomically integrated single constructs (pPAP 003 backbone; integration into chromosomal 3´ region of *P. pastoris* AOX1 gene) were chosen. These constructs were previously identified by screening 4 different colonies per individual construct (Signal peptide – gene combination) within an MTP screening setup and choosing a respective production strain based on a high as possible, clearly distinguishable NBD conversion in comparison to the background control (pPAP003 empty plasmid). Precultures were prepared in 50 mL YPD medium (+ 25 mg/L chloramphenicol) and cultivated for 48 h (30 °C; 160 rpm; 80 % humidity), typically resulting in a final $OD_{600nm}$ of approx. 17 to 19. The main expression culture was inoculated with a starting optical density of 0.3. For large scale peroxygenase production BM based expression media (20 g/L peptone; 10 g/L yeast extract; 100 mM potassium phosphate buffer pH 6.0; 1x YNB (3.4 g/L yeast nitrogen base without amino acids; 10 g/L ammonium sulfate); 400 µg/L biotin; 3.2 mM magnesium sulfate; 25 mg/L chloramphenicol) was utilised. Cultivation was performed in 2,5 L Ultra yield flasks (Thomson Instrument, Oceanside, US) in a final culture volume of 500 mL per flask after sealing the flask with breathable Aeraseal tape (Sigma Aldrich, Hamburg, DE) allowing for gas exchange. The main cultures were incubated for further 72 h (25 °C; 110 rpm; 80 % humidity).
In the case of constitutively expressing GAP constructs 2 % (w/v) Glucose was added (BMG media) as a carbon source for Pichia growth. In the case of the methanol inducible CAT1 promoter a two-phase feeding was applied, firstly inoculating the cells into BM medium (see above) supplemented with 0.5 % (w/v) glycerol as carbon source. 24 h and 48 h after inoculation 0.8 % (v/v) of methanol were added as an inducer of the CAT1 promoter. After cultivation, the cells were separated from the peroxygenase containing supernatant by centrifugation (4300 rpm; 35 min; 4 °C). Subsequent ultrafiltration of the supernatant and protein purification were performed as described before.

**Plasmid preparation of episomal plasmids from yeast.** Yeast plasmids of identified clones were recovered by means of digestive Zymolase cell treatment and alkaline cell lysis. Therefore clones were inoculated and cultivated for 48 h (30 °C; 250 rpm) in 4 mL of selection medium, in case of *S. cerevisiae* SC Drop out medium (-Uracil; 2 % (w/v) Glucose) was used and in the case of *P. pastoris* single colonies were inoculated into 4 mL of YPD (+150 µg/mL Hygromycin) to preserve the selection pressure. After cultivation cells were pelleted by centrifugation and 1 mL of washing buffer (10 mM EDTA NaOH; pH 8.0) added and the pellet resuspended by light vortexing. Cells were subsequently pelleted (5000 × g; 10 min) and the supernatant discarded. Afterwards, cells were resuspended by light vortexing in 600 µL of Sorbitol Buffer (1.2 M sorbitol, 10 mM $CaCl_2$, 100 mM Tris-HCl pH 7.5, 35 mM β-mercaptoethanol) and 200 units of Zymolase (Sigma Aldrich, Hamburg, DE) added followed by an incubation step for 45 min (30 °C; 800 rpm) to facilitate yeast cell wall digestion. After incubation cells were pelleted by centrifugation (2000 × g; 10 min) the supernatant discarded and the plasmid preparation started with an alkaline lysis step following the manufacturer's instructions (NucleoSpin Plasmid Kit, Macherey Nagel). In the final step, the yeast-derived episomal plasmids were eluted in 25 µL elution buffer (5 mM Tris-HCl pH 8.5), and the whole eluate used to transform one aliquot of *E.coli* DH10B

(transformation as described above), plating the whole transformation mix on a selective LB-Agar plate (50 µg × mL$^{-1}$ X-Gal; 100 µg × mL$^{-1}$ Ampicillin; 150 µM IPTG).
On the following day, single colonies were picked, inoculated into 4 mL of TB medium (100 µg × mL$^{-1}$ Ampicillin), plasmid prepared (NucleoSpin Plasmid Kit) and sent for Sanger Sequencing to elucidate the respective sequence of the open reading frame (Eurofins Genomics).

**Thermostability measurements.** Thermostability measurements of the purified enzymes were performed by Differential Scanning fluorimetry (DSF) on a Prometheus NT.48 nanoDSF instrument (NanoTemper Technologies GmbH, München, DE) in storage buffer (100 mM Tris-HCl pH 7.0). Approximately 10 µL of sample volume were loaded into a Prometheus NT.48 High Sensitivity Capillary (NanoTemper Technologies GmbH). Protein unfolding was subsequently monitored by following the ratio of intrinsic protein tyrosine and tryptophan fluorescence at 350 nm to 330 nm over time, increasing the temperature from 20 °C to 95 °C with a heating ramp of 1 °C per minute. The melting temperature corresponds to the maximum of the first derivative of the 350/330 nm ratio. All measurements were performed at least in triplicates.

**Split-GFP assay.** Protein normalisation was performed employing the principle of a split GFP normalisation assay as described by Santos-Aberturas *et al.* [6] with slight modifications. The GFP fluorescence complementation fragment sfGFP 1-10 was cloned into the Golden Mutagenesis plasmid pAGM22082_cRed[7] for T7 promoter controlled expression in *E.coli* (BL 21 DE3 strain). The sfGFP 1-10 fragment was then prepared as an inclusion body preparation according to the previous reports[6].
For measurement a 96 well Nunc MaxiSorp Fluorescence plate (ThermoFisherScientific, Waltham, US) was first blocked with 180 µL of BSA blocking buffer (100 mM Tris-HCl pH 7.4, 100 mM NaCl, 10 % (v/v) glycerol, 0.5 % (w/v) BSA) per well incubating for 25 min on an orbital shaker with light shaking. The blocking solution was discarded and 20 µL of the yeast media supernatant (*S. cerevisiae* or *P. pastoris*) derived from the peroxygenase expression plates added using a multichannel pipet. A 10 mL aliquot of the sfGFP 1-10 complementation fragment (storage: -80 °C) was quickly thawed in a water bath and diluted 1x fold into ice-cold TNG buffer (100 mM Tris-HCl pH 7.4, 100 mM NaCl, 10 % (v/v) glycerol) and 180 µL of this screening solution added to each well.
Immediate fluorescence values (GFP fluorophore: excitation wavelength: 485 nm; emission wavelength: 535 nm; top read mode) were measured using a 96 well plate fluorescence reader Spark 10M (TECAN, Grödig, AT), setting an empty plasmid control well as 10 % of the overall signal intensity (well calculated gain). After storage of the plate for at least one up to three nights (at 4 °C) final fluorescence values were measured in a comparable manner. Protein quantities were then normalised based on the relative fluorescence increase of each respective well (differential values) and in comparison to the empty plasmid backbone.

**DMP assay.** The use of 2,6-Dimethoxyphenol (DMP) as a suitable microtiter plate substrate for the measurement of peroxygenase catalysed conversion to the colourimetric product ceruglinone has been described before[8]. The described conditions have been adapted with slight modifications. In brief, 20 µL of peroxygenase containing supernatant (from 96 well plate setup) were transferred to a transparent polypropylene 96 well screening plate (Greiner Bio-One, Kremsmünster, AT) and 180 µL of screening solution (final: 100 mM potassium phosphate pH 6.0; 3 mM 2,6-Dimethoxyphenol; 1 mM hydrogen peroxide) added. Absorption values (λ: 469 nm) of each well were immediately measured after addition in a kinetic mode (measurement interval: 30 s) over a duration of 5 minutes utilising the

96 well microtiter plate reader SpectraMax M5 (Molecular Devices, San José, US). Slope values of absorption increase corresponding to ceruglinone formation were obtained, paying special attention to the linearity of the observed slope to obtain reliable relative DMP conversion values for comparison of the respective wells.

**NBD assay.** The use of 5-nitro-1,3-benzodioxole (NBD) as a suitable microtiter plate substrate for the measurement of peroxygenase catalysed conversion to the colourimetric product 4-Nitrocatechol has been described before[9,10]. The described conditions have been adapted with slight modifications. In brief 20 µL of peroxygenase containing supernatant (from 96 well plate setup) were transferred to a transparent polypropylene 96 well screening plate (Greiner Bio-One, Kremsmünster, AT) and 180 µL of screening solution (final: 100 mM potassium phosphate pH 6.0; 1 mM NBD; 1 mM hydrogen peroxide; 12 % (v/v) acetonitrile) added. Absorption values (λ: 425 nm) of each well were immediately measured after addition in a kinetic mode (measurement interval: 30 s) over a duration of 5 minutes utilising the 96 well microtiter plate reader SpectraMax M5 (Molecular Devices, San José, US). Slope values of absorption increase corresponding to 4-nitrocatechol formation were obtained, paying special attention to the linearity of the observed slope to obtain reliable relative NBD conversion values for comparison of the respective wells.

**Resting-state absorption and heme CO complex measurements.** The pooled and dialysed elution fractions (100 mM potassium phosphate pH 7.0) were subsequently used to record absorption spectra of the respective enzymes (*Mro*UPO, *Cgl*UPO, *Mth*UPO, *Tte*UPO) in their native, resting state (ferric iron; $Fe^{3+}$). For all measurements a QS High precision Quartz Cell cuvette (Hellma Analytics, Müllheim, DE) with a path length of 10 mm was used. Spectra were recorded on a Biospectrometer Basic device (Eppendorf, Hamburg, DE) in the spectral range from 250 to 600 nm (interval: 1 nm) and subtracting the utilised storage buffer (100 mM potassium phosphate pH 7.0) as previous blank measurement. Heme carbon dioxide spectra (CO assay) were recorded after reducing the heme iron to its ferrous form ($Fe^{2+}$). Therefore a spatula tip of sodium dithionite as the reducing agent was added to 1 mL of a respective enzyme fraction (see above) and mixed thoroughly till complete dissolution. This sample was immediately flushed with a constant carbon dioxide flow for 2 min (approx. 1 bubble/sec) to obtain the thiolate-heme carbon dioxide complex. The sample was immediately transferred to a cuvette and absorption measured as described above.
The CO assay was also employed for the measurement of Peroxygenase concentrations in the concentrated *P. pastoris* supernatant obtained after ultrafiltration. In this case, the supernatant was 10-fold diluted with potassium phosphate buffer (100 mM, pH 7.0). A spatula tip of sodium dithionite was then added to 2 mL of the diluted supernatant sample. After dividing the respective sample into two parts of 1 mL, one part was treated with carbon monoxide for 2 minutes as described above, and the CO untreated sample is used as a blank reference. Absorption measurements were performed by UV/Vis spectroscopy using a JASCO V-770 Spectrophotometer (JASCO Deutschland GmbH, Pfungstadt) with the following measurement parameters: photometer method: absorption, UV-VIS bandwidth: 1 nm, UV-VIS response: 0.96 sec, wavelength: 400-500 nm, data interval: 1 nm, scan speed: 100 nm/min. The CO absorption maximum was measured at 444 nm, and the reference absorption wavelength was measured at 490 nm. For calculation, an extinction coefficient of 91000 $M^{-1}$ $cm^{-1}$ was used, which appears to be generally valid for most P450 enzymes according to literature[11]. The enzyme concentration in the supernatant was then calculated using the formula:

$$c\,[\mu M] = \text{dilution factor} \times \frac{A_{444\,nm} - A_{490\,nm}}{0.091\ \mu M^{-1} cm^{-1}}$$

**pH range of NBD conversion.** pH dependency of NBD conversion of the respective enzymes was investigated using different buffer system in the range between pH 2.0 and 11.0 (even numbers only). Each buffer was prepared as a 100 mM stock solution, potassium phosphate buffer was used for the pH values 2.0, 7.0 and 8.0. Sodium citrate was used in the range of pH 3.0 to 6.0 and Tris-HCl was used in the range of pH 8.0 to 11.0. Purified enzyme solutions (100 mM potassium phosphate pH 7.0) were diluted 10 to 20x fold in $ddH_20$ prior to the measurements leading to weakly buffered solutions as screening samples. The NBD assay was then performed as described before, mixing 20 µL of the enzyme dilution with 180 µL screening solution (87 mM corresponding buffer pH x; 500 µM NBD; 1 mM $H_2O_2$). All samples were measured as three biological replicates.
Since the molar extinction coefficient of the corresponding detected product 4-nitrocatechol is strongly pH-dependent a normalisation of the obtained NBD conversion slopes was performed. Therefore, the product 4-Nitrocatechol was prepared as 10 mM stock solution dissolved in acetonitrile and diluted into 990 µl of the corresponding screening buffer (final 4-Nitrocatechol concentration: 10 µM) and after 5 minutes an absorption spectra in the interval of 400 to 600 nm (Biospectrometer Basic device) recorded. Calculation of the correction factor of the respective samples (pH 2.0 to pH 11.0) was then performed regarding the utilised measurement wavelength of 425 nm. Finally, in consideration of the obtained pH correction factor, individual activity values derived from the respective measured absorption values were calculated.

**Protein concentration determination and purification yield.** Protein concentrations of the respective protein samples were determined after dialysis of the elution fractions (storage buffer: 100 mM potassium phosphate pH 7.0). In this regard, the colourimetric BCA assay was utilised, employing a Pierce™ BCA Protein Assay Kit (ThermoFisherScientific, Waltham, US) following the instructions of the manufacturer. Samples were measured in biological triplicates (25 µL of a previously diluted sample) and concentrations calculated based on a previously performed calibration curve using BSA (0 - 1000 µg/mL) as reference protein.
To determine the overall yield of enzyme production per litre of culture volume, the determined concentration in the elution fraction was extrapolated to the overall NBD activity of the sample after ultrafiltration (column load). This calculation is based on the fact that NBD is a highly specific substrate for peroxygenase activity, comparable background samples processed in a similar manner but using empty plasmid controls (pAGT572_Nemo 2.0 and pPAP003) did not show any measurable conversion of NBD at all.
Therefore, samples of every purification step (load, flow-through, wash and elution fraction) were collected, and NBD conversion rates of the respective fractions measured immediately after purification. In the case of non-complete binding of the enzyme fraction to the affinity column (remaining NBD activity in flow-through fractions) this remaining non-bound enzyme amount was taken into consideration for calculation for the overall volumetric production yield. Therefore, the via BCA assay determined protein concentration of the elution fraction was extrapolated to the activity of the respective non-bond fraction, assuming a comparable specific enzyme activity for NBD conversion and considering the volumes of the respective fractions, leading to an approximate enzyme titre per litre of shake flask culture.

**SDS Gel analysis and PNGase F treatment.** Obtained elution fractions of the respective UPO enzymes were analysed for the apparent molecular weight and overall purity after the performed one step TwinStrep purification by means of SDS PAGE. Therefore, samples of the column load (after ultrafiltration; see above), elution fractions after dialysis and deglycosylated elution fraction samples were analysed on self-casted SDS PAGE (10 or 12 % of acrylamide) utilising a Bio-Rad (Hercules, US) Mini-Protean® Gel electrophoresis System. For the purpose of molecular weight determination, a defined PageRuler Prestained Protein Ladder (ThermoFisherScientific, Waltham, US) was included, covering a MW range between 10 and 180 kDa. Proteins were visualised using a colloidal Coomassie G-250 staining solution.

To obtain N-type deglycosylated protein samples elution fractions were enzymatically treated with Peptide-N-Glycosidase F (PNGaseF) from *Flavobacterium meningosepticum*, which is capable of cleaving Asparagine linked high mannose type glycan structures as typically occurring in *P. pastoris* and *S. cerevisiae* derived glycosylation patterns. Therefore, 45 µL of a respective elution fraction was mixed with 5 µL of denaturing Buffer (final 0.5 % SDS; 40 mM DTT) and denatured for 10 minutes (100 °C). After a cooling step to room temperature 6 µL of NP-40 solution (final: 1 %) and 6 µL of GlycoBuffer2 (500 mM sodium phosphate; pH 7.5) were added and the solution thoroughly mixed. Finally, 1 µL of PNGaseF (New England Biolabs, Ipswich, US) was added and the sample incubated under light shaking (37 °C) for 3 hours. After incubation, the sample was prepared for further analysis by adding 5x fold SDS sample buffer and subsequent SDS PAGE analysis executed as described before.

In the case of native deglycosylation, 90 µL of enzyme sample were mixed with 10 µL of GlycoBuffer2 (500 mM Sodium Phosphate; pH 7.5) and 1 µL of PNGaseF added. The mixture was incubated at 37 °C in a thermal PCR cycler (24 h) and subsequently analysed for UPO activity in comparison with an equally treated sample (without PNGAseF addition) by means of the NBD assay (see above).

**Protein identification by MS.** Protein samples after protein purification (in 100 mM Tris-HCl pH 8.0, 150 mM NaCl; 50 mM biotin) were enzymatically digested with trypsin and desalted according to[12]. The resulting peptides were separated using C18 reverse-phase chemistry employing a pre-column (EASY column SC001, length 2 cm, ID 100 µm, particle size 5 µm) in line with an EASY column SC200 with a length of 10 cm, an inner diameter (ID) of 75 µm and a particle size of 3 µm on an EASY-nLC II (all from Thermo Fisher Scientific). Peptides were eluted into a Nanospray Flex ion source (Thermo Fisher Scientific) with a 60 min gradient increasing from 5 % to 40 % acetonitrile in dd$H_2O$ with a flow rate of 300 nL/min and electrosprayed into an Orbitrap Velos Pro mass spectrometer (Thermo Fisher Scientific). The source voltage was set to 1.9 kV, the S Lens RF level to 50%. The delta multipole offset was -7.00. The AGC target value was set to 1e06 and the maximum injection time (max IT) to 500 ms in the Orbitrap. The parameters were set to 3e04 and 50 ms in the LTQ with an isolation width of 2 Da for precursor isolation and MS/MS scanning. Peptides were analysed using a Top 10 DDA scan strategy employing HCD fragmentation with stepped collision energies (normalised collision energy 40, 3 collision energy steps, width 15).

MS/MS spectra were used to search the TAIR10 database (ftp://ftp.arabidopsis.org, 35394 sequences, 14486974 residues) amended with target protein sequences with the Mascot software v.2.5 linked to Proteome Discoverer v.2.1. The enzyme specificity was set to trypsin, and two missed cleavages were tolerated. Carbamidomethylation of cysteine was set as a fixed modification and oxidation of methionine. Searches were performed with enzyme specificity set to trypsin and semi-trypsin to identify truncated protein N-termini. The precursor tolerance was set to 7 ppm, and the product ion mass tolerance was set to 0.8 Da. A decoy database search was performed to determine the peptide spectral match (PSM) and peptide identification false discovery rates (FDR). PSM, peptide and protein identifications surpassing respective FDR thresholds of q<0.01 were accepted.

**GC-MS calibration curves.** For product quantification, calibration curves were created as depicted in Supplementary Fig. 13. The quantification was achieved in Scan mode (*N*-(2-hydroxy-2-phenylethyl) phthalimide) or SIM mode (all other substrates) whereby each concentration data point was measured as triplicates and correlated to an internal standard (IS). The final product concentration was adjusted in 100 mM KPi buffer (pH 7.0) with the corresponding stock solutions in acetone yielding to 5 % (v/v) final co-solvent proportion in the buffer system. Extraction was achieved adding 650 µL (*N*-(2-hydroxy-2-phenylethyl)phthalimide) or 400 µL (all other substrates) of ethyl acetate (containing 1 mM of the internal standard) and vortexing for 30 s, followed by brief centrifugation (1 min, 8400 rpm). The organic layer was utilised for GC-MS measurements applying the corresponding temperature program as listed in Supplementary Table 5. For enantiomeric product identification corresponding R-enantiomer standards were utilised (Supplementary Fig. 14).

**UPO Bioconversions for subsequent GC-MS and chiral HPLC analytics.** For the tested hydroxylation (naphthalene, phenylethane, -propane, butane and pentane) and epoxidation (styrene) reactions, purified UPOs enzyme samples (stored in 100 mM potassium phosphate; pH 7.0) produced in *S. cerevisiae* were used. Respective reactions (total volume: 400 µl) were performed as biological triplicates in 100 mM potassium phosphate (pH 7.0) containing 100 nM of UPO, 1 mM of the respective substrate and 500 µM $H_2O_2$. The substrate was prior dissolved in pure acetone (20 mM stock solution) yielding a 5 % (v/v) co-solvent ratio in the final reaction mixture. Reactions were performed for 60 minutes (25 °C, 850 rpm) and subsequently quenched by the addition of 400 µl ethyl acetate (internal standard: 1 mM ethyl benzoate). Extraction was accomplished by 30 s of vigorous vortexing, followed by brief centrifugation (1 min, 8400 rpm). The organic layer was then utilised for respective GC-MS measurements as described in Supplementary Table 5.

In the case of the hydroxylation reaction of *N*-phthaloyl-phenylethyl amine, purified UPOs enzyme samples (stored in 100 mM potassium phosphate, pH 7.0.) previously produced in *P. pastoris* were used. Reactions (total volume: 500 µL) were performed as biological triplicates in 100 mM potassium phosphate (pH 7.0) containing 100 nM of the respective UPO, 250 µM of the substrate *N*-phthaloyl-phenylethyl amine and 250 µM $H_2O_2$. The substrate was prior dissolved in pure acetone (5 mM stock solution) yielding a 5% (v/v) co-solvent ratio in the final reaction mixture. Reactions were performed for 60 minutes (30 °C, 850 rpm) and subsequently quenched by the addition of 650 µL ethyl acetate (internal standard: 1 mM ethyl benzoate). Extraction was accomplished by 30 s of vigorous vortexing, followed by brief centrifugation (1 min, 8400 rpm). 200 µL of the resulting organic layer were utilised for GC-MS measurements. The remaining organic solvent was evaporated using a mild nitrogen stream, the precipitate was resolved in 200 µL isopropanol and utilised for chiral HPLC measurements.

For the larger scale hydroxylation reaction of *N*-phthaloyl-phenylethyl amine with *Cgl*UPO general procedures were followed as described above with some slight alterations. In contrast to the previous small scale reaction (500 µL), within this approach, ten reactions (each total volume: 1 mL) were performed in parallel in 100 mM potassium phosphate (pH 7.0) containing 250 nM *Cgl*UPO, 250 µM substrate and 250 µM $H_2O_2$. Reactions were performed for 60 minutes (30°C, 850 rpm) and subsequently quenched by the addition of 1 mL ethyl acetate to each reaction vial. Extraction was accomplished by 30 s of vigorous vortexing, followed by brief centrifugation (1 min, 8400 rpm). The organic layers of all samples were combined, and the solvent was gradually evaporated using a mild nitrogen stream. The precipitate was then resolved in 200 µL isopropanol and utilised for chiral HPLC measurements (Supplementary Figs. 16-18).

### II. Preparative work

##### ***N*-Phthaloyl-phenylethyl amine**

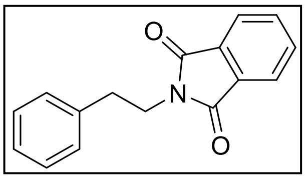

Phthalic anhydride (0.59 g, 4.0 mmol), phenylethyl amine (0.51 mL, 4.0 mmol) were dissolved in dichloromethane (40 mL) at room temperature. Molecular sieves (4Å pore diameter) and triethylamine (2.0 mL, 14.5 mmol) were added and the reaction mixture was refluxed for 36 h. After the reaction was completed (TLC control) the mixture was filtered, and the solvent was evaporated under reduced pressure. The residue was dissolved in ethyl acetate, washed with sodium bicarbonate solution and water and dried over sodium sulphate. After filtration, the product was obtained under reduced pressure to yield 0.31 g (80%) as an orange solid. No further purification was necessary.

**$^1$H-NMR** (400 MHz, $CDCl_3$): δ 7.83 (dd, *J* 5.4, 3.1 Hz, 2H), 7.70 (dd, *J* 5.5, 3.0 Hz, 2H), 7.32 – 7.17 (m, 5H), 3.96 – 3.90 (m, 2H), 3.02 – 2.95 (m, 2H) ppm;

**$^{13}$C-NMR** (100 MHz, $CDCl_3$): δ 168.15, 137.99, 133.88, 132.06, 128.83, 128.53, 126.62, 123.19, 39.27, 34.60 ppm;

**MS** (ESI, MeOH): m/z 274.1 ([M+Na]$^+$), calcd: 251.09.

##### **(R,S)-2-*N*-Phthaloyl-1-phenylethanol**

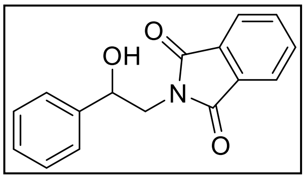

Phthalic anhydride (0.30 g, 2.0 mmol) and 2-amino-1-phenylethanol (0.27 g, 2.0 mmol) were placed into a microwave vessel under stirring (magnetic). The vessel was heated to 150 °C for 30 minutes in the microwave reactor. After cooling to room temperature, the product was washed with HCl (1 M, 20 mL) and recrystallised from dichloromethane/n-hexane to yield 0.47 g (89%) as colourless crystals.

**$^1$H-NMR** (400 MHz, $CDCl_3$): δ 7.82 (dd, *J* 5.4, 3.1 Hz, 2H), 7.70 (dd, *J* 5.5, 3.0 Hz, 2H), 7.48 – 7.40 (m, 2H), 7.39 – 7.27 (m, 3H), 5.06 (dt, *J* 8.6, 4.2 Hz, 1H), 4.07 – 3.85 (m, 2H), 3.03 (d, *J* 5.0 Hz, 1H) ppm;

**$^{13}$C-NMR** (100 MHz, $CDCl_3$): δ 168.69, 141.02, 134.06, 131.81, 128.53, 128.03, 125.83, 123.39, 72.47, 45.67 ppm;

**MS** (ESI, MeOH): m/z 268.1 ([M+H]$^+$), 290.0 ([M+Na]$^+$), calcd: 267.09.

##### **(S)-(+)-2-*N*-Phthaloyl-1-phenylethanol (chemical conversion)**

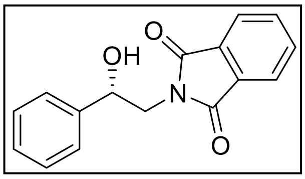

Phthalic anhydride (0.30 g, 2.0 mmol) and (S)-(+)-2-amino-1-phenylethanol (0.27 g, 2.0 mmol) were placed into a microwave vessel under stirring (magnetic). The vessel was heated to 150 °C for 30 minutes in the microwave reactor. After cooling to room temperature, the product was washed with HCl (1 M, 20 mL) and recrystallised from dichloromethane/n-hexane to yield 0.44 g (82%) as colourless crystals.

**$^1$H-NMR** (400 MHz, $CDCl_3$): δ 7.85 (dd, *J* 5.5, 3.1 Hz, 2H), 7.73 (dd, *J* 5.5, 3.1 Hz, 2H), 7.50 – 7.27 (m, 5H), 5.19 – 4.96 (m, 1H), 4.10 – 3.86 (m, 2H), 2.97 – 2.78 (m, 1H) ppm;

**$^{13}$C-NMR** (100 MHz, $CDCl_3$): δ 168.75, 141.05, 134.13, 131.88, 128.60, 128.11, 125.86, 123.46, 72.68, 45.76 ppm;
**MS** (ESI, MeOH): m/z 289.9 ($[M+Na]^+$), calcd: 267.09;
$[a]_{20}^{D}$ +23.9 (c 0.75, $CHCl_3$).

**(S)-(+)-2-*N*-Phthaloyl-1-phenylethanol (enzymatic conversion)**

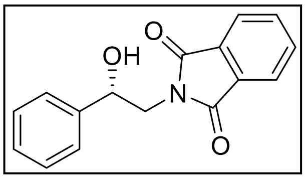

*N*-Phthaloyl-phenylethyl amine (15.8 mg, 62.9 µmol) was dissolved in acetone (15 mL) and poured into a solution of potassium phosphate buffer (100 mM, 263 mL, pH 7.0), hydrogen peroxide (210 µM, 3.2 mL) and *Mth*UPO (250 nM, 15 mL). The solution (total: 300 mL) was stirred at 30 °C for 1h. Afterwards the mixture was extracted using ethyl acetate (3x 60 ml). The organic phase was washed with brine, dried with sodium sulphate, filtered and concentrated under reduced pressure. The crude product was purified by column chromatography on silica gel using dichloromethane/ethyl acetate with 1 % formic acid (1/5 → 1/1) obtaining 9.70 mg (57 %) (S)-(+)-2-*N*-Phthaloyl-1-phenylethanol as a pale-yellow solid.
**$^1$H-NMR** (400 MHz, $CDCl_3$): δ 7.86 (dd, *J* 5.5, 3.1 Hz, 2H), 7.73 (dd, *J* 5.5, 3.0 Hz, 2H), 7.49 – 7.43 (m, 2H), 7.42 – 7.27 (m, 3H), 5.08 (dd, *J* 8.7, 3.6 Hz, 1H), 4.11 – 3.86 (m, 2H), 2.83 (s, 1H) ppm;
**$^{13}$C-NMR** (100 MHz, $CDCl_3$): δ 168.76, 141.04, 134.14, 131.89, 128.62, 128.13, 125.86, 123.48, 72.72, 45.77 ppm;
**MS** (ESI, MeOH): m/z 289.9 ($[M+Na]^+$), calcd: 267.09;
$[a]_{20}^{D}$ +21.0 (c 1.55, $CHCl_3$).

***N*-Phthaloyl-2-oxo-phenylethyl amine**

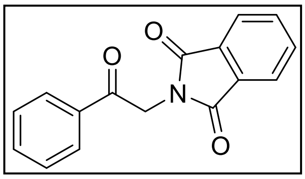

(R,S)-*N*-Phthaloyl-phenylethanol (0.18 g, 0.67 mmol) was dissolved in dimethyl sulfoxide (6 mL) at room temperature. Under ice cooling, acetic anhydride (1.2 mL) was added and the reaction mixture was stirred for 16 h at room temperature. After the reaction was completed (TLC control) the mixture was quenched with ethyl acetate (20 mL), and the mixture was washed with sodium perchlorate solution (6 %), sodium thiosulfate solution (10 %) and brine and dried over sodium sulfate. After filtration, the product was obtained under reduced pressure to yield 0.15 g (84 %) as a colourless solid. No further purification was necessary.
**$^1$H-NMR** (400 MHz, $CDCl_3$): δ 8.06 – 7.98 (m, 2H), 7.91 (dd, *J* = 5.5, 3.0 Hz, 2H), 7.76 (dd, *J* = 5.5, 3.0 Hz, 2H), 7.69 – 7.48 (m, 3H), 5.14 (s, 2H) ppm;
**$^{13}$C-NMR** (100 MHz, $CDCl_3$): δ 190.94, 167.88, 134.43, 134.11, 134.02, 132.25, 128.89, 128.14, 123.55, 44.19 ppm;
**MS** (ESI, MeOH): m/z 288.1 ($[M+Na]^+$), calcd: 265.07.

### III. Supplementary Figures

**Supplementary Fig. 1 Principle of modular UPO shuffling system**

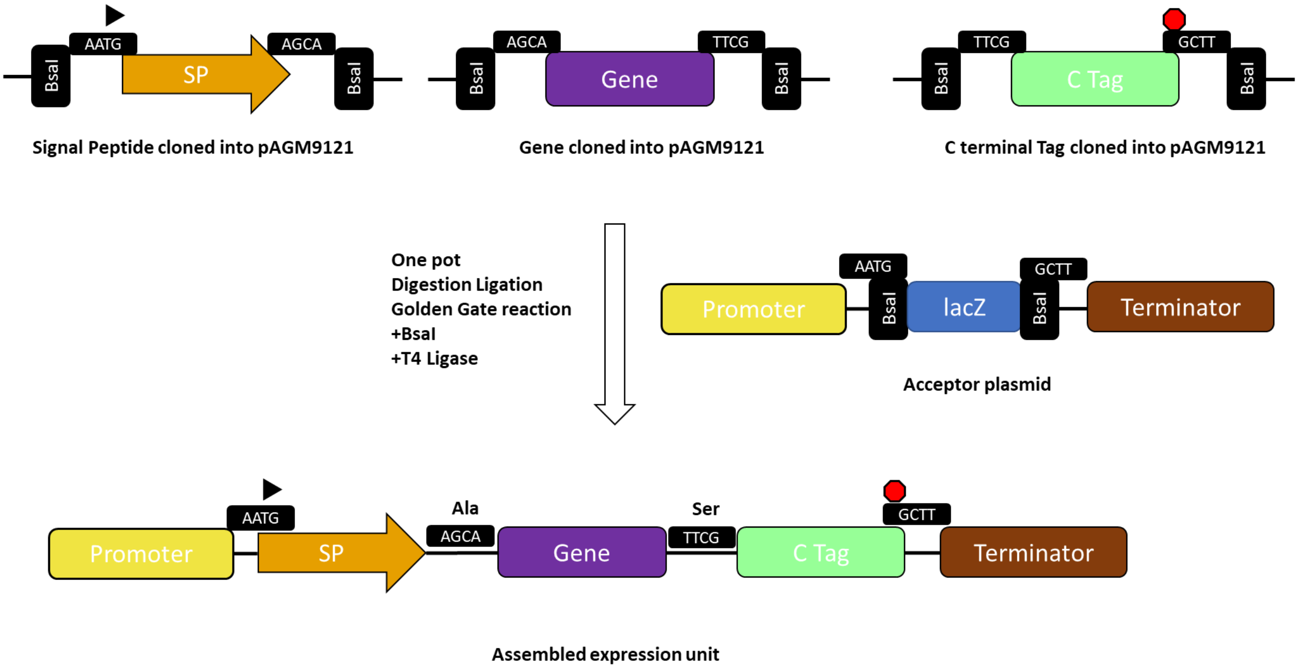

Primer design example for adding a gene unit into the modular *Yeast Secret and Detect* system (clone as Level 0 module into pAGM9121):

Forward Primer:

TTGAAGACAACTCAAGCAXXXXXXXXXXXXX

Reverse Primer:

TTGAAGACAACTCGCGAAXXXXXXXXXXXXX

TT- Prefix
GAAGAC- BbsI recognition
AA- Suffix (BbsI cleavage pattern)
CTCA- 4 bp overhang complementary to pAGM9121
AGCA- 4 bp overhang defining module 2 position (gene part) in *Yeast Secret and Detect* system
XXXX-Binding sequence of the protein-coding template (target protein)

**Supplementary Fig. 2 Comparison of C-terminal GFP11 and TwinStrep-GFP11 constructs (*Gma*-UPO-*Aae*UPO* constructs; 27 biological replicates)**

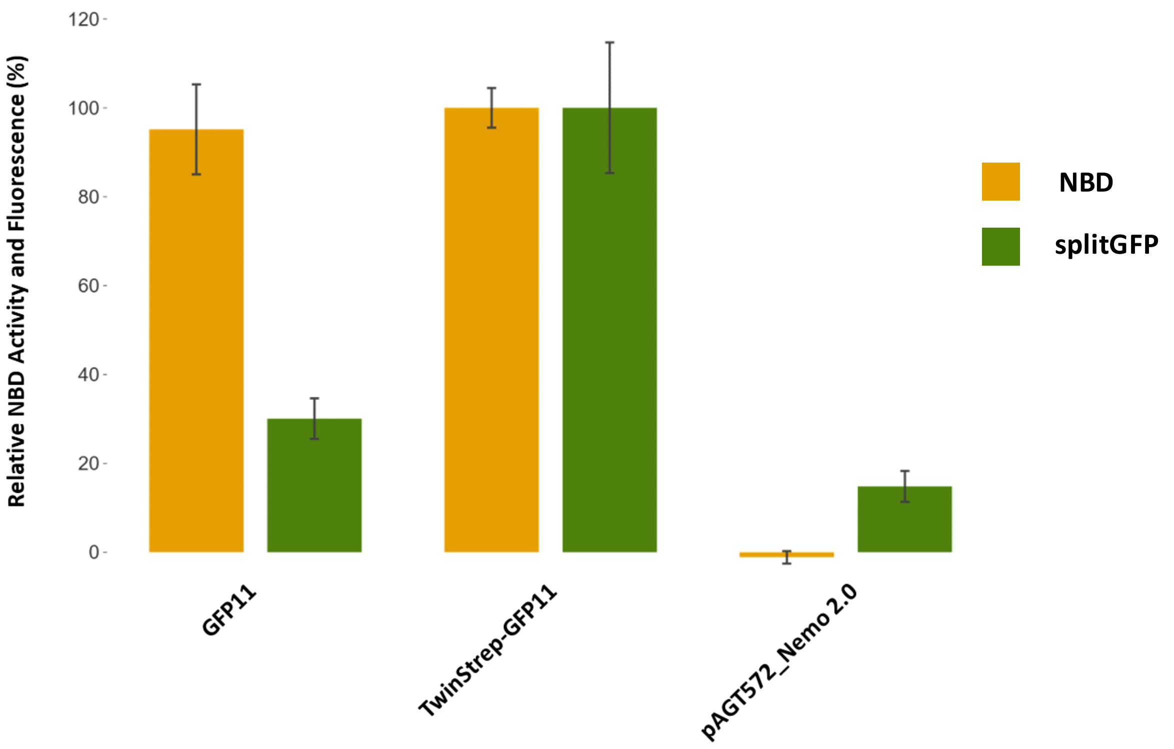

**Supplementary Fig. 3 Identification of suitable signal peptides for *Mth*UPO secretion in *S. cerevisiae* (5 biological replicates)**

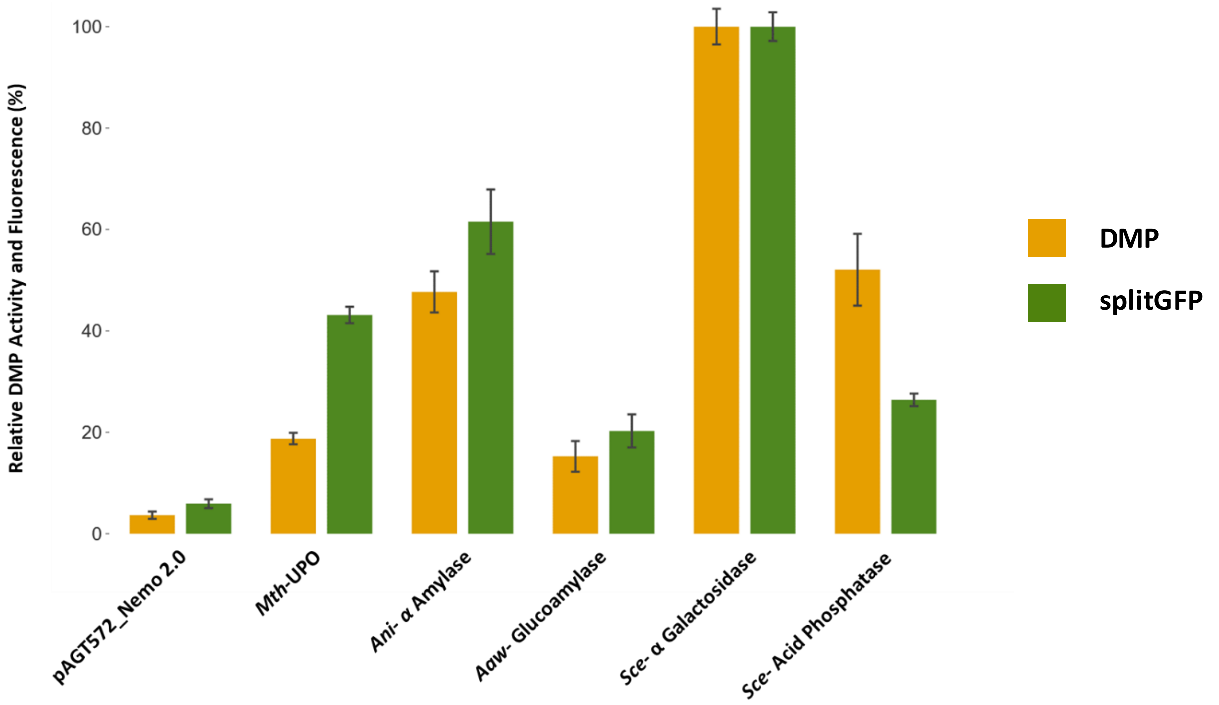

**Supplementary Fig. 4 Identification of suitable signal peptides for *Tte*UPO secretion in *S. cerevisiae* (8 biological replicates)**

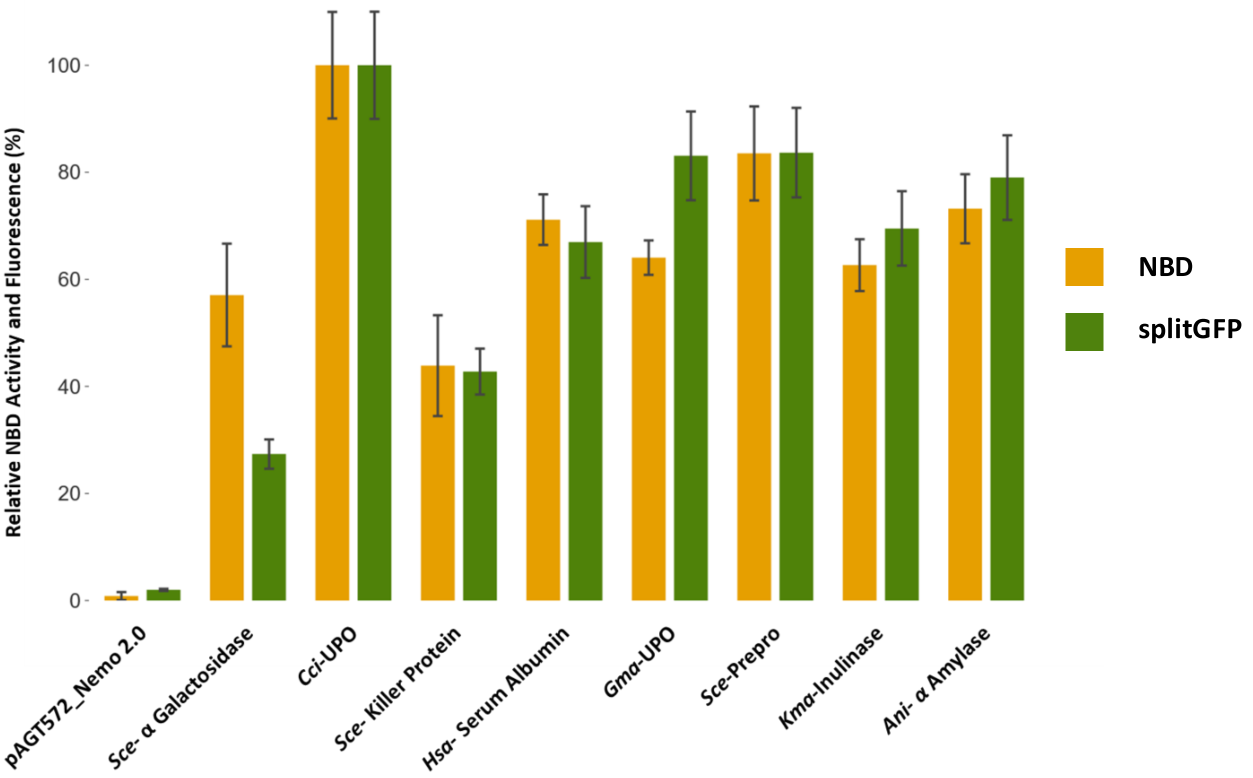

**Supplementary Fig. 5 CO absorption measurements of UPOs produced in *S. cerevisiae***

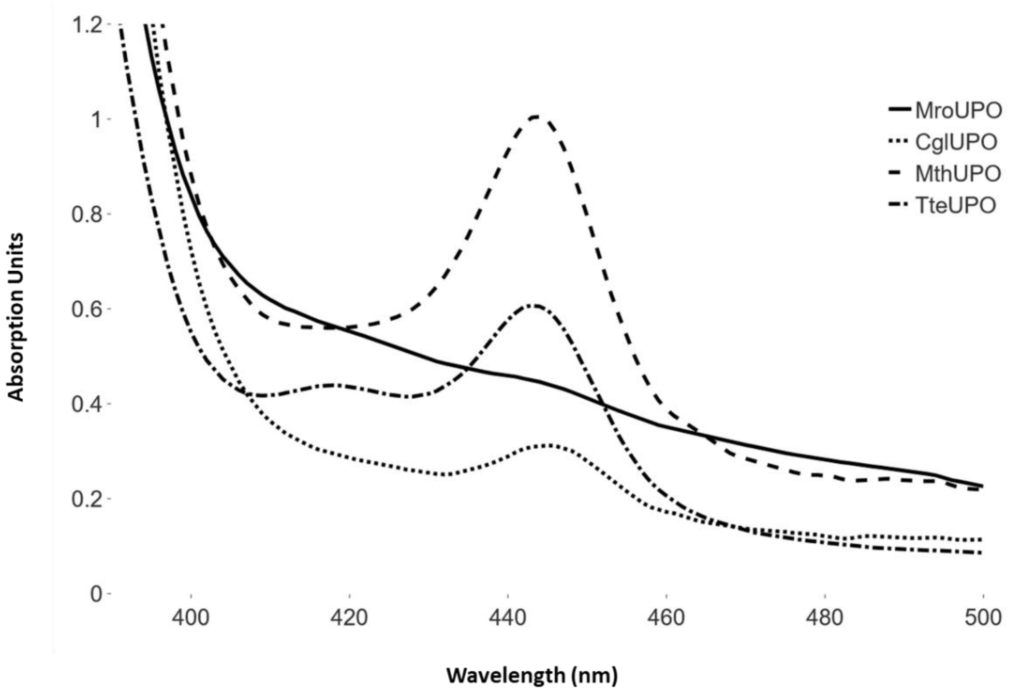

**Supplementary Fig. 6 SDS PAGE analysis of UPOs produced in *S. cerevisiae***

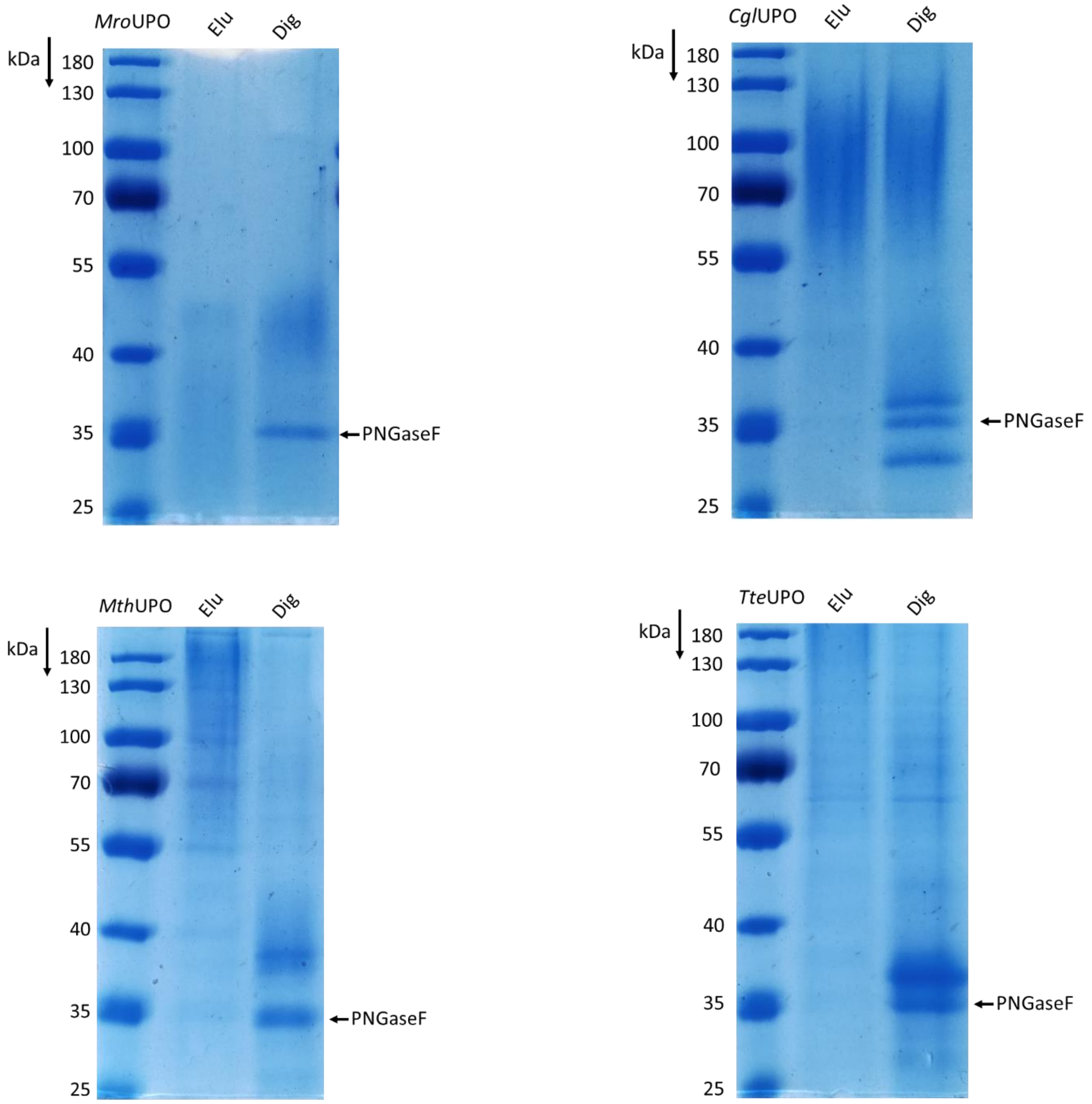

**Supplementary Fig. 7 Enzymatic activity of UPOs derived from *S. cerevisiae* after deglycosylation (24 h PNGase F treatment (37 °C); 3 technical replicates)**

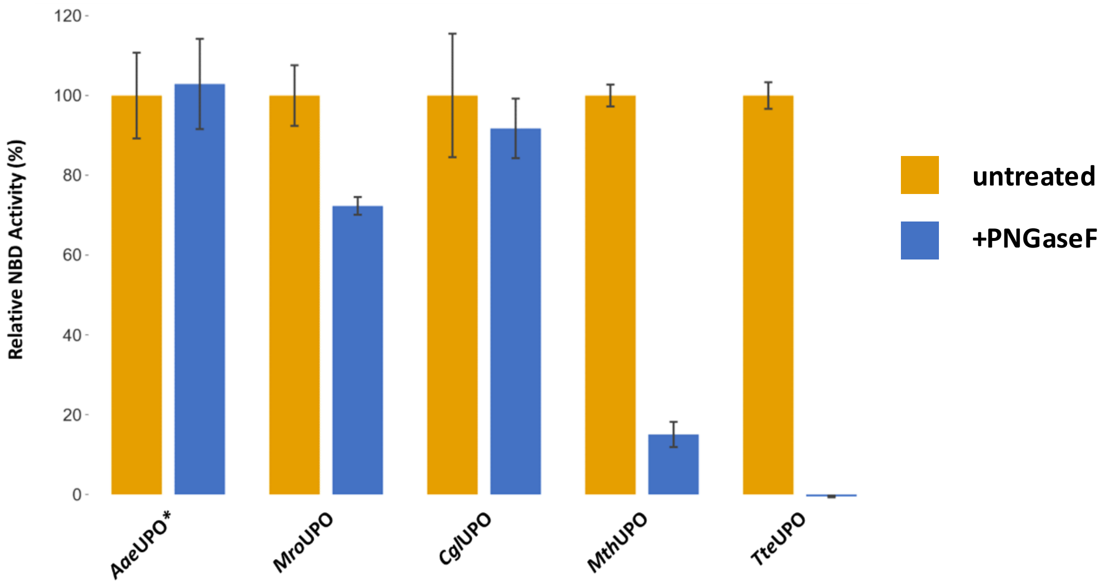

**Supplementary Fig. 8 Comparison of UPO production from episomal and integrative constructs *P. pastoris* (12 biological replicates)**

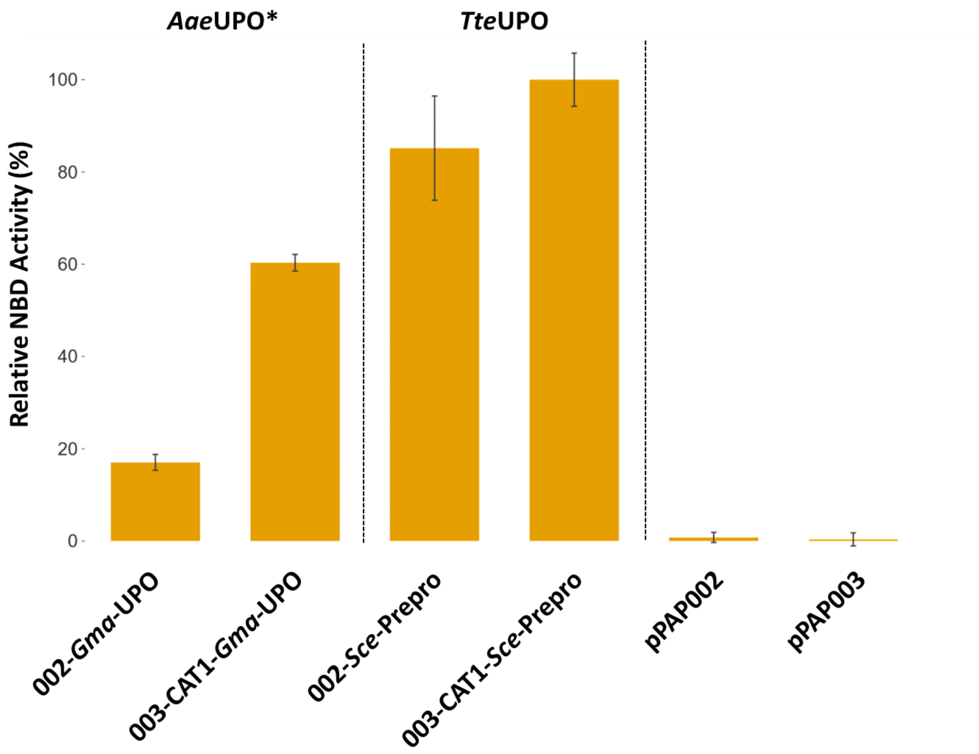

**Supplementary Fig. 9 Comparison colony shape and size episomal and integrative construct *P. pastoris***

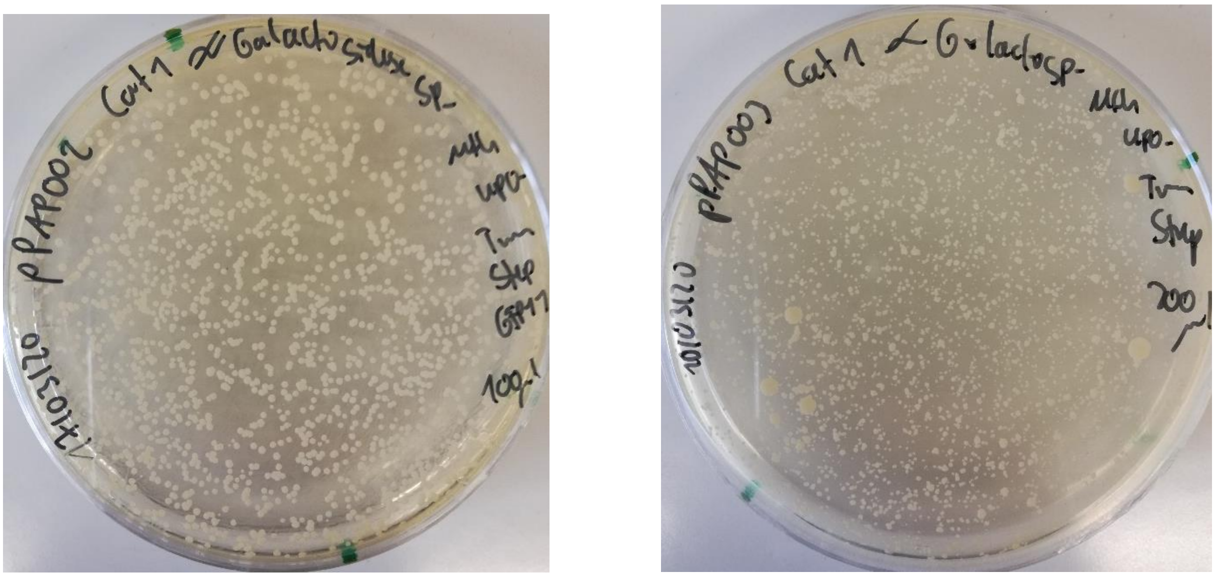

**episomal (pPAP002)** **integrative (pPAP003)**

**Supplementary Fig. 10 Identification of suitable signal peptides for *Mth*UPO secretion in *P. pastoris* (6 biological replicates)**

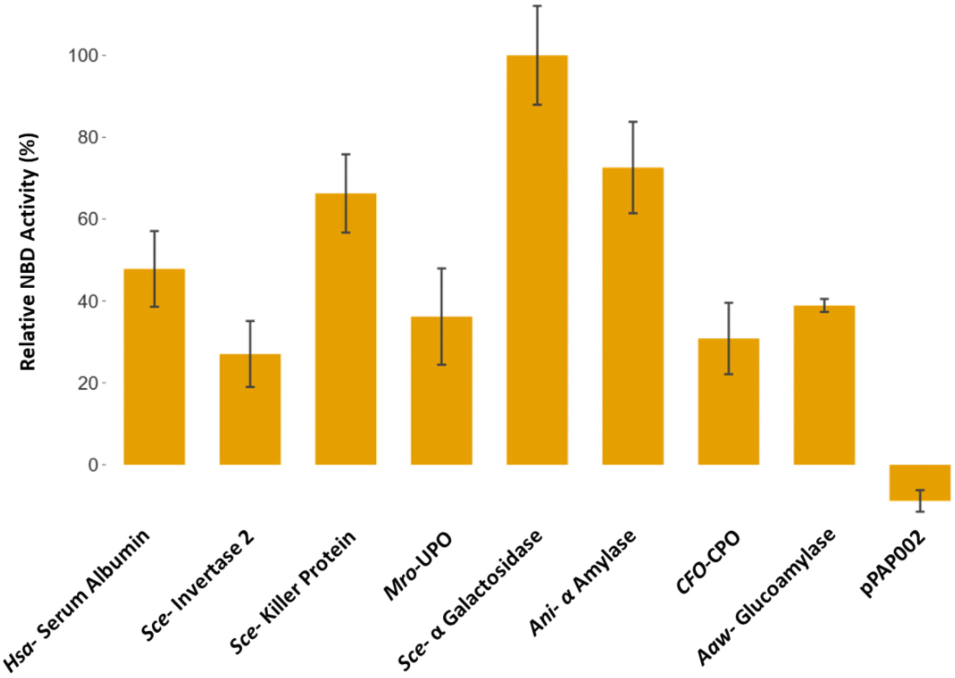

**Supplementary Fig. 11 Identification of suitable signal peptides for *Tte*UPO secretion in *P. pastoris* (6 biological replicates)**

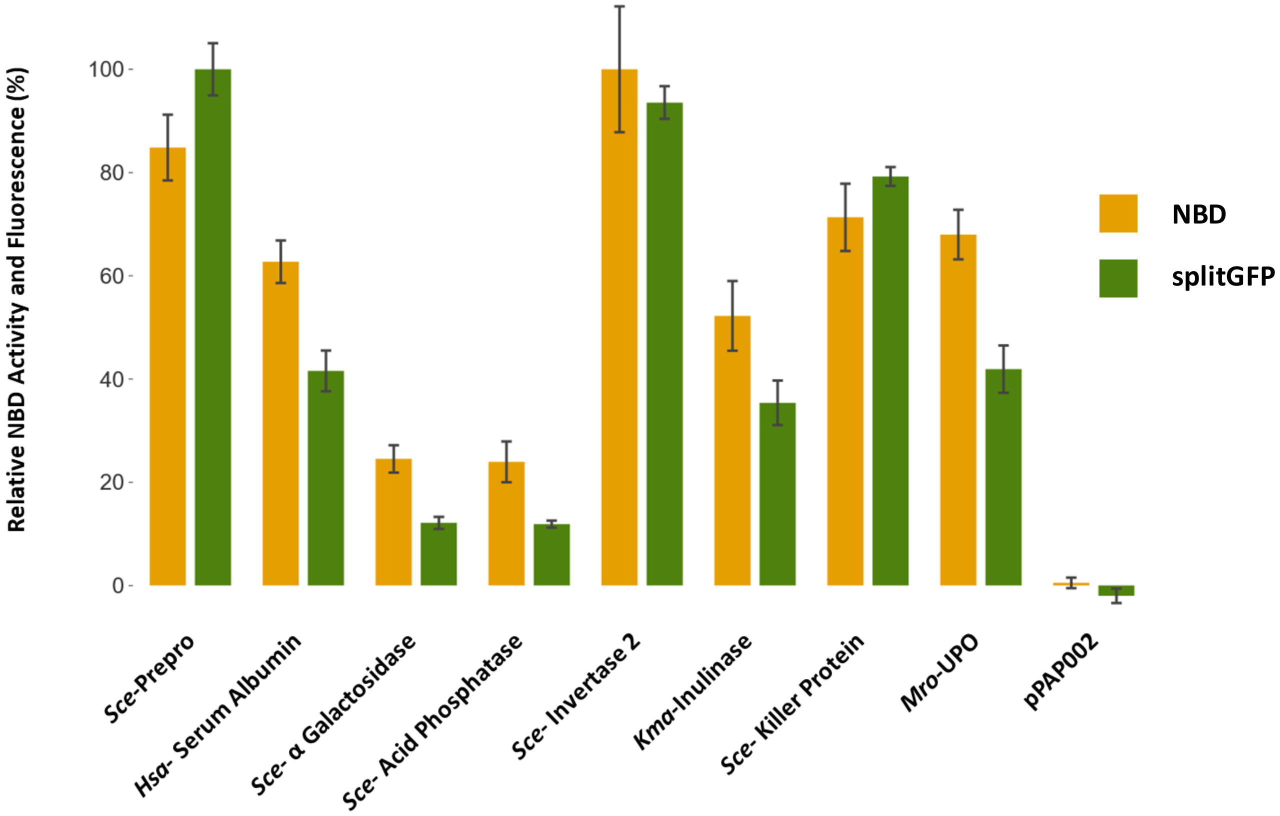

**Supplementary Fig. 12 SDS PAGE analysis of UPOs derived from *P. pastoris***

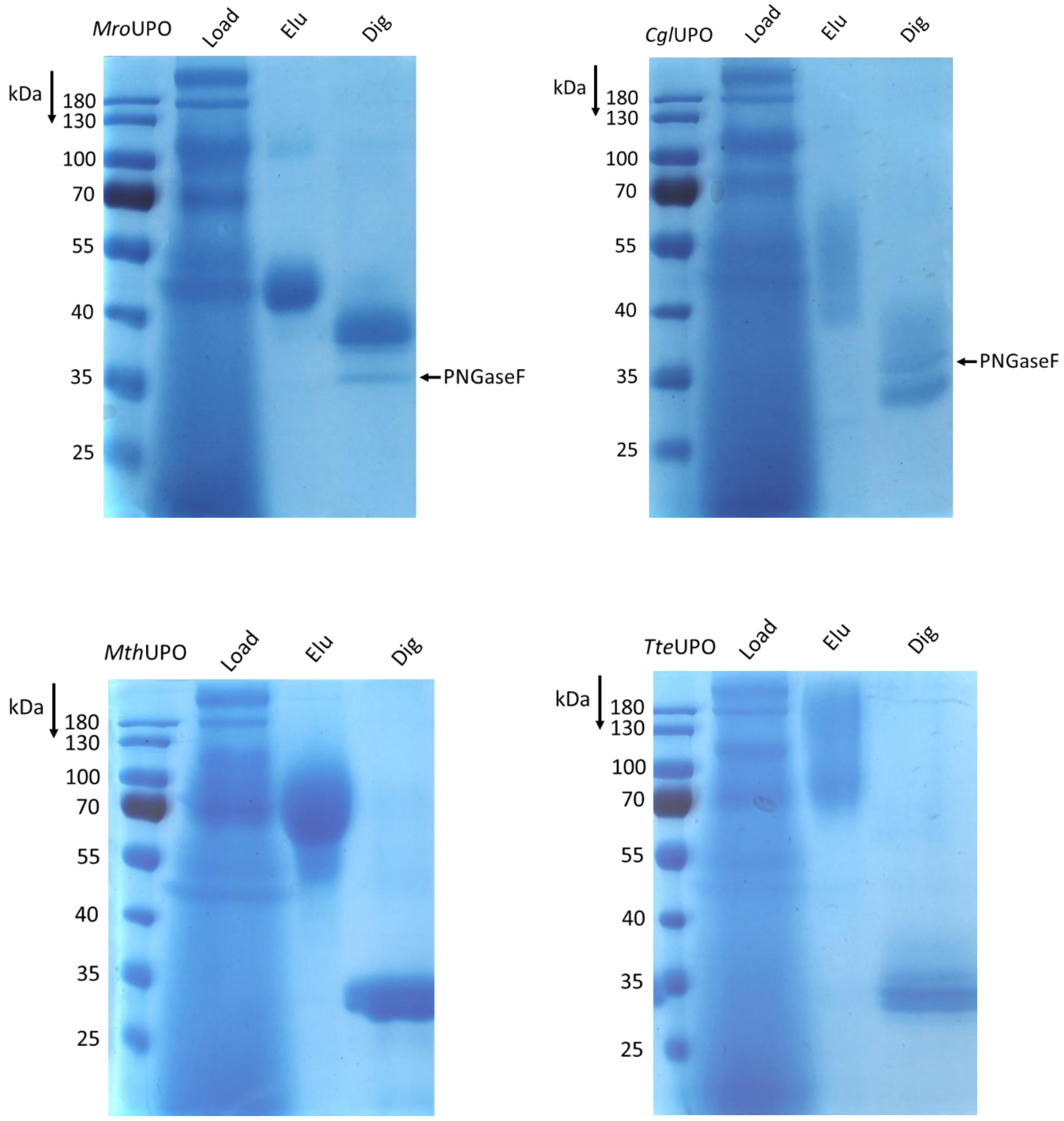

**Supplementary Fig. 13 Calibration curves GC-MS**

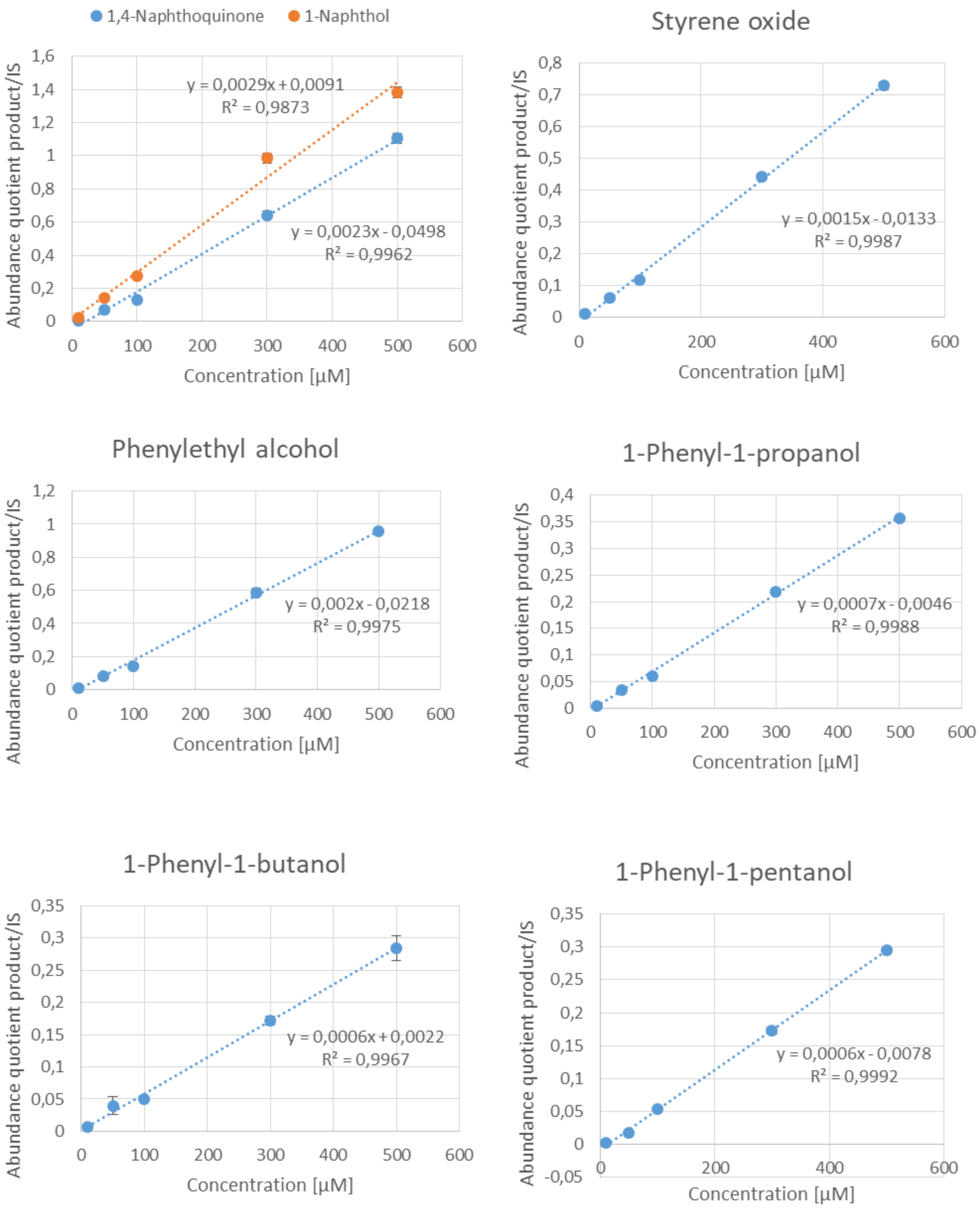

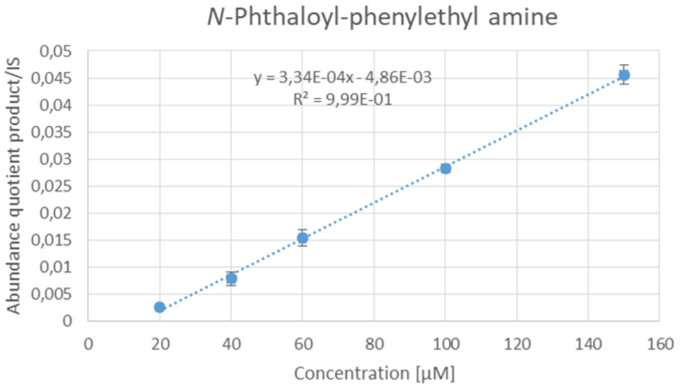

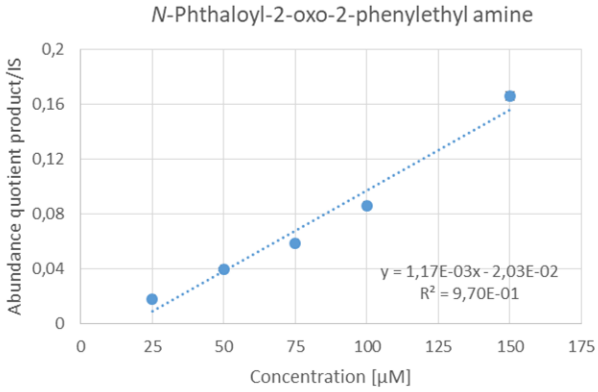

**Supplementary Fig. 14 Enantiomeric separation 1-phenylethyl alcohol products**

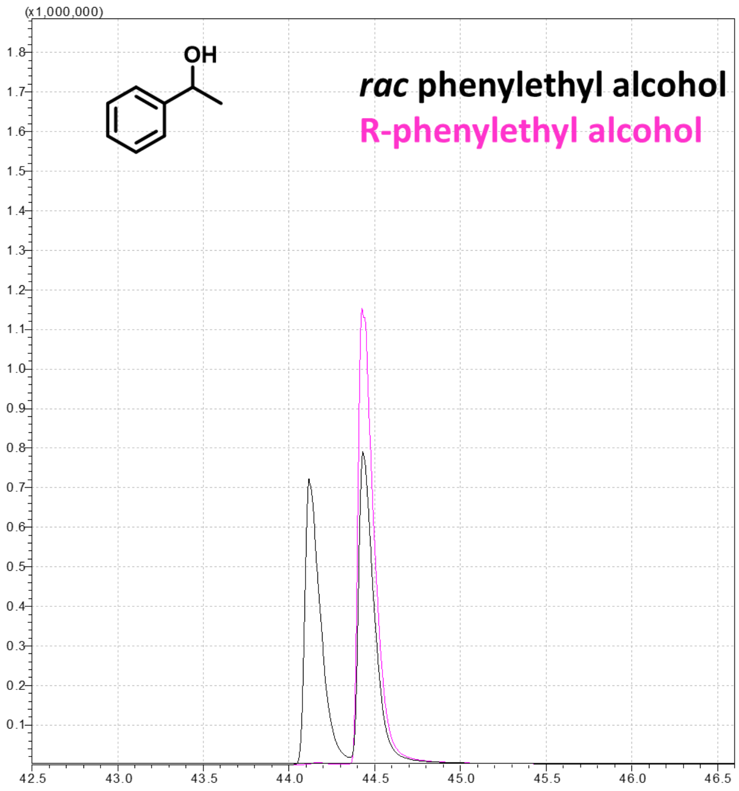

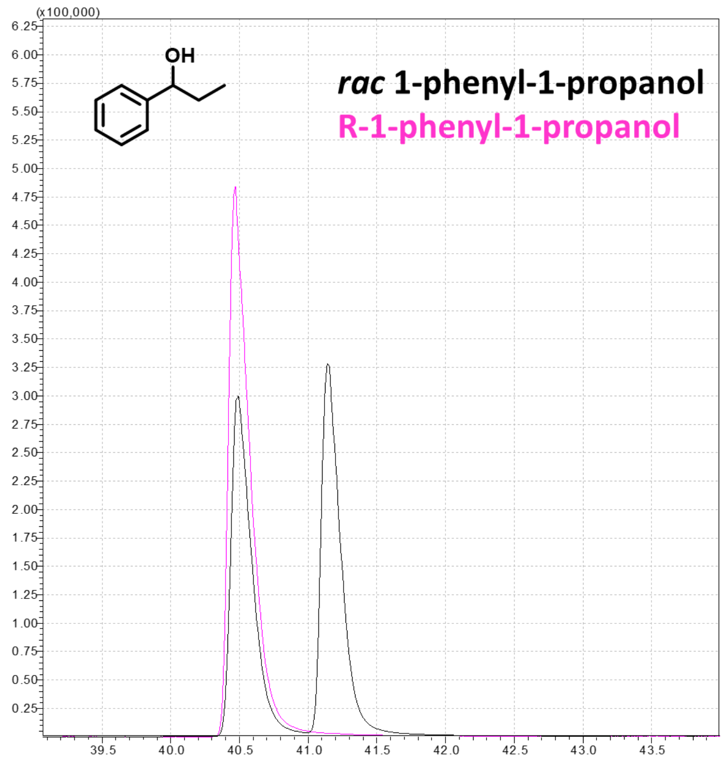
OH
rac 1-phenyl-1-propanol
R-1-phenyl-1-propanol
(x100,000)
6.25
6.00
5.75
5.50
5.25
5.00
4.75
4.50
4.25
4.00
3.75
3.50
3.25
3.00
2.75
2.50
2.25
2.00
1.75
1.50
1.25
1.00
0.75
0.50
0.25
39.5
40.0
40.5
41.0
41.5
42.0
42.5
43.0
43.5

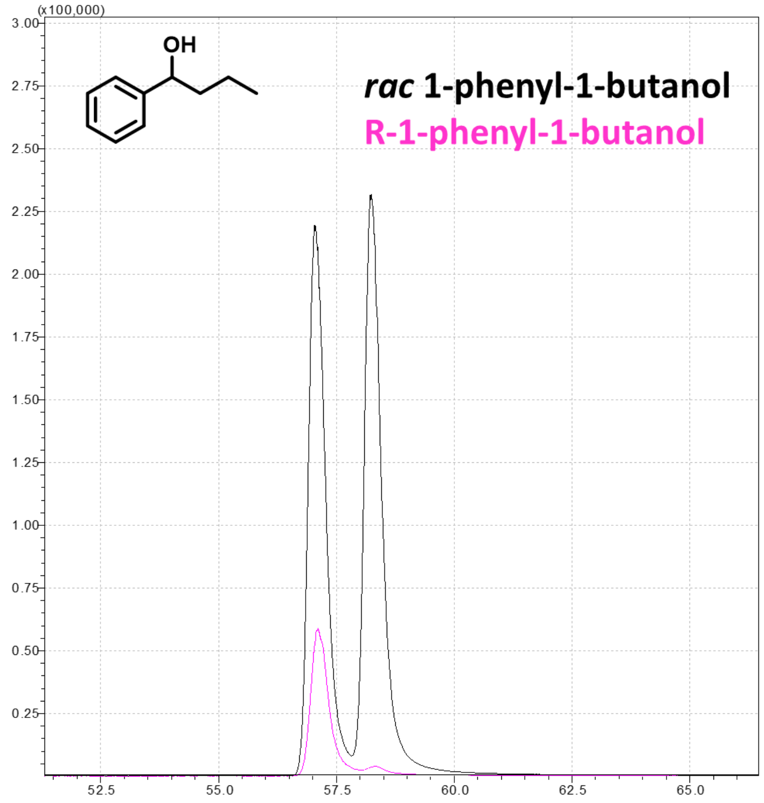
OH
rac 1-phenyl-1-butanol
R-1-phenyl-1-butanol
(x100,000)
3.00
2.75
2.50
2.25
2.00
1.75
1.50
1.25
1.00
0.75
0.50
0.25
52.5
55.0
57.5
60.0
62.5
65.0

**Supplementary Fig. 15 Scan Mode measurements of UPO catalysed Phenyl alkane hydroxylation**

**Phenylethane**

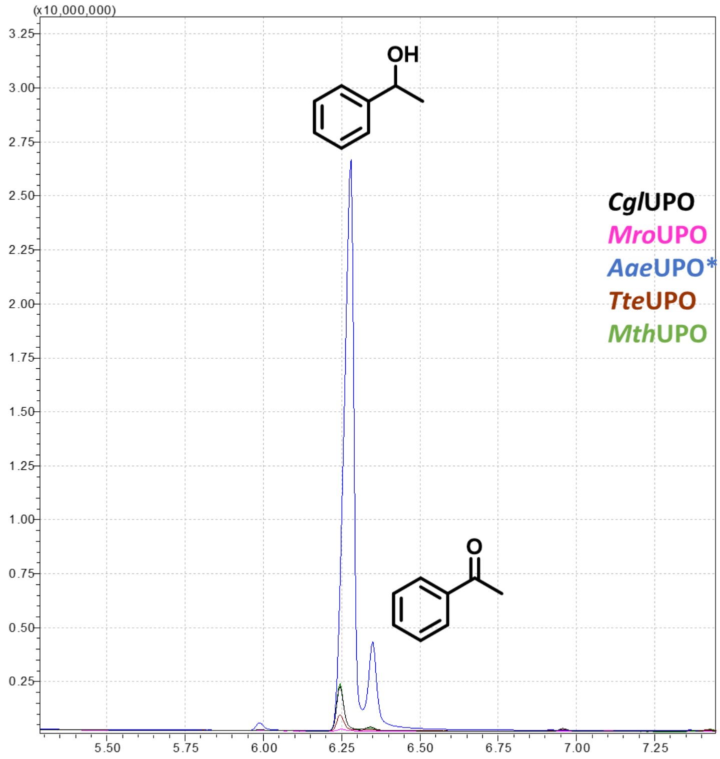

**Phenylpropane**

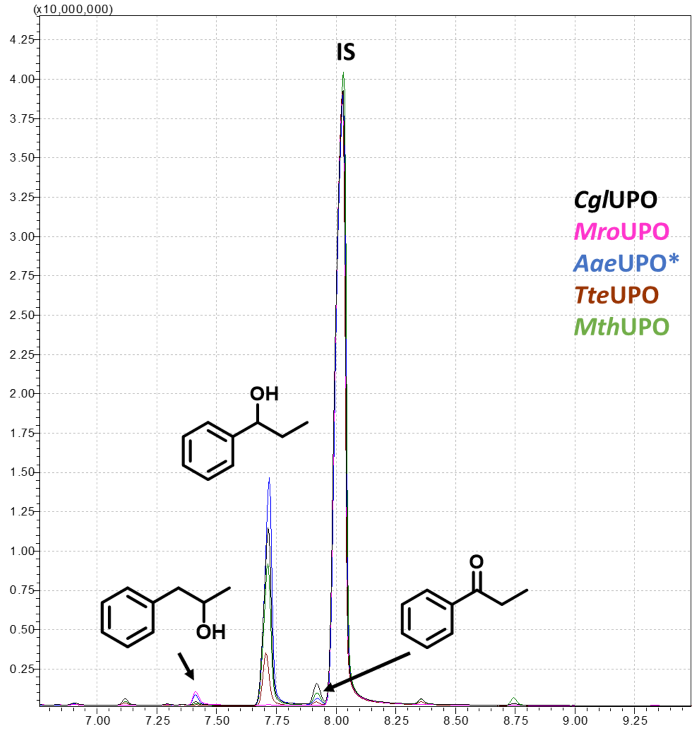

**Phenylbutane**

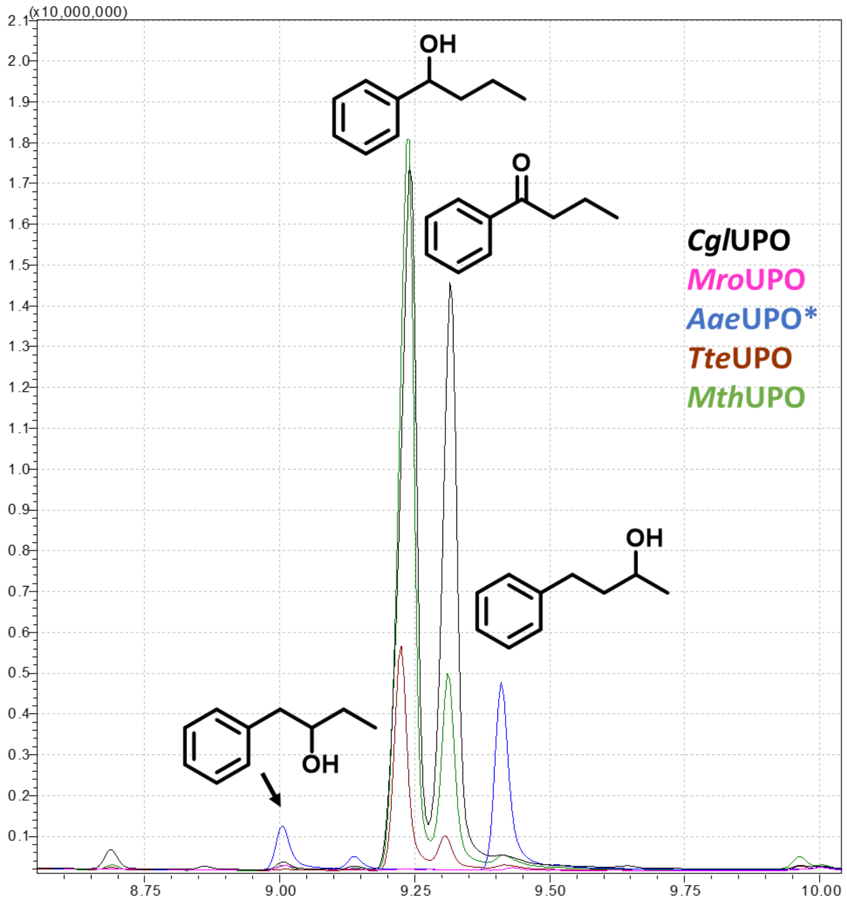

**Phenylpentane**

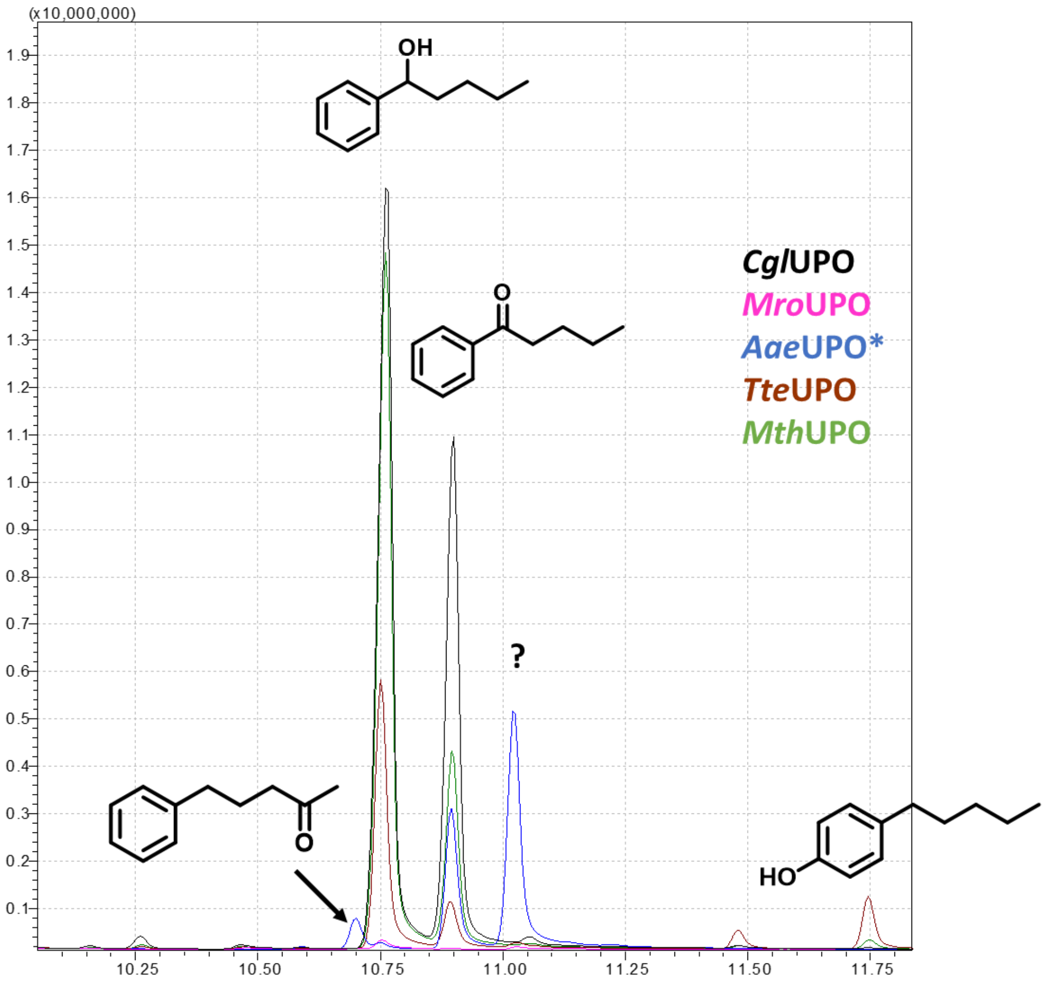

**Supplementary Fig. 16 SIM Mode measurements of UPO catalysed *N*-Phthaloyl-phenylethyl amine conversion**

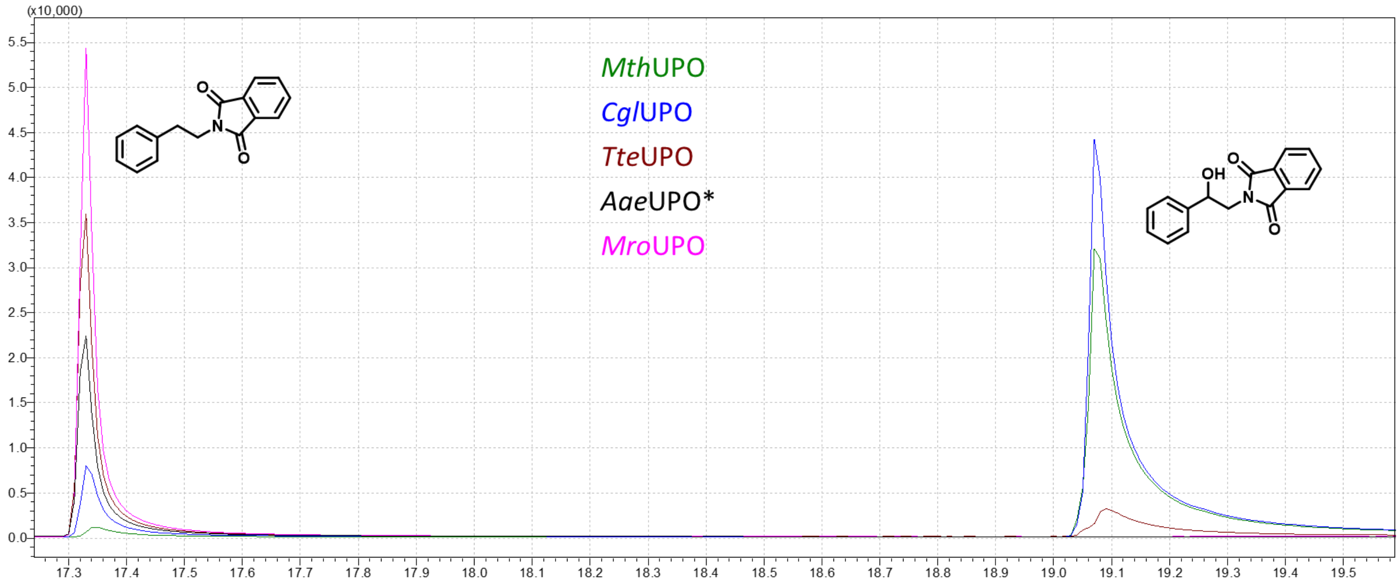

**Supplementary Fig. 17 Scan Mode measurements of UPO catalysed *N*-Phthaloyl-phenylethyl amine conversion**

***Mth*UPO**

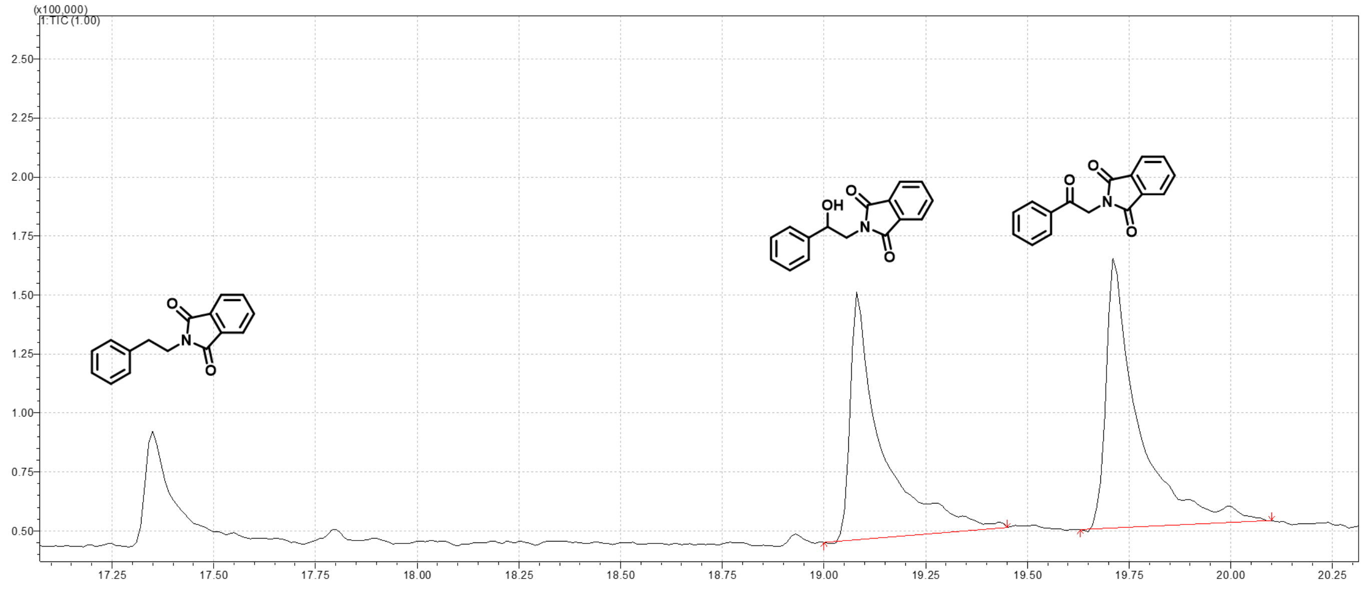

***Cgl*UPO**

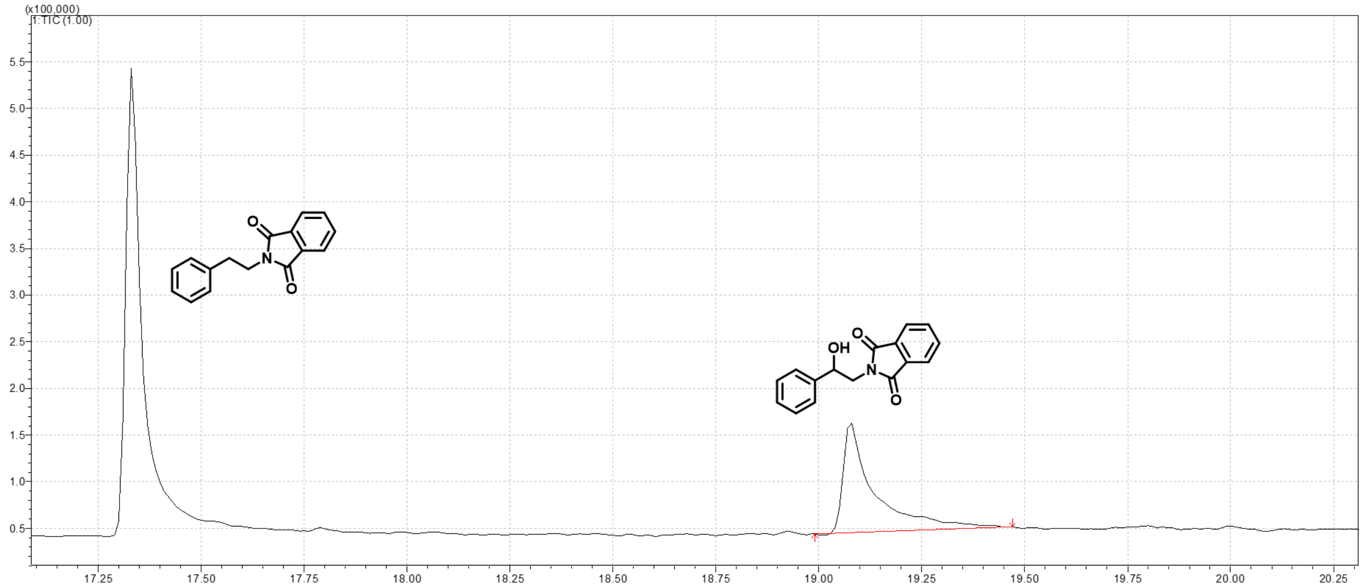

***Tte*UPO**

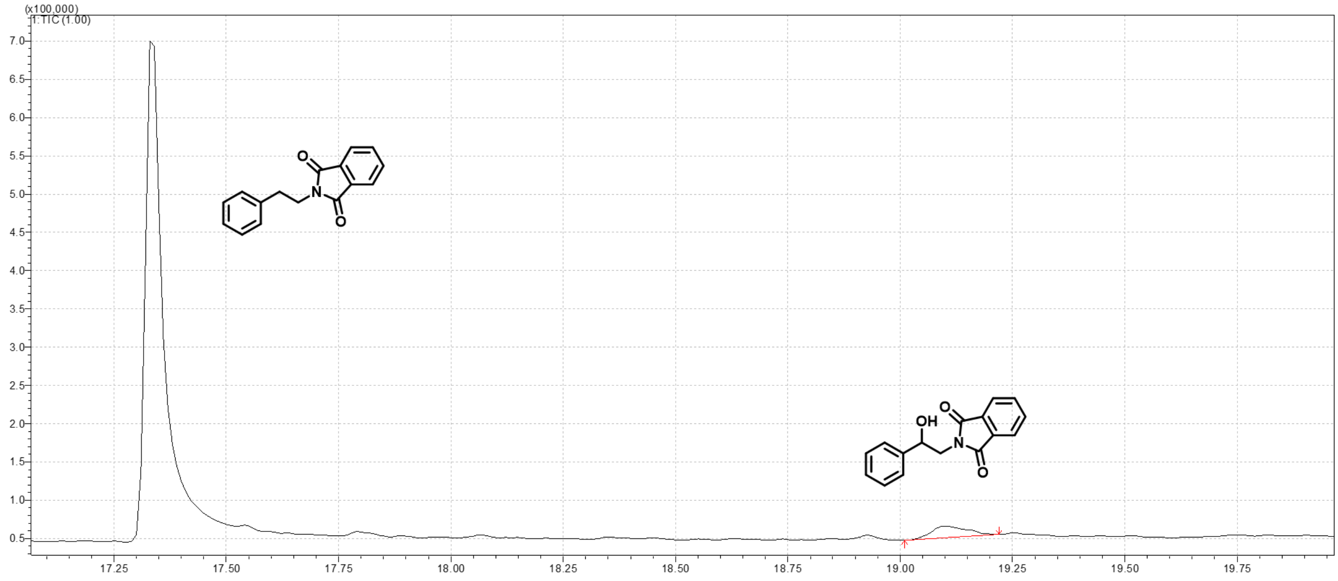

**Supplementary Fig. 18 Chiral HPLC analysis of UPO catalysed *N*-Phthaloyl-phenylethyl amine conversion**

**Small scale reactions (400 µL) and utilised standards**

*(R,S)-2-N-Phthaloyl-1-phenylethanol*

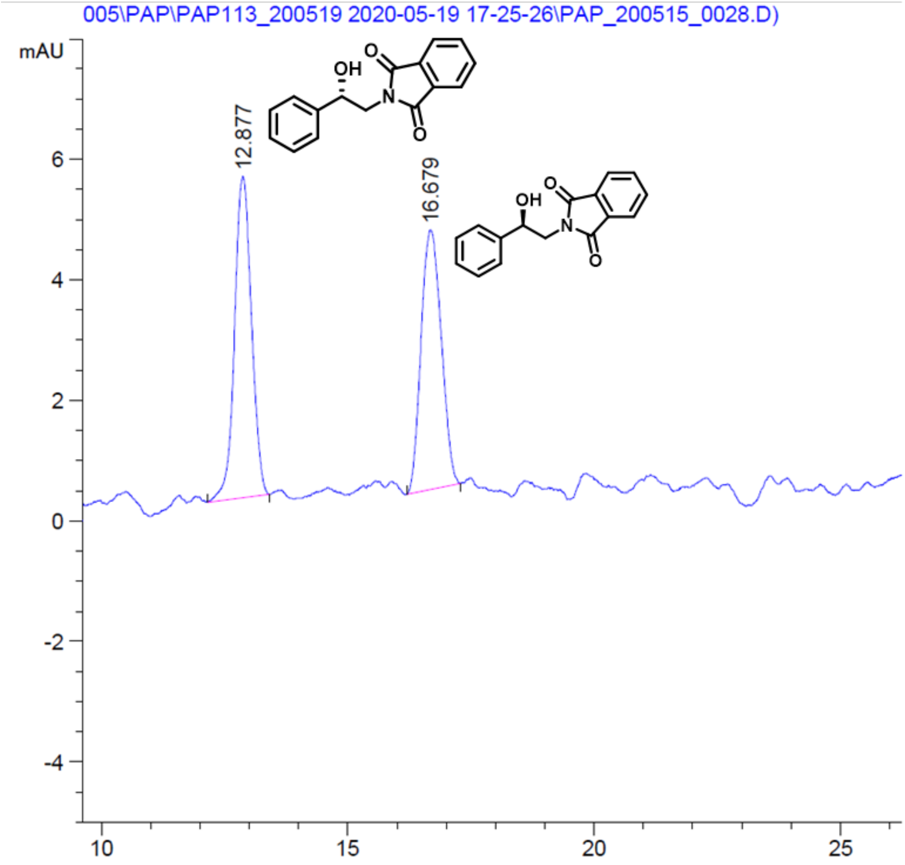

*(S)-(+)-2-N-Phthaloyl-1-phenylethanol*

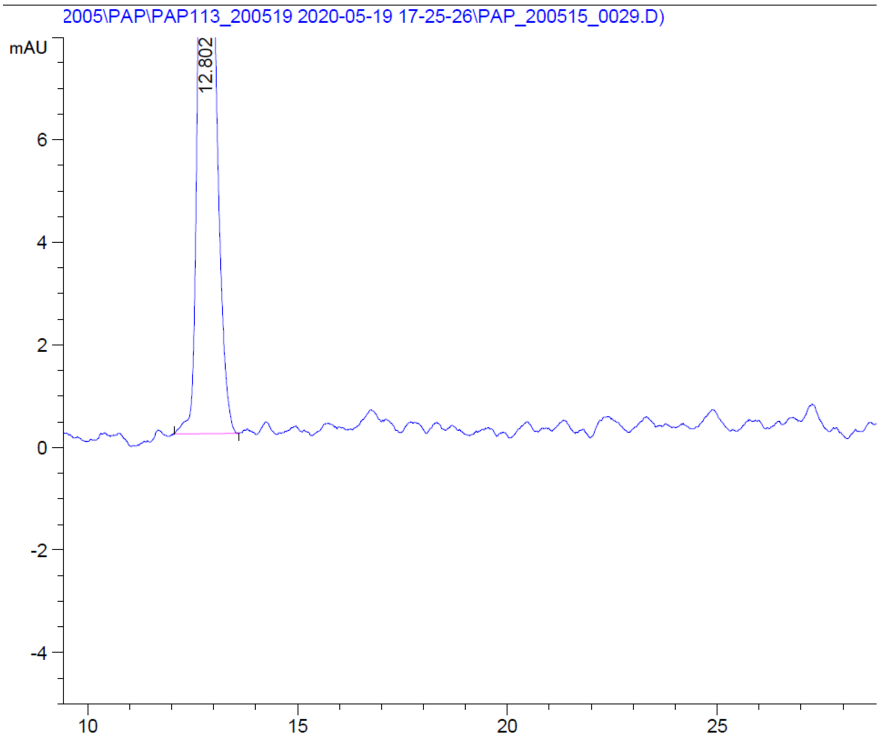

***Mth*UPO**

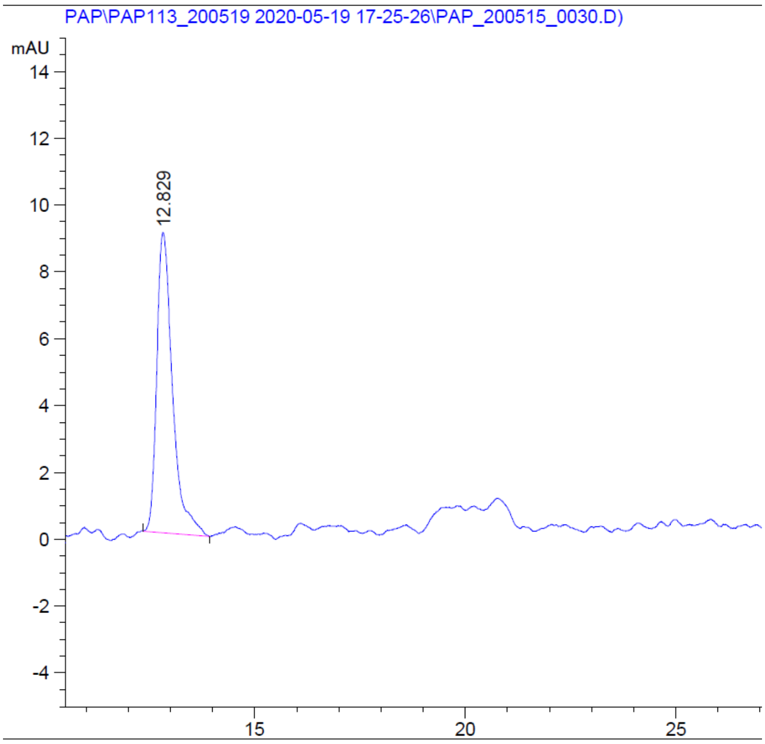

***Cgl*UPO**

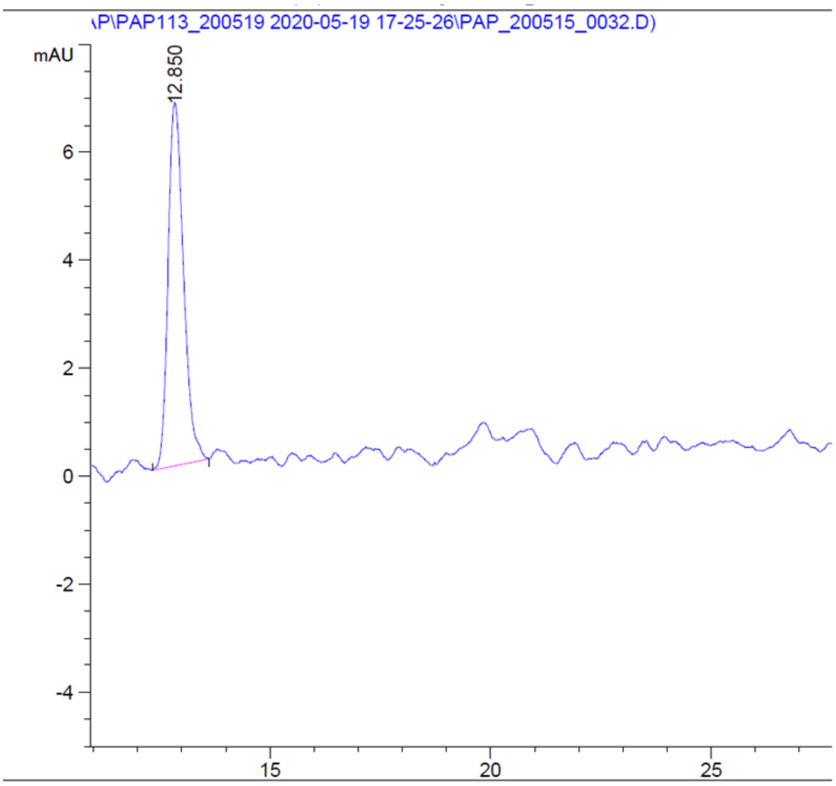

***Tte*UPO**

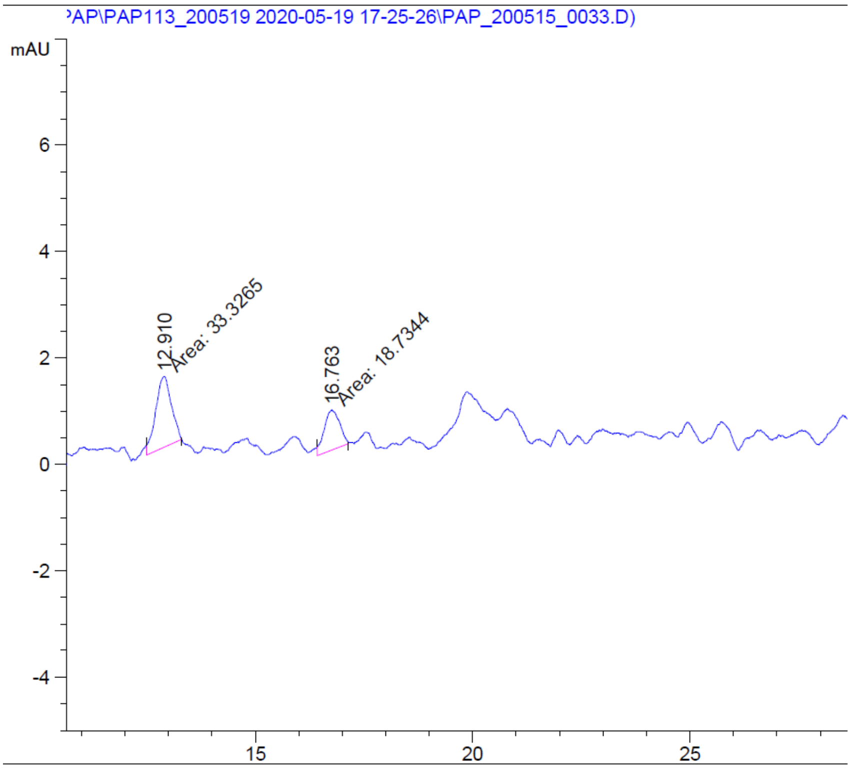

**Larger scale reactions and standards for enantiomeric excess determination**

***Mth*UPO**

*(R,S)-2-N-Phthaloyl-1-phenylethanol*

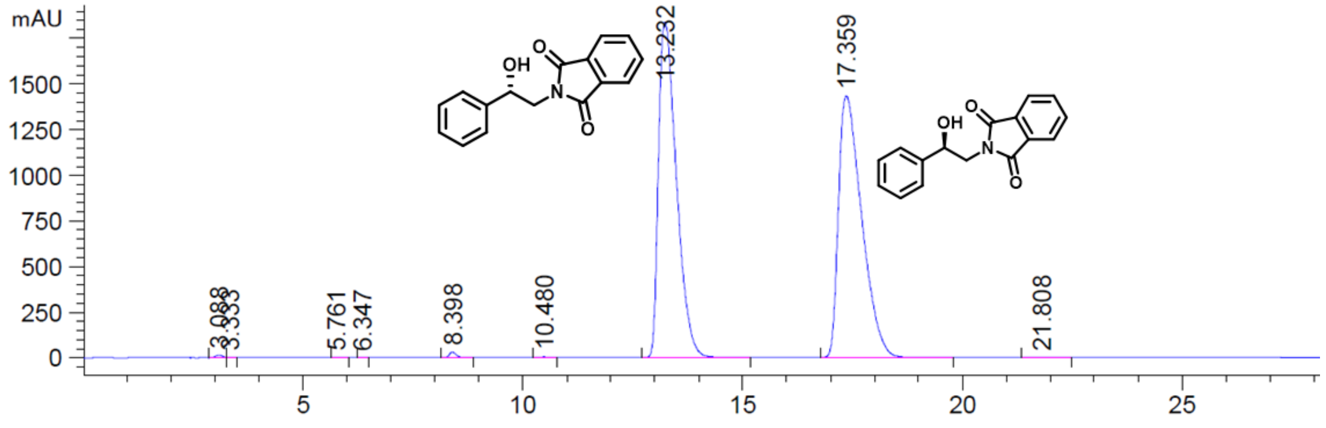

*(S)-(+)-2-N-Phthaloyl-1-phenylethanol*

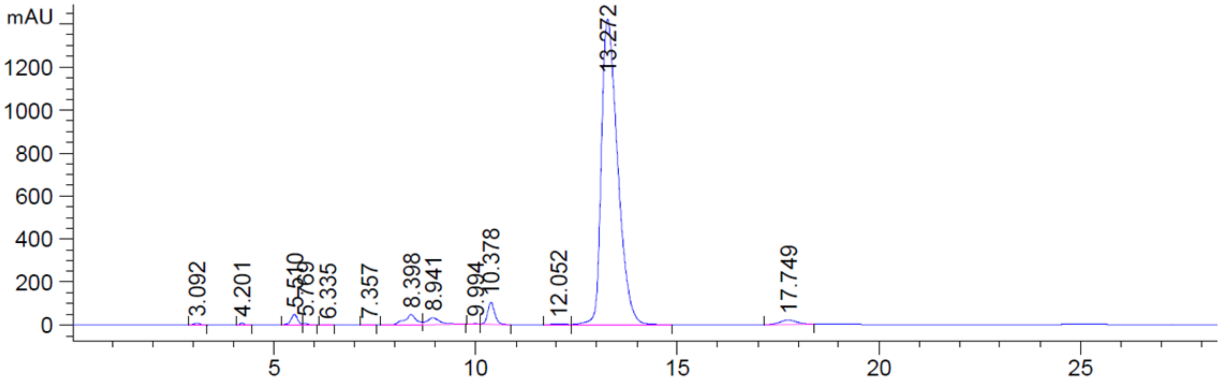

**Preparative enzymatic conversion (300 mL approach)**

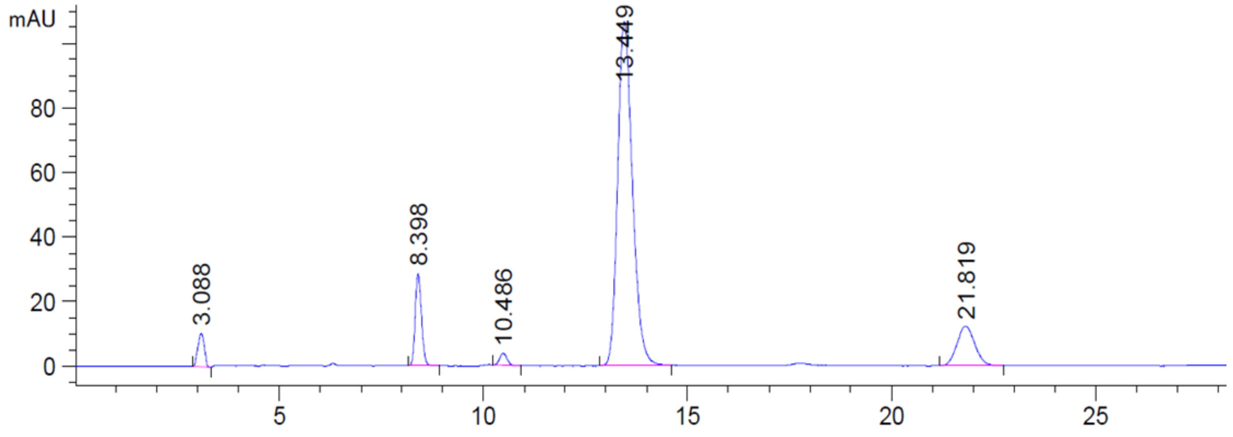

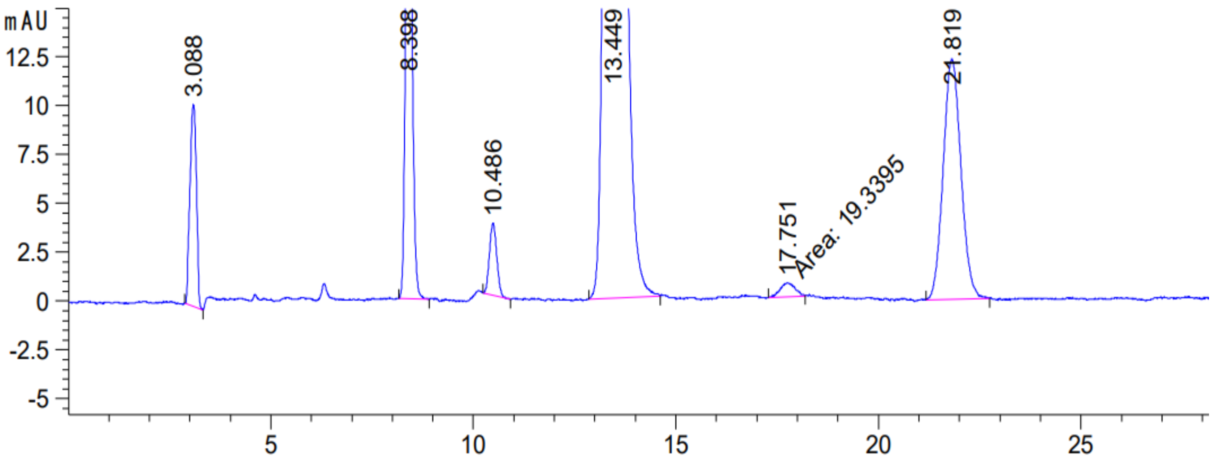
mAU
12.5
10
7.5
5
2.5
0
-2.5
-5
5
10
15
20
25
3.088
8.398
10.486
13.449
17.751
Area: 19.3395
21.819

#### *Cgl*UPO

*(R,S)-2-N-Phthaloyl-1-phenylethanol*

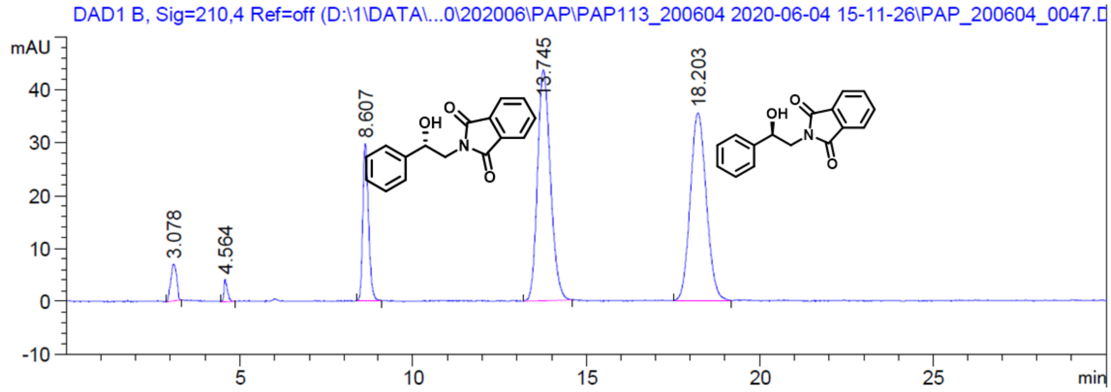

*(S)-(+)-2-N-Phthaloyl-1-phenylethanol*

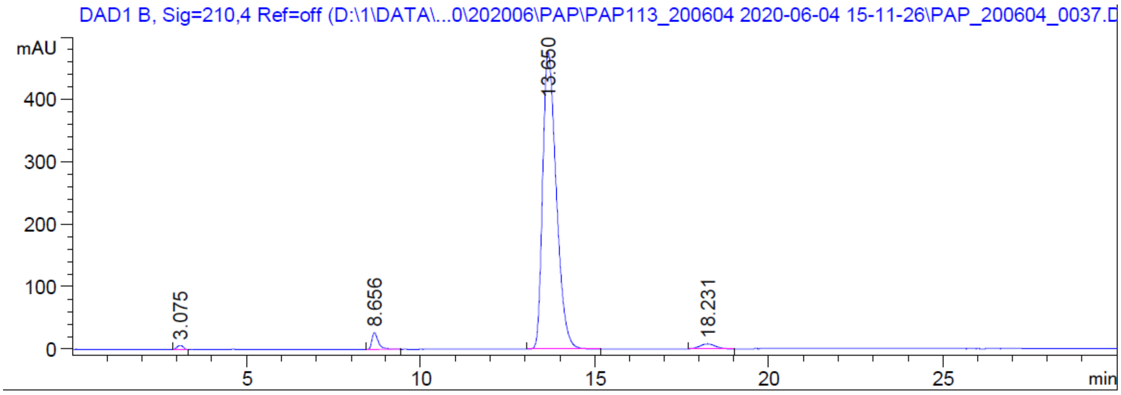

**Large scale enzymatic conversion (10 ml reaction setup)**

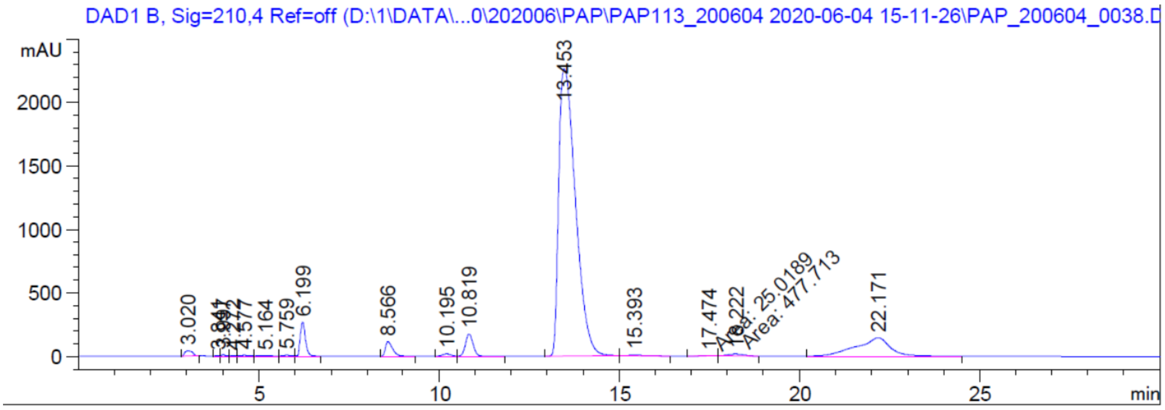

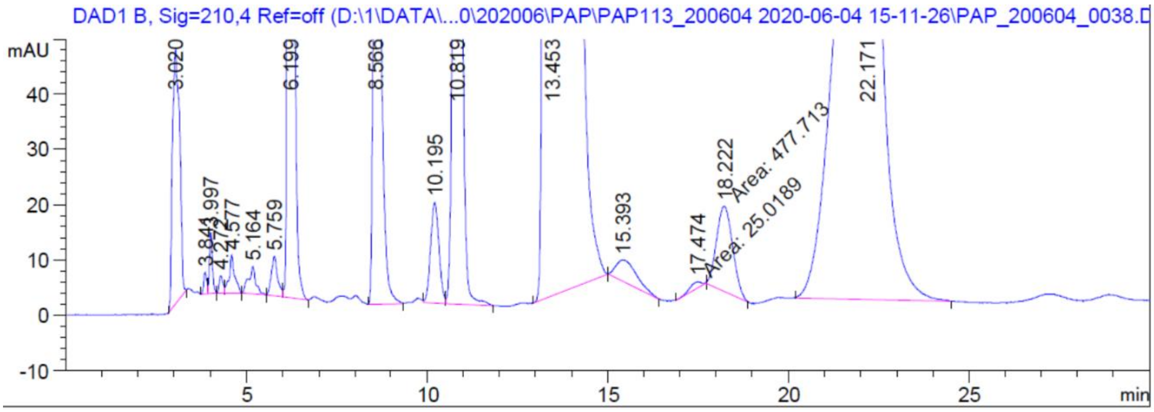

DAD1 B, Sig=210,4 Ref=off (D:\1\DATA\...0\202006\PAP\PAP113_200604 2020-06-04 15-11-26\PAP_200604_0038.D
mAU
3.020
3.841
3.997
4.272
4.577
5.164
5.759
6.199
8.566
10.195
10.819
13.453
15.393
17.474
18.222
Area: 477.713
Area: 25.0189
22.171
min

### IV. Supplementary Tables

**Supplementary Table 1 Overview of oligonucleotides for sequencing of the created plasmids**

| ***Name*** | ***Sequence (5´ → 3´)*** |
|---|---|
| *pAGM9121_for* | CCTGTCGGGTTTCGCCACCT |
| *pAGM9121_rev* | GCCGTTACCACCGCTGCGTT |
| *pAGT572_Nemo 2.0_for* | CATCTTATTAAAGTATCATCAAGAAATTGTTA |
| *pAGT572_Nemo_rev* | AAAACGAACTAACTAATGTTTAAGTAAAAGAA |
| *pAGT572_Nemo_for* | GTAATAAAAGTATCAACAAAAAATTGTTAATATACCTC |
| *pGAP_for* | GGTTTCTCCTGACCCAAAGACTTTAAA |
| *pCat1_for* | TAAGCTGTAGACCCAGCACTTCAAT |
| *tGAP_rev* | TCATTATGGCTGTATCTACTTTAGCGTA |

**Supplementary Table 2 Employed signal peptides, their origins and amino acid sequences**.

| ***Naturally secreted protein*** | ***Kingdom*** | ***Genus*** | ***Species*** | ***Abbreviation*** | ***Sequence*** |
|---|---|---|---|---|---|
| *Mating pheromone alpha factor* | Fungi | Yeast | *Saccharomyces cerevisiae* | *Sce*–Prepro | **MRFPSIFTAVLFAASSALAAPVNTTTEDETAQIPAEAVIGYLDLEGDFDVAVLPFSNSTNNGLLFINTTIASIAAKEEGVSLDKREA** |
| *Inulinase* | Fungi | Yeast | *Kluyveromyces marxianus* | *Kma*–Inulinase | **MKLAYSLLLPLAGVSAA** |
| *Invertase 2* | Fungi | Yeast | *Saccharomyces cerevisiae* | *Sce*–Invertase2 | **MLLQAFLFLLAGFAAKISA** |
| *Acid Phosphatase 5* | Fungi | Yeast | *Saccharomyces cerevisiae* | *Sce*–Acid Phosphatase | **MFKSVVYSILAASLANAA** |
| *Gma UPO* | Fungi | Basidiomycete | *Galerina marginata* | *Gma*–UPO | **MRGTPIFASLIALFAHAAIAFPAYGSLAGLTREQLDEILPTLEIRA** |
| *Serum Albumin* | Animal | Homo | *Homo sapiens* | *Hsa*–Serum Albumin | **MKWVTFISLLFLFSSAYSA** |
| *Glucoamylase* | Fungi | Ascomycete | *Aspergillus awamori* | *Aaw*–Glucoamylase | **MSFRSLLALSGLVCSGLA** |
| *Killer Protein K1 Toxin* | Fungi | Yeast | *Saccharomyces cerevisiae* | *Sce*–Killer Protein | **MTKPTQVLVRSVSILFFITLLHLVVA** |
| *Mro UPO* | Fungi | Basidiomycete | *Marasmius rotula* | *Mro*–UPO | **MKLAISSSLIALVSVTTALANSQDVVDFGA** |

| | | | | | |
|---|---|---|---|---|---|
| *Cfo CPO* | Fungi | Ascomycete | *Caldariomyces fumago* | *Cfo*–CPO | **MFSKVLPFVGAVAALPHSVRA** |
| *Cgl UPO* | Fungi | Ascomycete | *Chaetomium globosum* | *Cgl*–UPO | **MRTSLLPALAAVSPVLA** |
| *Cci UPO* | Fungi | Basidiomycete | *Coprinopsis cinerea* | *Cci*–UPO | **MISTSKHLFVLLPLFLVSHLSLVLGFPAYASLGGLTERQVEEYTSKLPIVA** |
| *α Amylase* | Fungi | Ascomycete | *Aspergillus niger* | *Ani*–α Amylase | **MVAWWSLFLYGLQVAAPALA** |
| *α Galactosidase* | Fungi | Yeast | *Saccharomyces cerevisiae* | *Sce*–α Galactosidase | **MFAFYFLTACISLKGVFGA** |
| *Lysozyme g2* | Animal | Bird | *Gallus gallus* | *Gga*–Lysozym | **MLGKNDPMCLVLVLLGLTALLGICQGA** |
| *Aae UPO* (variant PaDa-I)* | Fungi | Basidiomycete | *Agrocybe aegerita* | *Aae*–UPO* | **MKYFPLFPTLVYAVGVVAFPDYASLAGLSQQELDAIIPTLEARA** |
| *Mth UPO* | Fungi | Ascomycete | *Myceliophthora thermophila* | *Mth*–UPO | **MRASVLPVLIAISPALA** |

**Supplementary Table 3 Employed C-terminal Tags, their amino acid sequence and purpose. All sequences are terminated by introduction of a stop codon (*), terminating the open reading frame of tripartite signal peptide-gene-C-terminal Tag constructs.**

| *Name* | *Sequence* | *Purpose* |
|---|---|---|
| *GFP11* | SDGGSGGGSTSRDHMVLHEYVNAAGIT* | Protein detection (split GFP) |
| *OctaHis-GFP11* | SGGSGGGHHHHHHHHDGGSGGGSTSRDHMVLHEYVNAAGIT* | Protein purification (His Tag)<br>Protein detection (split GFP) |
| *TEV-His-GFP11* | SGSENLYFEGMGSHHHHHHSGMSDGGSGGGSTSRDHMVLHEYVNAAGIT* | Protein purification (His Tag)<br>Protein detection (split GFP) |
| *His2-GFP11* | SGGSGGGHHHHHHGGSGGSHHHHHHDGGSGGGSTSRDHMVLHEYVNAAGIT* | Protein purification (His Tag)<br>Protein detection (split GFP) |
| *Strep II* | SGGSAWSHPQFEK* | Protein purification (Strep Tag) |
| *TwinStrep* | SGGSAWSHPQFEKGGGSGGGSGGSAWSHPQFEK* | Protein purification (Strep Tag) |
| *TwinStrep-GFP11* | SGGSAWSHPQFEKGGGSGGGSGGSAWSHPQFEKDGGSGGGSTSRDHMVLHEYVNAAGIT* | Protein detection (split GFP)<br>Protein purification (Strep Tag) |

**Supplementary Table 4 Protein identification by MS: Analysis of the different unspecific peroxygenases by tryptic protein digest and MS analysis**.

| *Protein Name* | *SUM PEP Score* | *Sequence Coverage (%)* | *MW (kDa)†* | *calc. pI†* |
|---|---|---|---|---|
| *Aae*UPO* | 283.796 | 55 | 46.6 | 5.92 |
| *Gma*UPO | 155.372 | 56 | 43.6 | 6.18 |
| *Mro*UPO | 379.202 | 76 | 34.4 | 5.91 |
| *Mwe*UPO | 215.479 | 65 | 34.3 | 5.91 |
| *Cgl*UPO | 34.652 | 27 | 35.5 | 5.58 |
| *Mth*UPO | 359.253 | 62 | 35.6 | 6.35 |
| *Tte*UPO | 271.531 | 57 | 42.0 | 5.86 |

**†including the respective, attached signal peptide**

#### Supplementary Table 5 Measurement parameters for achiral and chiral GC-MS.

| Substrate | GC-MS | Column | Products | Internal Standard | Temperature program |
|---|---|---|---|---|---|
| Napthalene | Achiral | SH-Rxi-5Sil MS | 1,4-Naphthoquinone (*m/z* 158)<br>1-Naphthol (*m/z* 144) | Ethyl benzoate (*m/z* 150) | 50 °C<br>9 °C/min to 190 °C<br>55 °C/min to 300 °C hold 2 min |
| Styrene | Chiral | Lipodex E | Styrene oxide (*m/z* 119) | Ethyl benzoate (*m/z* 150) | 80 °C hold 30 min<br>50 °C/min to 200 °C hold 5 min |
| Phenylethane | Chiral | Lipodex E | Phenylethyl alcohol (*m/z* 122) | Ethyl benzoate (*m/z* 150) | 70 °C hold 40 min<br>10 °C/min to 170 °C<br>50 °C/min to 200 °C hold 5 min |
| Phenylpropane | Chiral | Lipodex E | 1-Phenyl-1-propanol (*m/z* 136) | Ethyl benzoate (*m/z* 150) | 70 °C hold 35 min<br>5 °C/min to 110 °C<br>100 °C/min to 200 °C hold 5 min |
| Phenylbutane | Chiral | Lipodex E | 1-Phenyl-1-butanol (*m/z* 150) | Ethyl benzoate (*m/z* 150) | 80 °C hold 40 min<br>1 °C/min to 110 °C<br>100 °C/min to 200 °C hold 5 min |
| Phenylpropane | Chiral | Lipodex E | 1-Phenyl-1-pentanol (*m/z* 164) | Ethyl benzoate (*m/z* 150) | 90 °C hold 60 min<br>1 °C/min to 110 °C<br>100 °C/min to 200 °C hold 5 min |
| *N*-phenethyl phthalimide | Achiral | OPTIMA 5MS Accent | *N*-(2-hydroxy-2-phenylethyl)phthalimide<br>*N*-Phthaloyl-2-oxo-phenylethyl amine | Ethyl benzoate | 50°C<br>10°C/min to 300°C hold 3 min |

#### Supplementary Table 6 Protein coverage of the protein digest and MS analysis (see above)

Signal Peptide

**NDEAHP** – Detected peptide fragments

PAP230120_1 *Aae*UPO*

MRGTPIFASL IALFAHAAIA FPAYGSLAGL TREQLDEILP TLEIR**AEPGL PPGPLENSSA KLVNDEAHPW KPLRPGDIRG PCPGLNTLAS HGYLPR**NGVA TPAQIINAVQ EGFNFDNQAA IFATYAAHLV DGN**LITDLLS IGR**K**TRLTGP DPPPPASVGG LNEHGTFEGD ASMTRGDAFF GNNH**DFNETL F**EQLVDYSNR** FGGGKYNLTV AGELRFK**RIQ DSIATNPNFS FVDFR**FFTAY **GETTFPANLF VDGRRDDGQL DMDAARSFFQ FSRMPDDFFR** APSPR**SGTGV EVVVQAHPMQ PGR**NVGKINS YTVDPTSSDF STPCLMYEKF VNITVK**SLYP NPTVQLRKAL NTNLDFLFQG VAA**GCTQVFP YGR**DSGGSAW SHPQ**FEK**GGG SGGGSGGSAW SHPQ**FEKDGG SGGGSTSRDH MVLHEYVNAA GIT

PAP230120_2 *Gma*UPO

MVAWWSLFLY GLQVAAPALA **EPAKPPGPLK** DTSAKLVNDK **AHTWKPLTPT DIRGPCPGLN TLASHG**WLPR NGIASPSEII TAVQEGFNMD NSLAIF**VTYA AHLVDGNILT DK**LSIGGK**TA LTGPNPPAPA IVGGLNTHAV FEGDTSMTR**G DFFFGNNHDF NETLF**DEFVD FSNR**FGGGKY NLTVAGEFRW QR**IQDSIATN PNFSFVSPR**Y FTAY**AESTFP INFFIDGRQN DGQLNLTVAR** GFFQN**SRMPD GFHR**ANGTRG TEGIDVIAEA HPIEPGSNVG GVNNYVVDPT SADFNTFCLL YENFVNKTIK **GLYPNPTGAL RKALNTNLGF FFSGISDTGC TQVFPYGKGS GGSAWSHPQF EKGGGSGGGS GGSAWSHPQF EK**DGGSGGGS TSR**DHMVLHE YVNAAGIT**

PAP230120_3 *Mro*UPO

MKLAISSSLI ALVSVTTA**LA NSQDVVDFGA SAHPWK**APGP NDSR**GPCPGL NTLANHGFLP R**NGRNISVPM IVK**AGFEGYN VQSDILILAG KIGMLTSREA DTISLEDLKL HGTIEHDASL SRED**VAIGDN LHFNEAIFTT LANSNPGADV YNISSAAQVQ H**DRLADSLAR NPNVTNTDLT ATIRSSESAF FLTVMSAGDP LRGEAPKKFV NVFFR**EER**MP IKEGWKRSTT PITIPLLGPI IERITELSDW KPTGDNCGAI VLSPELSGGS AWSHPQFEKG GGSGGGSGGS AWSHPQFEKD GGSGGGSTSR DHMVLHEYVN AAGIT**

PAP230129_4 *Mwe*UPO

MKLAISSSLI ALVSVTTALA **NSQDVVDFGA SAHPWK**APGP NDSR**GPCPGL NTLANHGFLP R**NGRNISVPM IVK**AGFEGYN VQSDVLITAG K**VGMLTSR**EA DTISLEDLKL HGTIEHDASL SR**EDAAIGDN LHFNEAIFTT LANSNPGADV YNISSAAQVQ H**DRLADSLAR** NPNVTNTDVT ATIR**ASESAF YLTVMSAGDP LRGEAPKKFV NVCFR**EERMP VKEGWKR**STT PINIPLLVPI IERIIELSDW KPTGDNCGAI VLSPDLSGGS AWSHPQFEKG GGSGGGSGGS AWSHPQFEKD GGSGGGSTSR DHMVLHEYVN AAGIT**

PAP230120_5 *Cgl*UPO

MVAWWSLFLY GLQVAAPALA GFDTWAPPGP YDVRGPCPML NTLTNHGFFP HDGQDIDRET TENALFDALH VNKTLASFLF DFALTTNPIA NSTTFSLNDL GNHNVLEHDA SLSRADAYHG SVLAFNHTIF EETK**SYWTDE TVTLK**MAADA RYYRIKSSQA TNPTYQMSEL GDAFTYGESA AYVVLFGDKE SQTVPRSWVE WLFEK**EQLPQ HLGWKRPATS FELNDLDKFM ALIQNYTQEI EEPSCESR**KQ RRKPRGPSHF GFSGGSAWSH PQFEK**GGGSG GGSGGSAWSH PQFEK**DGGSG GGSTSR**DHMV LHEYVNAAGI T**

#### PAP230110_6 *Mth*UPO

MFAFYFLTAC ISLKGVFG**AG FDTWSPPGPY DVRAPCPMLN TLANHGFLPH DGKDITR**EQT ENALFEALHI NKTLASF**LFD FALTTNPK**NT STF**SLNDLGN HNILEHDASL SR**ADAYFGNV LQFNQTVFDE TK**TYWEGDTI DLR**MAAKARL GRIK**TSQATN PTYSMSELGD AFTYGESAAY VVVLGDKESR** TVKRSWVEWF FEH**EQLPQHL GWKRPAASFE EEDLNSSMEE IEK**YTKELEG SNSTSGSQKH RRRLPRR**RAH FGFSGGSAWS HPQFEKGGGS GGGSGGSAWS HPQFEK**DGGS GGGSTSR**DHM VLHEYVNAAG IT**

#### PAP230120_7 *Tte*UPO

MRFPSIFTAV LFAASSALAA PVNTTTEDET AQIPAEAVIG YLDLEGDFDV AVLPFSNSTN NGLLFINTTI ASIA**AKEEGV SLDKRE**AGFD **SWHPPAPGDR RGPCPMLNTL ANHGFLPHNG RNITKEITVN ALNSALNVNK** TLGELLFNF**A VTTNPQPNAT FFDLDHLSRH NILEHDASLS R**ADYYFGHDD HTFNQTVFDQ TKSYWK**TPII DVQQAANARL** ARVLTSNATN PTFVLSQIGE AF**SFGETAAY ILALGDR**VSG TVP**RQWVEYL FENERLPLEL GWR**R**AKEVIS NSDLDQLTNR VINATGALAN ITR**KIKVRDF HAGR**FPGEGS GGSAWSHPQF EKGGGSGGGS GGSAWSHPQF EK**DGGSGGGS TSR**DHMVLHE YVNAAGIT**

**Supplementary Table 7 Enantiomeric excess determination for UPO catalysed *N*-Phthaloyl-phenylethyl amine conversion by chiral HPLC analysis.**

| Enzyme | Retention time S-Enantiomer [min] | Retention time R-Enantiomer [min] | Area S [mAU*s] | Area R [mAU*s] | ee |
|---|---|---|---|---|---|
| *Cgl*UPO | 13.45 | 18.22 | 7.294E04 | 477.713 | 98.7 % |
| *Tte*UPO | 12.91 | 16.76 | 33.33 | 18.73 | 28.0 % |
| *Mth*UPO | 13.45 | 17.75 | 2683.68945 | 19.33951 | 98.6 % |

**Supplementary Table 8 Primary data NBD conversion/ split GFP assay *Aae*UPO* signal peptide library (Primary data for 1c)**

#### NBD

| **Construct** | *Aae*-UPO* | *Sce*-Prepro | *Kma*-Inulinase | *Sce*-Invertase 2 | *Sce*-Acid Phosphatase | *Gma*-UPO | *Hsa*-Serum Albumin | *Aaw*-Glucoamylase | *Sce*-Killer Protein |
|---|---|---|---|---|---|---|---|---|---|
| | 27,889018 | 2,1547257 | 2,91738402 | 3,81613218 | 2,79263844 | 65,14925184 | 3,64885788 | 2,3616933 | 3,1215162 |
| | 28,214256 | 3,2122419 | 3,51843888 | 3,5354514 | 4,01742894 | 59,94956112 | 3,5609676 | 2,21426436 | 2,56057002 |
| | 26,922692 | 3,3681771 | 2,51195508 | 3,71690022 | 2,57648862 | 57,6247257 | 3,3681771 | 2,6196933 | 2,94573564 |
| | 27,322982 | 2,49778056 | 2,53180302 | 3,12718704 | 3,75092268 | 61,1601771 | 3,19651938 | 1,69542894 | 2,8635162 |
| | 25,912581 | 2,85784536 | 2,99960346 | 4,19037408 | 4,5277581 | 65,61705486 | 3,13852872 | 1,99432968 | 3,3284838 |
| Average | 27,252306 | 2,818154124 | 2,895836892 | 3,677208984 | 3,533047356 | 61,90015412 | 3,382610136 | 2,177081916 | 2,963964372 |
| SD | 0,8048917 | 0,447701315 | 0,368442918 | 0,348377042 | 0,739561305 | 3,066061132 | 0,198551286 | 0,315200477 | 0,257233791 |
| SD (%) | 2,9534812 | 15,88633181 | 12,72319306 | 9,473952751 | 20,93267455 | 4,953236668 | 5,869765595 | 14,47811746 | 8,678707267 |

| **Construct** | *Mro*-UPO | *Cfo*-CPO | *Cgl*-UPO | *Cci*-UPO | *Ani*-α Amylase | *Sce*-α Galactosidase | *Gga*-Lysozym | *Mth*-UPO | Negative control |
|---|---|---|---|---|---|---|---|---|---|
| | 2,46092268 | 1,927913 | 2,15189 | 9,588219 | 16,7184 | 6,5532 | 4,64869044 | 3,06764838 | 0,83177394 |
| | 2,54314212 | 2,163232 | 2,699077 | 8,50931 | 16,90603566 | 6,4359261 | 3,43843566 | 2,57149374 | 1,27865832 |
| | 2,75578056 | 2,174318 | 2,636703 | 8,31229 | 17,20625478 | 5,8448739 | 3,70581912 | 3,84164838 | 0,99577422 |
| | 2,39180964 | 1,638914 | 2,713788 | 8,603126 | 16,3712739 | 6,03250956 | 3,87469044 | 4,09397754 | 0,46780302 |
| | 2,90037408 | 1,650699 | 2,801142 | 8,368581 | 17,10305478 | 7,20054522 | 4,01541912 | 3,4447257 | 0,75906954 |
| Average | 2,610405816 | 1,911015 | 2,60052 | 8,676305 | 16,86100382 | 6,413410956 | 3,936610956 | 3,40389875 | 0,866615808 |
| SD | 0,189667768 | 0,234534 | 0,230378 | 0,467339 | 0,296643221 | 0,470733155 | 0,404609828 | 0,54322421 | 0,267691136 |
| SD (%) | 7,265834565 | 12,27274 | 8,858924 | 5,386388 | 1,759344961 | 7,339825221 | 10,27812585 | 15,9588828 | 30,88925149 |

#### split GFP

| **Construct** | *Aae*-UPO* | *Sce*-Prepro | *Kma*-Inulinase | *Sce*-Invertase 2 | *Sce*-Acid Phosphatase | *Gma*-UPO | *Hsa*-Serum Albumin | *Aaw*-Glucoamylase | *Sce*-Killer Protein |
|---|---|---|---|---|---|---|---|---|---|
| | 753 | 351 | 149 | 329 | 182 | 1578 | 308 | 192 | 593 |
| | 793 | 228 | 202 | 356 | 231 | 1517 | 360 | 203 | 470 |
| | 604 | 330 | 204 | 360 | 152 | 1456 | 423 | 249 | 427 |
| | 594 | 294 | 205 | 389 | 130 | 1507 | 305 | 191 | 395 |
| | 649 | 288 | 180 | 448 | 149 | 1551 | 355 | 154 | 333 |
| Average | 678,6 | 298,2 | 188 | 376,4 | 168,8 | 1521,8 | 350,2 | 197,8 | 443,6 |
| SD | 80,276024 | 42,0685155 | 21,56849554 | 40,53936359 | 35,27832196 | 41,43138907 | 42,99488342 | 30,48540634 | 87,04849223 |
| SD (%) | 11,829653 | 14,1074834 | 11,47260401 | 10,77028788 | 20,89947983 | 2,722525238 | 12,27723684 | 15,41223779 | 19,62319482 |

| **Construct** | *Mro*-UPO | *Cfo*-CPO | *Cgl*-UPO | *Cci*-UPO | *Ani*-α Amylase | *Sce*-α Galactosidase | *Gga*-Lysozym | *Mth*-UPO | Negative control |
|---|---|---|---|---|---|---|---|---|---|
| | 292 | 170 | 217 | 889 | 622 | 644 | 706 | 270 | 89 |
| | 311 | 121 | 190 | 806 | 736 | 640 | 768 | 298 | 60 |
| | 285 | 142 | 261 | 954 | 759 | 742 | 796 | 374 | 78 |
| | 302 | 191 | 258 | 979 | 611 | 641 | 804 | 281 | 36 |
| | 282 | 217 | 200 | 933 | 667 | 685 | 703 | 252 | 2 |
| Average | 294,4 | 168,2 | 225,2 | 912,2 | 679 | 670,4 | 755,4 | 295 | 53 |
| SD | 10,7814656 | 34,11393 | 29,32166 | 60,75986 | 59,44072678 | 39,56058645 | 43,25551988 | 42,237424 | 31,17691454 |
| SD (%) | 3,6621826 | 20,28176 | 13,02028 | 6,660804 | 8,75415711 | 5,901042131 | 5,726174196 | 14,317771 | 58,82436705 |

#### Supplementary Table 9 Primary data DMP conversion and split GFP assay signal peptide library screening of various UPOs (Primary data for 2a)

##### DMP

| **Enzyme** | ***Gma* UPO** | ***Gma* UPO** | ***Gma* UPO** | ***Mwe* UPO** | ***Mwe* UPO** | ***Mwe* UPO** | ***Mro* UPO** | ***Mro* UPO** | ***Mro* UPO** |
|---|---|---|---|---|---|---|---|---|---|
| **Construct** | Negative control | *Sce*-Invertase 2 | *Ani*-α Amylase | Negative control | *Sce*-Prepro | *Mro*-UPO | Negative control | *Ani*-α Amylase | *Mro*-UPO |
| | n.d. | n.d. | n.d. | n.d. | n.d. | n.d. | 0,029257 | 2,301843 | 5,670507 |
| | n.d. | n.d. | n.d. | n.d. | n.d. | n.d. | 0,153038 | 3,576037 | 5,903226 |
| | n.d. | n.d. | n.d. | n.d. | n.d. | n.d. | 0,173526 | 1,788018 | 5,313364 |
| | n.d. | n.d. | n.d. | n.d. | n.d. | n.d. | 0,248848 | 1,645161 | 6,232719 |
| | n.d. | n.d. | n.d. | n.d. | n.d. | n.d. | | | |
| | n.d. | n.d. | n.d. | n.d. | n.d. | n.d. | | | |
| | n.d. | n.d. | n.d. | n.d. | n.d. | n.d. | | | |
| | n.d. | n.d. | n.d. | n.d. | n.d. | n.d. | | | |
| Average | n.d. | n.d. | n.d. | n.d. | n.d. | n.d. | 0,15116725 | 2,32776475 | 5,779954 |
| SD | n.d. | n.d. | n.d. | n.d. | n.d. | n.d. | 0,07890984 | 0,760941847 | 0,335364356 |
| SD (%) | n.d. | n.d. | n.d. | n.d. | n.d. | n.d. | 52,20035423 | 32,68980886 | 5,802197665 |

| **Enzyme** | ***Cgl* UPO** | ***Cgl* UPO** | ***Cgl* UPO** | ***Mth* UPO** | ***Mth* UPO** | ***Mth* UPO** | ***Tte* UPO** | ***Tte* UPO** | ***Tte* UPO** |
|---|---|---|---|---|---|---|---|---|---|
| **Construct** | Negative control | *Ani*-α Amylase | *Cgl*-UPO | Negative control | *Sce*-Acid Phosphatase | *Sce*-α Galactosidase | Negative control | *Sce*-Prepro | *Cci*-UPO |
| | 0,064399501 | 0,611693683 | 1,70773008 | 0,064399501 | 1,234390424 | 2,481110322 | 0,26012928 | 130,120258 | 155,7058 |
| | 0,077320214 | 0,693623557 | 1,39861584 | 0,077320214 | 1,037651775 | 2,28238296 | 0,11148384 | 130,008774 | 153,3646 |
| | 0,08014067 | 0,625613763 | 1,18972416 | 0,08014067 | 1,074905298 | 2,441205789 | 0,0371616 | 142,141935 | 153,216 |
| | 0,115205971 | 0,603738831 | 1,28262768 | 0,115205971 | 1,296169889 | 2,303462589 | 0,26012928 | 126,366968 | 149,0539 |
| | 0,092536647 | 0,618322119 | 1,55289168 | 0,092536647 | 1,502188895 | 2,298424759 | 0,16722624 | 126,738581 | 141,3987 |
| | | | | | | | 0,35303232 | 136,307613 | 150,1316 |
| | | | | | | | -0,0371616 | 137,403871 | 147,809 |
| | | | | | | | 0,11148384 | 135,415742 | 155,1484 |
| Average | 0,085920601 | 0,630598391 | 1,42631789 | 0,085920601 | 1,229061256 | 2,361317284 | 0,1579356 | 133,062968 | 150,7285 |
| SD | 0,017157721 | 0,032331251 | 0,18579807 | 0,017157721 | 0,167052282 | 0,082783825 | 0,120416439 | 5,24590668 | 4,418351 |
| SD (%) | 19,96927509 | 5,127074729 | 13,0264139 | 19,96927509 | 13,59185974 | 3,505832326 | 76,24401304 | 3,94242423 | 2,931331 |

n.d.- no detectable DMP conversion above background level (negative control)

##### split GFP

| **Enzyme** | ***Gma* UPO** | ***Gma* UPO** | ***Gma* UPO** | ***Mwe* UPO** | ***Mwe* UPO** | ***Mwe* UPO** | ***Mro* UPO** | ***Mro* UPO** | ***Mro* UPO** |
|---|---|---|---|---|---|---|---|---|---|
| **Construct** | Negative control | *Sce*-Invertase 2 | *Ani*-α Amylase | Negative control | *Sce*-Prepro | *Mro*-UPO | Negative control | *Ani*-α Amylase | *Mro*-UPO |
| | 447 | 1579 | 1938 | 237 | 557 | 860 | 176 | 690 | 577 |
| | 449 | 1847 | 1405 | 395 | 801 | 873 | 148 | 865 | 718 |
| | 422 | 2143 | 1913 | 507 | 777 | 742 | 209 | 957 | 653 |
| | 342 | 1328 | 1858 | 483 | 708 | 823 | 217 | 871 | 649 |
| | 496 | 1574 | 1783 | 701 | 852 | 966 | 115 | 878 | 529 |
| | 369 | 1725 | 1769 | 507 | 948 | 790 | 169 | 800 | 770 |
| | | | | 352 | 853 | 727 | 224 | 759 | 459 |
| | | | | 370 | 919 | 766 | 183 | 844 | 593 |
| | | | | 424 | 980 | 995 | 222 | 888 | 561 |
| | | | | 519 | 789 | 1028 | 110 | 781 | 528 |
| | | | | 510 | 808 | 934 | 110 | 747 | 683 |
| | | | | 471 | 711 | 891 | | | |
| Average | 420,8333333 | 1699,333333 | 1777,666667 | 456,3333333 | 808,5833333 | 866,25 | 171,1818182 | 825,4545455 | 610,909091 |
| SD | 51,68951108 | 254,0935961 | 177,6979335 | 109,5379183 | 111,5208937 | 95,97753643 | 42,90543128 | 73,54820761 | 88,3273401 |
| SD (%) | 12,2826561 | 14,95254586 | 9,996133515 | 24,00392659 | 13,79213361 | 11,07965789 | 25,06424557 | 8,910025151 | 14,4583444 |

| **Enzyme** | ***Cgl* UPO** | ***Cgl* UPO** | ***Cgl* UPO** | ***Mth* UPO** | ***Mth* UPO** | ***Mth* UPO** | ***Tte* UPO** | ***Tte* UPO** | ***Tte* UPO** |
|---|---|---|---|---|---|---|---|---|---|
| **Construct** | Negative control | *Ani*-α Amylase | *Cgl*-UPO | Negative control | *Sce*-Acid Phosphatase | *Sce*-α Galactosidase | Negative control | *Sce*-Prepro | *Cci*-UPO |
| | 143 | 873 | 849 | 143 | 766 | 2913 | 463 | 23586 | 25138 |
| | 164 | 668 | 969 | 164 | 789 | 2780 | 508 | 20994 | 28146 |
| | 151 | 793 | 997 | 151 | 743 | 2870 | 555 | 20363 | 28977 |
| | 178 | 549 | 877 | 178 | 701 | 2822 | 522 | 23449 | 24324 |
| | 214 | 636 | 1147 | 214 | 802 | 3015 | 473 | 22536 | 23760 |
| | | | | | | | 504 | 21380 | 23802 |
| | | | | | | | 531 | 20566 | 25972 |
| | | | | | | | 511 | 18680 | 24927 |
| Average | 170 | 703,8 | 967,8 | 170 | 760,2 | 2880 | 508,375 | 21444,25 | 25630,75 |
| SD | 25,00399968 | 115,262136 | 105,1901 | 25,00399968 | 35,80726183 | 80,96665981 | 27,8564961 | 1564,70403 | 1834,735 |
| SD (%) | 14,70823511 | 16,37711509 | 10,86899 | 14,70823511 | 4,710242283 | 2,811342354 | 5,479517305 | 7,29661343 | 7,158335 |

#### Supplementary Table 10 Primary data NBD conversion regarding the pH profile accession of various UPOs (Primary data for 2c)

| ***Mro* UPO** | | | | | | | | | |
|---|---|---|---|---|---|---|---|---|---|
| **pH value** | 3 | 4 | 5 | 6 | 7 | 8 | 9 | 10 | 11 |
| | n.d. | 0,200353 | 1,721818 | 9,365455 | 12,56909 | 7,629091 | 4,94 | 3,161818 | 2,667273 |
| | n.d. | 0,405195 | 1,721818 | 8,134545 | 11,90909 | 6,556364 | 4,041818 | 2,878182 | 2,645455 |
| | n.d. | 0,458182 | 1,512727 | 8,123636 | 10,30546 | 7,518182 | 4,161818 | 3,343636 | 1,234916 |
| Average | n.d. | 0,354577 | 1,652121 | 8,541212 | 11,59455 | 7,234546 | 4,381212 | 3,127879 | 2,182548 |
| SD | n.d. | 0,111177 | 0,098566 | 0,582845 | 0,950514 | 0,48168 | 0,398148 | 0,19153 | 0,670136 |
| SD (%) | n.d. | 31,35496 | 5,966055 | 6,823912 | 8,197943 | 6,65805 | 9,087628 | 6,123328 | 30,70431 |

| ***Cgl* UPO** | | | | | | | | | |
|---|---|---|---|---|---|---|---|---|---|
| **pH value** | 3 | 4 | 5 | 6 | 7 | 8 | 9 | 10 | 11 |
| | n.d. | 0,023077 | -0,22 | 2,572727 | 13,88364 | 16,78364 | 6,98 | 3,938182 | 0,577489 |
| | n.d. | 0,017857 | 0,66 | 5,130909 | 13,93273 | 21,00182 | 6,189091 | 1,6 | 0,692727 |
| | n.d. | -0,04048 | 0 | 4,427273 | 12,66546 | 18,40727 | 6,254545 | 2,085455 | 1,581818 |
| Average | n.d. | 0,000153 | 0,146667 | 4,043636 | 13,49394 | 18,73091 | 6,474545 | 2,541212 | 0,950678 |
| SD | n.d. | 0,028808 | 0,373928 | 1,079029 | 0,58617 | 1,737205 | 0,358408 | 1,007492 | 0,448756 |
| SD (%) | n.d. | 18869,7 | 254,951 | 26,68463 | 4,343947 | 9,274535 | 5,535646 | 39,64611 | 47,20381 |

| ***Mth* UPO** | | | | | | | | | |
|---|---|---|---|---|---|---|---|---|---|
| **pH value** | 3 | 4 | 5 | 6 | 7 | 8 | 9 | 10 | 11 |
| | n.d. | 11,48571 | 55,34026 | 65,56364 | 186,3584 | 119,5481 | 79,51169 | 43,26234 | 22,61299 |
| | n.d. | 13,41818 | 47,15844 | 57,49091 | 159,7558 | 96,56104 | 70,42597 | 40,28571 | 16,97143 |
| | n.d. | 11,73507 | 42,85714 | 48,43636 | 196,8623 | 93,13247 | 65,25195 | 30,96623 | 10,47273 |
| Average | n.d. | 12,21299 | 48,45195 | 57,16364 | 180,9922 | 103,0805 | 71,72987 | 38,17143 | 16,68571 |
| SD | n.d. | 0,85826 | 5,177639 | 6,996008 | 15,61666 | 11,72813 | 5,894074 | 5,23776 | 4,960356 |
| SD (%) | n.d. | 7,027437 | 10,68613 | 12,23856 | 8,628363 | 11,37764 | 8,217042 | 13,72168 | 29,72816 |

| ***Tte* UPO** | | | | | | | | | |
|---|---|---|---|---|---|---|---|---|---|
| **pH value** | 3 | 4 | 5 | 6 | 7 | 8 | 9 | 10 | 11 |
| | 9,75 | 27,87 | 56,01 | 117,81 | 98,94 | 56,67 | 35,55 | 24,6 | 11,13 |
| | 7,83 | 26,94 | 59,58 | 103,8 | 107,43 | 47,79 | 73,56 | 26,64 | 12,3 |
| | 8,58 | 33,75 | 62,97 | 121,56 | 116,07 | 52,71 | 38,16 | 28,98 | 15,21 |
| Average | 8,72 | 29,52 | 59,52 | 114,39 | 107,48 | 52,39 | 49,09 | 26,74 | 12,88 |
| SD | 0,790063 | 3,015062 | 2,841725 | 7,643154 | 6,993383 | 3,6323 | 17,33568 | 1,789525 | 1,715401 |
| SD (%) | 9,060359 | 10,21363 | 4,774403 | 6,681663 | 6,506683 | 6,933192 | 35,31408 | 6,692315 | 13,31833 |

n.d.- no detectable NBD conversion above background level (buffer control)

#### Supplementary Table 11 Primary data NBD conversion *Pichia pastoris* construct comparison (Primary data for 4b)

| | | | | |
|---|---|---|---|---|
| **GAP promoter (1.5 % glucose)** | | | | |
| | episomal *Mth* UPO | episomal (-) | integrative *Mth* UPO | integrative (-) |
| | 1,496364 | 0,969091 | 12,265455 | 0,796364 |
| | 0,789091 | 0,610909 | 10,270909 | 0,56 |
| | 0,898182 | 0,861818 | 10,152727 | 0,674545 |
| | 1,336364 | 0,836364 | 10,165455 | 0,485455 |
| | 0,705455 | 1,110909 | 10,285455 | 0,610909 |
| | 0,54 | 0,66 | 8,436364 | 0,549091 |
| Average | 0,960909333 | 0,841515167 | 10,2627275 | 0,612727333 |
| SD | 0,34243144 | 0,170984566 | 1,108201741 | 0,100478154 |
| SD (%) | 35,63618632 | 20,31865525 | 10,79831596 | 16,39851022 |
| **CAT1 promoter (0.5 % glycerol; 1.5 %methanol)** | | | | |
| | episomal *Mth* UPO | episomal (-) | integrative *Mth* UPO | integrative (-) |
| | 2,910909 | 0,843636 | 15,810909 | 0,581818 |
| | 2,354545 | 0,450909 | 16,934545 | 0,601818 |
| | 2,570909 | 0,914545 | 15,289091 | 0,558182 |
| | 2,434545 | 0,690909 | 15,025455 | 0,72 |
| | 2,289091 | 0,494545 | 14,212727 | 0,643636 |
| | 2,365455 | 0,650909 | 13,945455 | 0,647273 |
| Average | 2,487575667 | 0,674242167 | 15,20303033 | 0,6254545 |
| SD | 0,208556381 | 0,168043009 | 0,997396224 | 0,052810799 |
| SD (%) | 8,383921112 | 24,92324234 | 6,560509332 | 8,44358767 |

#### Supplementary Table 12 Primary data NBD conversion *Pichia pastoris* $P_{CAT1}$ construct comparison; single colonies (Primary data for 4c)

| Construct | episomal *Mth* UPO | integrative *Mth* UPO | episomal (-) | integrative (-) |
|---|---|---|---|---|
| | 3,251049 | 11,964336 | 0,799301 | 0,439161 |
| | 4,848951 | 12,427273 | 0,716084 | 0,316783 |
| | 3,958741 | 12,511888 | 0,61958 | 0,393007 |
| | 3,733566 | 1,838462 | 0,409091 | 0,302797 |
| | 4,75035 | 1,612587 | 0,49021 | 0,469231 |
| | 3,941259 | 11,023077 | 0,79021 | 0,358741 |
| | 2,857343 | 2,312587 | 0,838462 | 0,311189 |
| | 3,648252 | 1,979021 | 0,397203 | 0,428671 |
| | 3,693007 | 2,081119 | 0,351748 | 0,361538 |
| | 3,296503 | 2,497203 | 0,586014 | 0,332867 |
| | 4,16993 | 2,605594 | 0,505594 | 0,487413 |
| Average | 3,831722818 | 5,713922455 | 0,591227 | 0,381945273 |
| SD | 0,576568352 | 4,758735234 | 0,166826251 | 0,062669499 |
| SD (%) | 15,0472354 | 83,28316095 | 28,21695409 | 16,40797875 |

#### Supplementary Table 13 Primary data NBD conversion *Pichia pastoris* $P_{GAP}$ and $P_{CAT1}$ constructs comparison of three different UPOs in combination with two signal peptides each (Primary data for 4d)

| **pGAP constructs** | | | | | | | |
|---|---|---|---|---|---|---|---|
| **Enzyme** | *Aae* UPO* | *Aae* UPO* | *Mth* UPO | *Mth* UPO | *Tte* UPO | *Tte* UPO | Negative control |
| **Signal Peptide** | *Aae* -UPO* | *Gma* -UPO | *Sce* -Acid Phosphatase | *Sce* -α Galactosidase | *Sce* -Prepro | *Cci* -UPO | |
| | 0,652727 | 5,267273 | 1,14 | 0,38 | 4,278182 | 4,616364 | 0,192727 |
| | 0,549091 | 5,703636 | 0,956364 | 0,727273 | 3,852727 | 4,461818 | 0,298182 |
| | 0,567273 | 4,536364 | 1,130909 | 0,470909 | 4,781818 | 3,563636 | 0,42 |
| | 0,638182 | 6,02 | 0,934545 | 0,723636 | 4,336364 | 3,296364 | 0,389091 |
| | 0,64 | 5,114545 | 0,716364 | 0,38 | 4,209091 | 4,870909 | 0,398182 |
| | 0,414545 | 5,310909 | 0,86 | 0,698182 | 3,254545 | 3,66 | 0,461818 |
| Average | 0,57696967 | 5,3254545 | 0,956363667 | 0,563333333 | 4,11878783 | 4,078182 | 0,36 |
| SD | 0,0823731 | 0,46478633 | 0,148071057 | 0,15626996 | 0,47226645 | 0,593872 | 0,0895105 |
| SD (%) | 14,2768508 | 8,72763685 | 15,48271455 | 27,74022953 | 11,4661514 | 14,56218 | 24,86402791 |
| **pCAT1 constructs** | | | | | | | |
| **Enzyme** | *Aae* UPO* | *Aae* UPO* | *Mth* UPO | *Mth* UPO | *Tte* UPO | *Tte* UPO | Negative control |
| **Signal Peptide** | *Aae* -UPO* | *Gma* -UPO | *Sce* -Acid Phosphatase | *Sce* -α Galactosidase | *Sce* -Prepro | *Cci* -UPO | |
| | 0,794406 | 4,204196 | 0,686713 | 2,145455 | 20,23986 | 1,577622 | 0,174825 |
| | 0,565734 | 4,509091 | 0,640559 | 2,34965 | 24,111888 | 1,20979 | 0,163636 |
| | 0,92028 | 3,958741 | 0,497902 | 1,967832 | 22,964336 | 1,183217 | 0,04965 |
| | 0,587413 | 4,877622 | 0,38042 | 3,296503 | 21,632867 | 1,072727 | 0,032168 |
| | 0,917483 | 4,358042 | 0,220979 | 2,952448 | 24,879021 | 0,692308 | 0,093706 |
| | 0,717483 | 5,958741 | 0,796503 | 2,275524 | 21,720979 | 0,958741 | 0,013287 |
| Average | 0,7504665 | 4,6444055 | 0,537179333 | 2,497902 | 22,5914918 | 1,115734 | 0,087878667 |
| SD | 0,14169349 | 0,65142178 | 0,194248675 | 0,469219094 | 1,57534829 | 0,268492 | 0,062532421 |
| SD (%) | 18,8807213 | 14,0259454 | 36,16086146 | 18,78452772 | 6,97319284 | 24,06413 | 71,15768106 |

#### Supplementary Table 14 Primary data NBD conversion and split GFP assay species comparison *Mth*UPO and *Tte*UPO (Primary data for 4e)

#### NBD

| **Species** | *P. pastoris* | *P. pastoris* | *P. pastoris* | *P. pastoris* | *P. pastoris* | *S. cerevisiae* | *S. cerevisiae* | *S. cerevisiae* | *S. cerevisiae* | *S. cerevisiae* |
|---|---|---|---|---|---|---|---|---|---|---|
| **Enzyme** | *Mth* UPO | *Mth* UPO | *Tte* UPO | *Tte* UPO | Negative control | *Mth* UPO | *Mth* UPO | *Tte* UPO | *Tte* UPO | Negative control |
| **Signal Peptide** | *Sce* - α Galactosidase | *Ani* - α Amylase | *Sce* -Prepro | *Sce* -Invertase 2 | - | *Sce* - Acidic Phosphatase | *Sce* - α Galactosidase | *Sce* -Prepro | *Cci* -UPO | - |
| | 1,534521982 | 1,560373703 | 2,794396888 | 3,776451794 | -0,055570317 | 1,367665324 | 2,260848245 | 9,098299725 | 10,12491882 | -0,051841238 |
| | 1,578937553 | 1,504809236 | 2,134699997 | 4,658551682 | -0,070416075 | 0,782030806 | 2,631491617 | 9,806180734 | 9,502345619 | -0,125971286 |
| | 1,434365421 | 1,223069599 | 2,423739041 | 3,813601055 | -0,185294275 | 1,389846688 | 2,271948993 | 9,131631522 | 10,69197952 | -0,092603975 |
| | 1,367633656 | 1,515859584 | 2,127236318 | 3,728202141 | -0,21873231 | 1,211887772 | 2,375681676 | 9,331705802 | 9,353994337 | -0,144516844 |
| | 1,50852016 | 1,334306967 | 2,3199386 | 4,573244613 | -0,029642928 | 1,323126596 | 2,1422252 | 9,739426805 | 10,90681092 | -0,181611037 |
| | 1,074874295 | 1,382726166 | 2,368184901 | 4,176786516 | -0,18152581 | 1,519616267 | 2,301637404 | 7,890140813 | 10,03227098 | -0,081469389 |
| Average | 1,416475511 | 1,420190876 | 2,361365958 | 4,121139634 | -0,123530286 | 1,265695575 | 2,330638856 | 9,1662309 | 10,10205337 | -0,113002295 |
| SD | 0,167430617 | 0,118039853 | 0,223369297 | 0,379494566 | 0,073593715 | 0,234667232 | 0,151215773 | 0,632414547 | 0,565726921 | 0,042855594 |
| SD (%) | 11,82022672 | 8,311548494 | 9,459325718 | 9,208486003 | -59,57544301 | 18,54057454 | 6,488168349 | 6,899395765 | 5,600118115 | -37,92453386 |

#### split GFP

| **Species** | *P. pastoris* | *P. pastoris* | *P. pastoris* | *P. pastoris* | *P. pastoris* | *S. cerevisiae* | *S. cerevisiae* | *S. cerevisiae* | *S. cerevisiae* | *S. cerevisiae* |
|---|---|---|---|---|---|---|---|---|---|---|
| **Enzyme** | *Mth* UPO | *Mth* UPO | *Tte* UPO | *Tte* UPO | Negative control | *Mth* UPO | *Mth* UPO | *Tte* UPO | *Tte* UPO | Negative control |
| **Signal Peptide** | *Sce* - α Galactosidase | *Ani* - α Amylase | *Sce* -Prepro | *Sce* -Invertase 2 | - | *Sce* - Acidic Phosphatase | *Sce* - α Galactosidase | *Sce* -Prepro | *Cci* -UPO | - |
| | 685 | 497 | 7141 | 8539 | 273 | 2003 | 4088 | 28263 | 31755 | -101 |
| | 381 | 624 | 5324 | 6953 | 225 | 2026 | 4010 | 26143 | 28928 | -193 |
| | 685 | 652 | 5266 | 6451 | 276 | 2672 | 3719 | 30688 | 28782 | -193 |
| | 560 | 634 | 7409 | 7227 | 266 | 2418 | 4157 | 28609 | 28955 | -161 |
| | 588 | 619 | 5707 | 7845 | 242 | 2502 | 3853 | 21753 | 30681 | -100 |
| | 511 | 669 | 6431 | 7205 | 225 | 2022 | 4571 | 30344 | 28453 | -200 |
| Average | 568,3333333 | 615,8333333 | 6213 | 7370 | 251,1666667 | 2273,833333 | 4066,333333 | 27633,33333 | 29592,33333 | -158 |
| SD | 104,9280177 | 55,77160767 | 844,8165481 | 665,6212637 | 21,47414466 | 267,5733025 | 268,5464992 | 3022,87174 | 1201,705409 | 42,49705872 |
| SD (%) | 18,4624078 | 9,056282707 | 13,59756234 | 9,031496115 | 8,549758989 | 11,76749846 | 6,604143762 | 10,93922222 | 4,060867371 | -26,89687261 |

**Protein sequences**

**A**: Signal Peptide- Gene overhang amino acid (A**GCA**)

**S**: Gene- C-terminal Tag overhang amino acid (T**TCG**)

**Long-type UPOs:**

***Aae*UPO*** (330 amino acids)

**A**EPGLPPGPLENSSAKLVNDEAHPWKPLRPGDIRGPCPGLNTLASHGYLPRNGVATPAQIINAVQEGFNFDNQAAI
FATYAAHLVDGNLITDLLSIGRKTRLTGPDPPPPASVGGLNEHGTFEGDASMTRGDAFFGNNHDFNETLFEQLVDY
SNRFGGGKYNLTVAGELRFKRIQDSIATNPNFSFVDFRFFTAYGETTFPANLFVDGRRDDGQLDMDAARSFFQFSR
MPDDFFRAPSPRSGTGVEVVVQAHPMQPGRNVGKINSYTVDPTSSDFSTPCLMYEKFVNITVKSLYPNPTVQLRKA
LNTNLDFLFQGVAAGCTQVFPYGRD**S**

***Gma*UPO** (331 amino acids)

**A**EPAKPPGPLKDTSAKLVNDKAHTWKPLTPTDIRGPCPGLNTLASHGWLPRNGIASPSEIITAVQEGFNMDNSLAIF
VTYAAHLVDGNILTDKLSIGGKTALTGPNPPAPAIVGGLNTHAVFEGDTSMTRGDFFFGNNHDFNETLFDEFVDFS
NRFGGGKYNLTVAGEFRWQRIQDSIATNPNFSFVSPRYFTAYAESTFPINFFIDGRQNDGQLNLTVARGFFQNSRM
PDGFHRANGTRGTEGIDVIAEAHPIEPGSNVGGVNNYVVDPTSADFNTFCLLYENFVNKTIKGLYPNPTGALRKALN
TNLGFFFSGISDTGCTQVFPYGKG**S**

***Cci*UPO** (339 amino acids)

**A**FPPPPPPEPIKDPWLKLVNDRAHPWRPLRRGDVRGPCPGLNTLASHGYLPRDGVATPAQIITAVQEGFNMEYGIAT
FVTYAAHLVDGNPLTNLISIGGKTRKTGPDPPPPAIVGGLNTHAVFEGDASMTRGDFHLGDNFNFNQTLWEQFKD
YSNRYGGGRYNLTAAAELRWARIQQSMATNGQFDFTSPRYFTAYAESVFPINFFTDGRLFTSNTTAPGPDMDSALS
FFRDHRYPKDFHRAPVPSGARGLDVVAAAYPIQPGYNADGKVNNYVLDPTSADFTKFCLLYENFVLKTVKGLYPNP
KGFLRKALETNLEYFYQSFPGSGGCPQVFPWGKSD**S**

**Short-type UPOs:**

***Mro*UPO** (238 amino acids)

**A**SAHPWKAPGPNDSRGPCPGLNTLANHGFLPRNGRNISVPMIVKAGFEGYNVQSDILILAGKIGMLTSREADTISLE
DLKLHGTIEHDASLSREDVAIGDNLHFNEAIFTTLANSNPGADVYNISSAAQVQHDRLADSLARNPNVTNTDLTATI
RSSESAFFLTVMSAGDPLRGEAPKKFVNVFFREERMPIKEGWKRSTTPITIPLLGPIIERITELSDWKPTGDNCGAIVLS
PEL**S**

***Mwe*UPO** (238 amino acids)

**A**SAHPWKAPGPNDSRGPCPGLNTLANHGFLPRNGRNISVPMIVKAGFEGYNVQSDVLITAGKVGMLTSREADTIS
LEDLKLHGTIEHDASLSREDAAIGDNLHFNEAIFTTLANSNPGADVYNISSAAQVQHDRLADSLARNPNVTNTDVTA
TIRASESAFYLTVMSAGDPLRGEAPKKFVNVCFREERMPVKEGWKRSTTPINIPLLVPIIERIIELSDWKPTGDNCGAI
VLSPDL**S**

***Cgl*UPO** (244 amino acids)

**A**GFDTWAPPGPYDVRGPCPMLNTLTNHGFFPHDGQDIDRETTENALFDALHVNKTLASFLFDFALTTNPIANSTTF
SLNDLGNHNVLEHDASLSRADAYHGSVLAFNHTIFEETKSYWTDETVTLKMAADARYYRIKSSQATNPTYQMSELG
DAFTYGESAAYVVLFGDKESQTVPRSWVEWLFEKEQLPQHLGWKRPATSFELNDLDKFMALIQNYTQEIEEPSCES
RKQRRKPRGPSHFGF**S**

***Mth*UPO** (246 amino acids)

**A**GFDTWSPPGPYDVRAPCPMLNTLANHGFLPHDGKDITREQTENALFEALHINKTLASFLFDFALTTNPKNTSTFSL
NDLGNHNILEHDASLSRADAYFGNVLQFNQTVFDETKTYWEGDTIDLRMAAKARLGRIKTSQATNPTYSMSELGD
AFTYGESAAYVVVLGDKESRTVKRSWVEWFFEHEQLPQHLGWKRPAASFEEEDLNSSMEEIEKYTKELEGSNSTSG
SQKHRRRLPRRRAHFGF**S**

***Tte*UPO** (244 amino acids)

**A**GFDSWHPPAPGDRRGPCPMLNTLANHGFLPHNGRNITKEITVNALNSALNVNKTLGELLFNFAVTTNPQPNATF
FDLDHLSRHNILEHDASLSRADYYFGHDDHTFNQTVFDQTKSYWKTPIIDVQQAANARLARVLTSNATNPTFVLSQI
GEAFSFGETAAYILALGDRVSGTVPRQWVEYLFENERLPLELGWRRAKEVISNSDLDQLTNRVINATGALANITRKIK
VRDFHAGRFPGEG**S**

**Sequence identity comparison**

| | PaDa-I | GmaUPO | CciUPO | MroUPO | MweUPO | CglUPO | MthUPO | TteUPO |
|---|---|---|---|---|---|---|---|---|
| PaDa-I | | 71.4% | 61.8% | 30.1% | 30.1% | 22.6% | 24.5% | 24.8% |
| GmaUPO | 71.4% | | 64.5% | 28.6% | 28.2% | 22.2% | 24.9% | 24.1% |
| CciUPO | 61.8% | 64.5% | | 25.8% | 25.0% | 22.1% | 22.6% | 23.6% |
| MroUPO | 30.1% | 28.6% | 25.8% | | 94.5% | 30.4% | 31.0% | 31.0% |
| MweUPO | 30.1% | 28.2% | 25.0% | 94.5% | | 30.8% | 31.0% | 31.4% |
| CglUPO | 22.6% | 22.2% | 22.1% | 30.4% | 30.8% | | 72.1% | 49.8% |
| MthUPO | 24.5% | 24.9% | 22.6% | 31.0% | 31.0% | 72.1% | | 51.8% |
| TteUPO | 24.8% | 24.1% | 23.6% | 31.0% | 31.4% | 49.8% | 51.8% | |

### V. NMR spectra

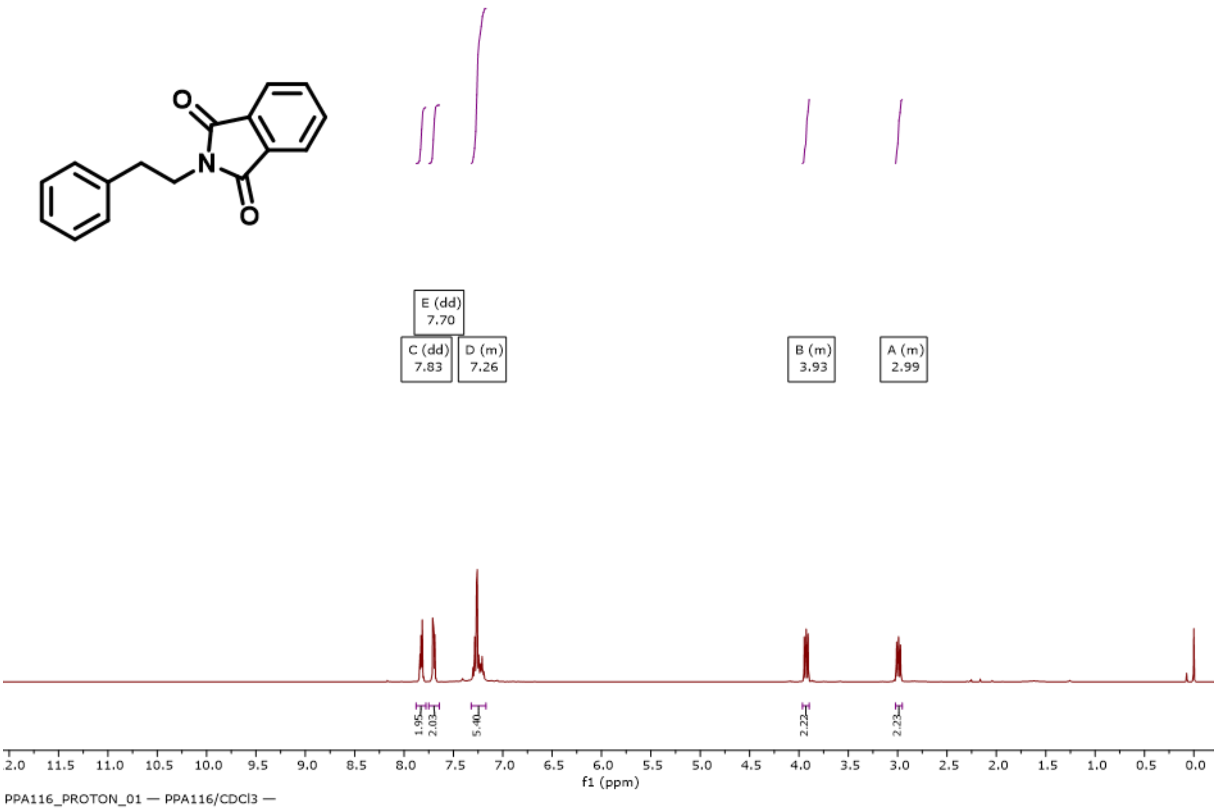

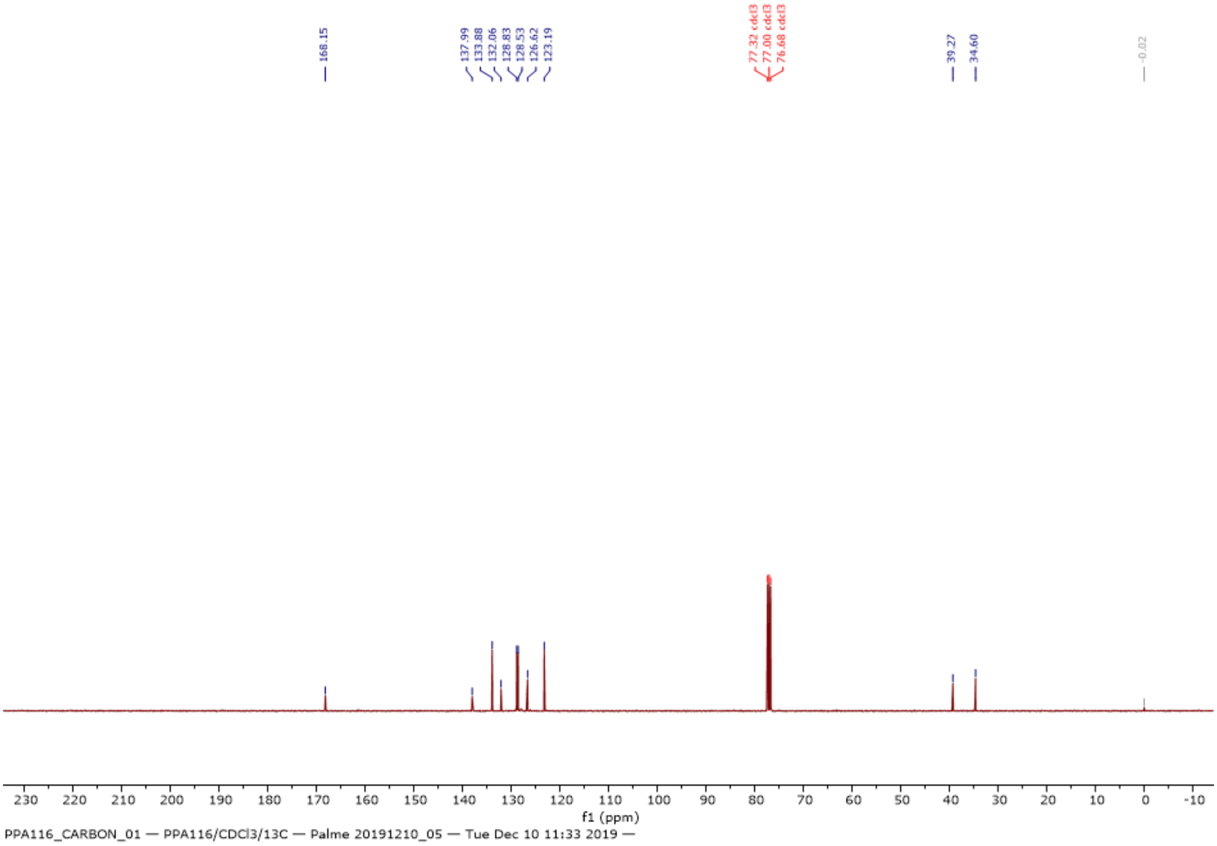

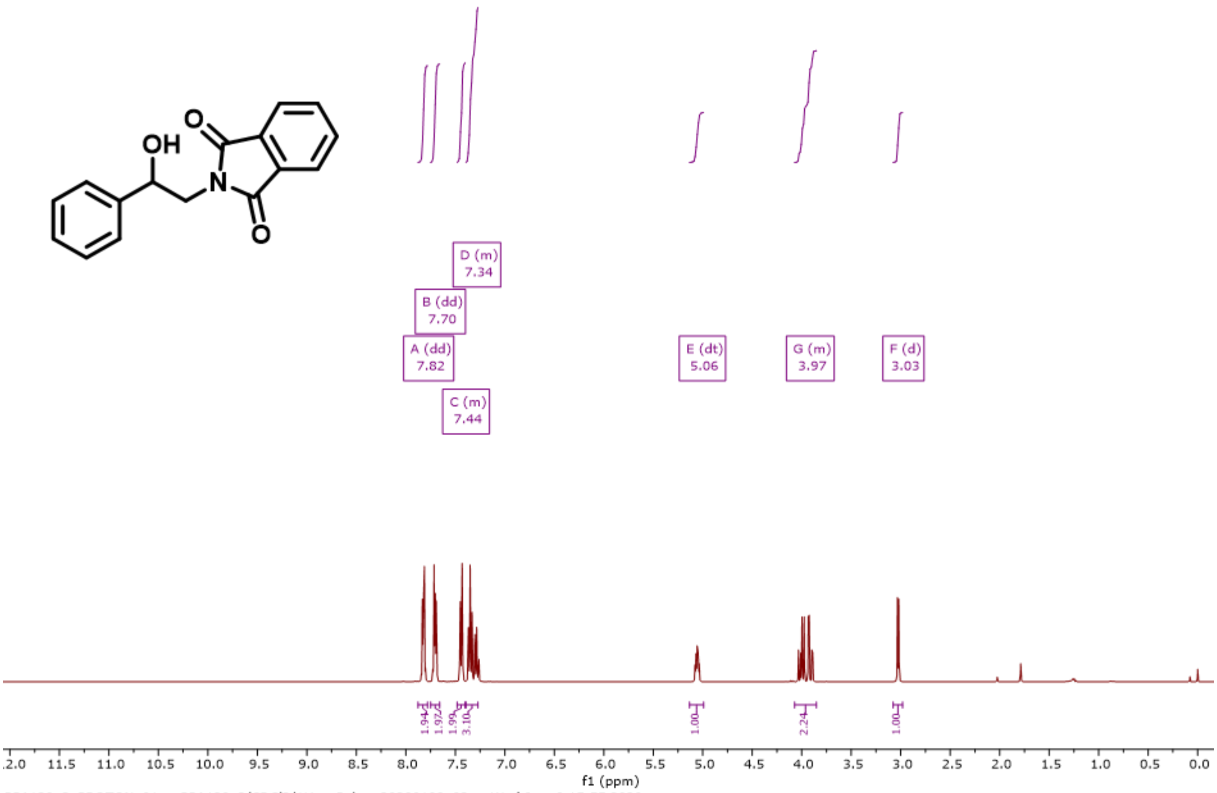

PPA120_2_PROTON_01 — PPA120_2/CDCl3/1H — Palme 20200108_03 — Wed Jan 8 15:55 2020 —

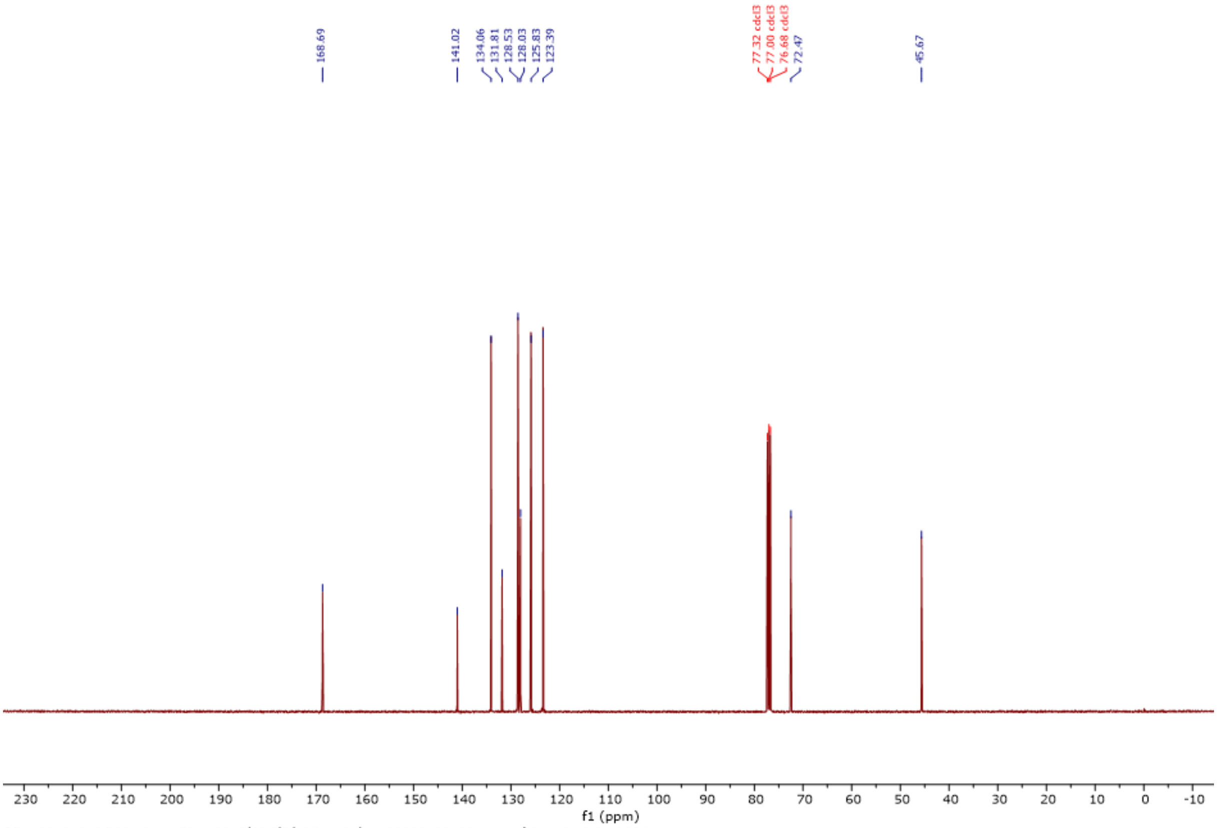

PPA120_2_CARBON_01 — PPA120_2/CDCl3/13C — Palme 20200108_03 — Wed Jan 8 18:01 2020 —

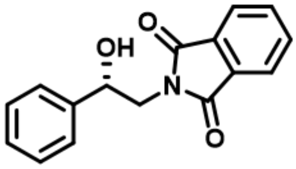

### by chemical conversion

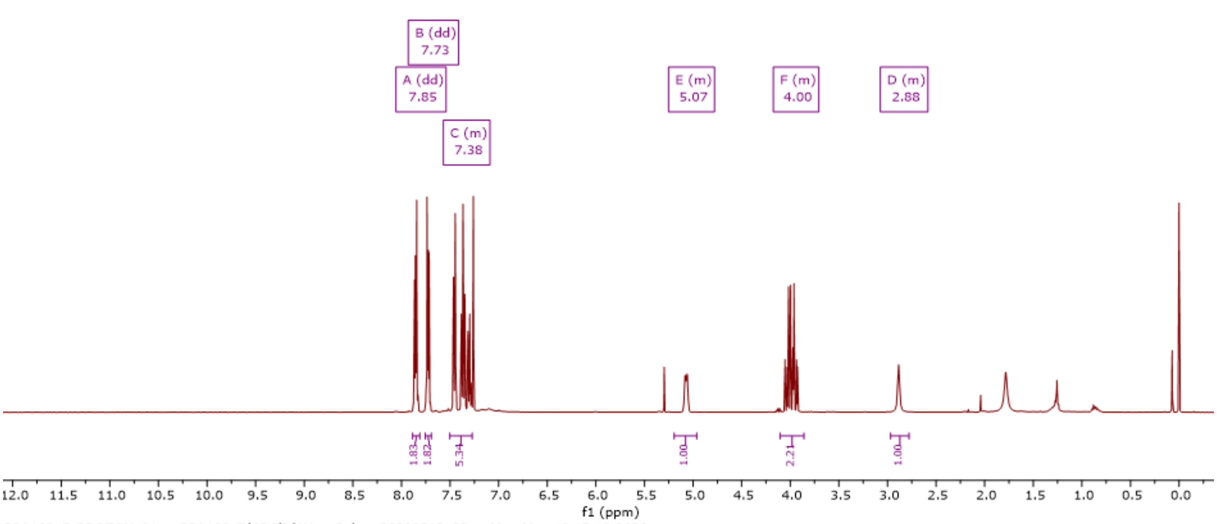

PPA160_F_PROTON_01 — PPA160_F/CDCl3/1H — Palme 20200518_03 — Mon May 18 15:10 2020 —

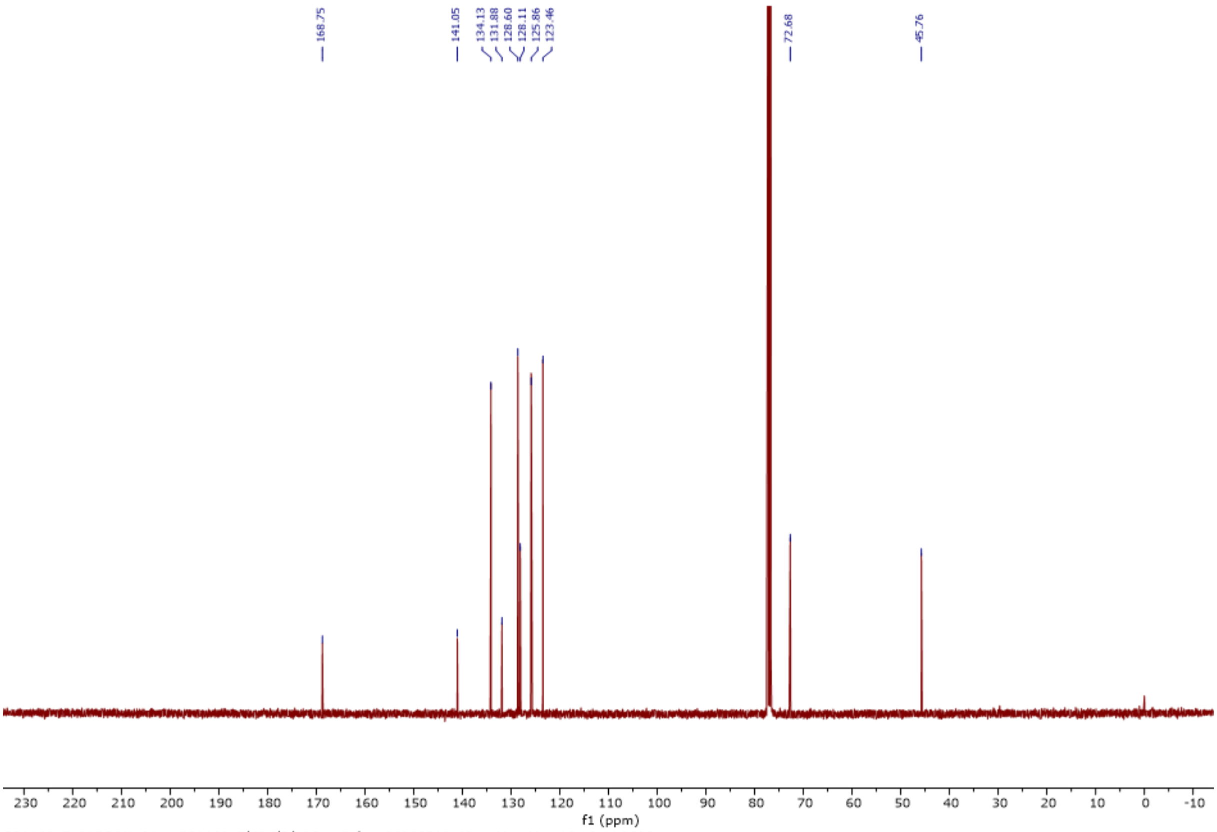

PPA160_F_CARBON_01 — PPA160_F/CDCl3/13C — Palme 20200518_03 — Mon May 18 16:14 2020 —

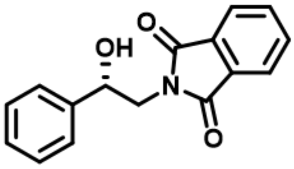

### by enzymatic conversion

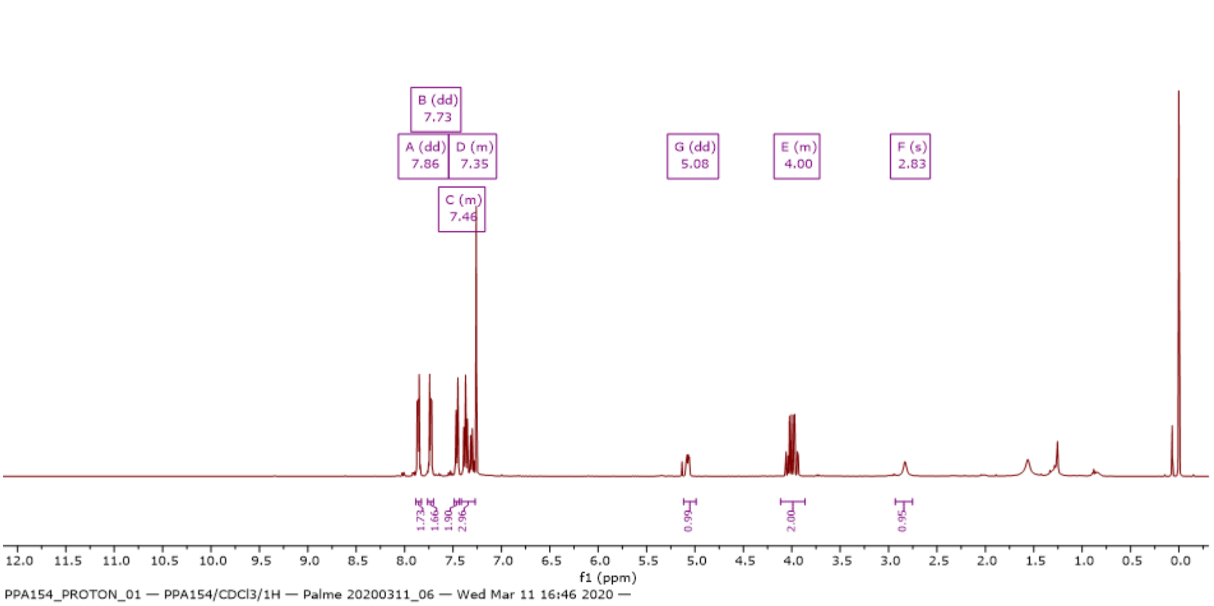

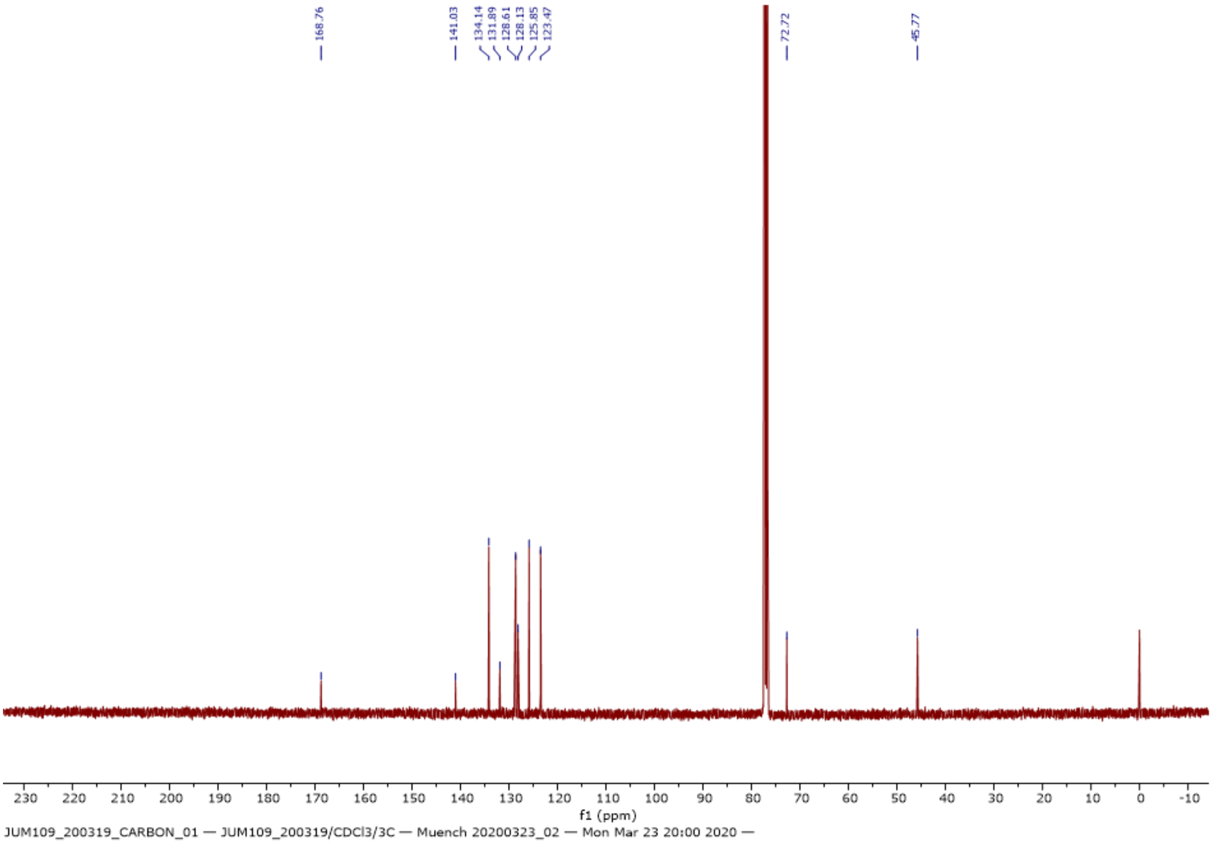

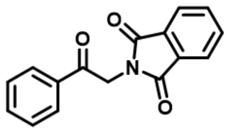

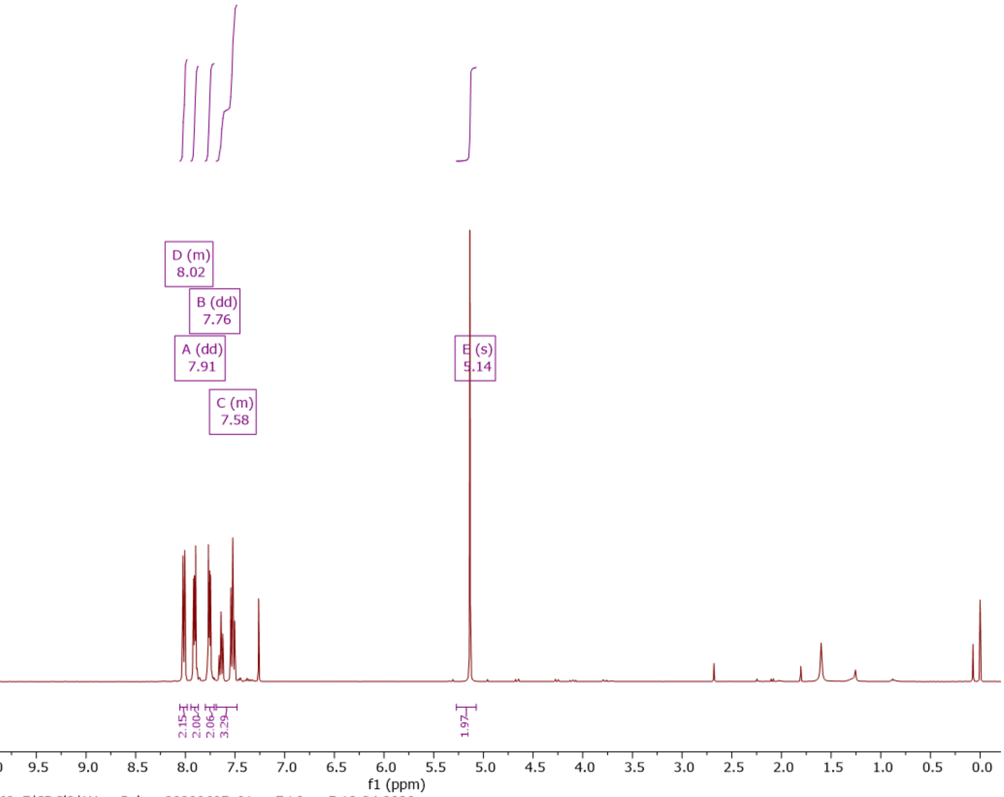

PPA162_F_PROTON_01 — PPA162_F/CDCl3/1H — Palme 20200605_01 — Fri Jun 5 13:34 2020 —

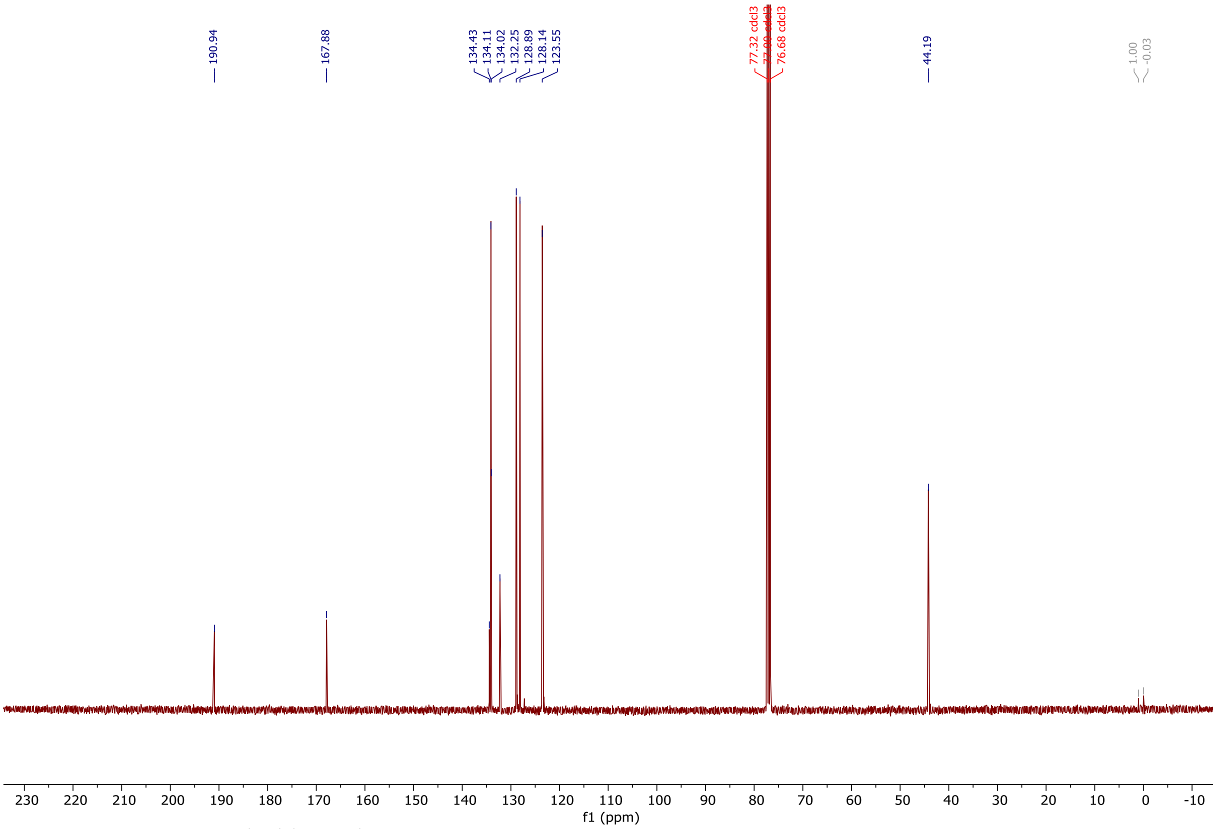

PPA162_F_CARBON_01 — PPA162_F/CDCl3/13C — Palme 20200605_01 — Fri Jun 5 13:58 2020 —
